# Addition of soluble fiber in low fat purified diets improves gut and metabolic health compared to traditional AIN diets

**DOI:** 10.1101/2022.02.28.482306

**Authors:** Laura Griffin, Sridhar Radhakrishnan, Michael Pellizzon

**Affiliations:** Givaudan, 245 Merry Lane, East Hanover, NJ 07936, USA; Research Diets, Inc. 20 Jules Lane, New Brunswick, NJ 08901, USA

**Keywords:** Purified diets, Grain-based diets (chow), Sucrose, Dextrose, Fiber, Fructo-oligosaccharides, Inulin, Cellulose, Glucose tolerance, Gut Microbiota

## Abstract

**Background:** Purified diets (PDs) contain refined ingredients with one main nutrient, allowing for greater control relative to grain-based diets (GBDs), which contain unrefined grains and animal byproducts. Traditional PDs like the AIN-76A (76A) and AIN-93G (93G) can negatively impact metabolic and gut health when fed long-term, in part due to lower total fiber, no soluble fiber, and higher sucrose content.

**Objective:** Two studies were conducted to determine how PDs with reduced sucrose and increased fiber (soluble and insoluble) influence metabolic and gut health in mice compared to traditional AIN PDs or GBDs.

**Methods:** In study 1, C57Bl/6N mice consumed a GBD (5002), 76A, 93G, or 2 Open standard PDs (OSDs) with reduced sucrose and higher fiber for 88 days. Body composition and metabolic parameters were assessed. In study 2, C57Bl/6N mice consumed either 2 GBDs (5001 or 5002) or OSDs with different types/levels of fiber for 14 days. Microbiome alterations and predicted functional metagenomic changes were measured.

**Results:** OSD marginally influenced body weight and adiposity, but improved glucose tolerance relative to 93G (p = 0.0131) and 76A (p = 0.0014). Cecal and colonic weights were lower in mice fed cellulose-based PDs compared to those fed GBDs and soluble fiber PDs. Soluble fiber diets reduced alpha diversity and showed similar beta diversity, which differed from cellulose fed PDs and GBDs. Certain genera associated with improved gut health such as *Bifidobacteria* and *Akkermansia* were significantly elevated by soluble fibers PDs (p≤0.01). Some metabolic pathways related to carbohydrate and fatty acid metabolism were affected by PDs.

**Conclusion:** PDs formulated with lower sucrose and increased fiber content, particularly soluble fiber, blunted elevations in metabolic parameters and favorably impacted microbiota and metagenome in C57BL/6N mice.

## INTRODUCTION

Of the many environmental variables that affect the phenotype of an animal, diet is one that can be easily controlled. Laboratory rodent diets are classified into two main types: grain-based diets (GBDs) or purified diets (PDs). GBDs (or cereal based diets or natural ingredient diets) are typically closed formulas and made with grain-based ingredients and animal byproducts (1). While they provide nutrition for growth and overall health, they contain non-nutritive ingredients such as phytochemicals and potential toxins such as endotoxins, mycotoxins and heavy metals from several ingredients, which may vary from batch-to-batch and potentially influence phenotype (1–3).

PDs are ‘open’ formulas made with defined concentrations of ingredients that are highly refined, each providing one main nutrient (i.e., sucrose is mainly carbohydrate, corn oil is mainly fat and casein is mainly protein). Being highly refined, the non-nutrient content is minimal and the nutrient compositions of both macro- and micronutrients in PDs are well defined, limiting the variability from batch-to-batch (1,4). Since each nutrient/ingredient is added individually, it also allows the researcher to selectively manipulate nutrients, thus providing a wide range of modifications (for e.g. high-fat/protein, high fructose, ketogenic) to study different phenotypes in rodents and in other animal models. The AIN-76A and AIN-93G diets (5,6) are two of the most commonly used PDs, which can provide adequate growth and health of rats and mice; however, there have been reports (7,8) of mild metabolic dysfunction (increased body weight, body fat, mild insulin resistance, hyperlipidemia, etc.) in animals consuming these diets, relative to GBD fed animals. While several differences exist between these two types of diets, these perturbations may be in part due to certain ingredients in these diets, including their higher sucrose content (10% and 50% w/w in 93G and 76A, respectively) and a low amount of total and mostly non-fermentable fiber (5% cellulose)(9). This is in stark contrast to the presence of minimal amounts of sucrose and relatively higher amounts (15-25% w/w) of fiber in GBDs. In addition, GBDs also contain diverse sources of fiber including soluble (beta-glucan, pectin, etc.), partially soluble (hemicellulose) and insoluble fibers (celluloses, lignin, etc.) (1,2).

Metabolic differences that are observed in rodents consuming AIN PDs relative to those fed GBDs may be driven in part by differences in sucrose and fiber, in particular, soluble fiber. Modifications to these PDs include replacement of sucrose with sources such as corn starch and dextrose to minimize fructose, an initiator of metabolic disease, including insulin resistance, glucose intolerance, and hyperlipidemia(11,12). Even relatively low levels of sucrose may influence glucose tolerance over more chronic feeding periods(13). The fiber content of PDs can also be increased and furthermore, refined soluble fiber sources, such as fructo-oligosaccharides (FOS) or inulin, which have been long known to promote metabolic health via the gut (14–16), can be added to these diets. We and others have shown that the addition of soluble fibers such as inulin, in the context of a diet higher in fat can reduce body weight, adiposity, and blood and liver lipids, inflammation, and improve glucose tolerance of rodents, perhaps in part, due to elevated short-chain fatty acid production by the gut bacteria or other factors (17–19).

The purpose of this study, therefore, was to assess whether changes to the carbohydrate and fiber components of the AIN PDs could improve metabolic health of rodents. Furthermore, we determined how changes in the type and amount of fiber affected gut health/microbiome profile. Metabolic and gut microbiome effects in mice fed PDs were also compared to those fed GBDs.

## METHODS

### Dietary Formulations

#### Study 1

In addition to the traditional AIN PDs, AIN-76A (76A) and AIN-93G (93G), we used two modified versions of the AIN diets, referred to as the Open Standard Diets (OSDs). The nutritional profiles of the four PDs utilized are presented in **Table 1**. The OSDs contained only trace levels of sucrose (in the vitamin and mineral mixes), providing around 1% of total kcals. The diet D11112201 (OSD) was formulated with 100 gm of added fiber per 4084 kcals in a 3:1 ratio of cellulose to inulin (75 g cellulose and 25 g inulin per 4084 kcals, 9.3% fiber w/w), with inulin providing approximately 1.5 kcal/g from fermentation (20). D11112202 (OSD + F) (20.5% total fiber w/w) contained three times as much cellulose, (225 g per 4084 kcals) as D11112201, but the same amount of inulin (25 g per 4084 kcals), to be more in line with GBDs which contain higher amounts of fiber as insoluble fiber with some soluble fiber. All PDs were formulated by Research Diets, Inc. (New Brunswick, NJ, USA). These diets were compared to the GBD, LabDiet 5002 (chow) (St. Louis, MO). Fiber content of this chow was analyzed by the Laboratory of Dr. Kelly Swanson, Dept. of Animal Sciences, University of Illinois. The total fiber content was 23.9%. Insoluble fiber content was 18.6% and soluble fiber content was 5.3%.

**Table 1:** Composition of the PDs used for Study 1.

| Table 1: Composition of the PDs used for Study 1 |  |  |  |  |  |  |  |  |
| --- | --- | --- | --- | --- | --- | --- | --- | --- |
| Product # | D10001 |  | D10012G |  | D11112201 |  | D11112202 |  |
|  | AIN-76A |  | AIN-93G |  | OSD |  | OSD-F |  |
|  | gm% | kcal% | gm% | kcal% | gm% | kcal% | gm% | kcal% |
| Protein | 20 | 21 | 20 | 20 | 19 | 20 | 17 | 20 |
| Carbohydrate | 66 | 68 | 64 | 64 | 63 | 65 | 42 | 65 |
| Fat | 5 | 12 | 7 | 16 | 7 | 15 | 6 | 15 |
| Total |  | 100 |  | 100 |  | 100 |  | 100 |
| kcal/gm | 3.90 |  | 4.00 |  | 3.81 |  | 3.34 |  |
| Ingredient | gm | kcal | gm | kcal | gm | kcal | gm | kcal |
| Casein | 200 | 800 | 200 | 800 | 200 | 800 | 200 | 800 |
| DL-Methionine | 3 | 12 | 0 | 0 | 0 | 0 | 0 | 0 |
| L-Cystine | 0 | 0 | 3 | 12 | 3 | 12 | 3 | 12 |
| Corn Starch | 150 | 600 | 397.486 | 1590 | 381 | 1524 | 381 | 1524 |
| Maltodextrin | 0 | 0 | 132 | 528 | 110 | 440 | 110 | 440 |
| Sucrose | 500 | 2000 | 100 | 400 | 0 | 0 | 0 | 0 |
| Dextrose, Monohydrate | 0 | 0 | 0 | 0 | 150 | 600 | 150 | 600 |
| Cellulose (Insoluble Fiber) | 50 | 0 | 50 | 0 | 75 | 0 | 225 | 0 |
| Inulin (Soluble Fiber) | 0 | 0 | 0 | 0 | 25 | 38 | 25 | 38 |
| Corn Oil | 50 | 450 | 0 | 0 | 0 | 0 | 0 | 0 |
| Soybean Oil | 0 | 0 | 70 | 630 | 70 | 630 | 70 | 630 |
| t-BHQ | 0 | 0 | 0.014 | 0 | 0 | 0 | 0 | 0 |
| Mineral Mix S10001 | 35 | 0 | 0 | 0 | 0 | 0 | 0 | 0 |
| Mineral Mix S10022G | 0 | 0 | 35 | 0 | 0 | 0 | 0 | 0 |
| Mineral Mix S10026 | 0 | 0 | 0 | 0 | 10 | 0 | 10 | 0 |
| Dicalcium Phosphate | 0 | 0 | 0 | 0 | 13 | 0 | 13 | 0 |
| Calcium Carbonate | 0 | 0 | 0 | 0 | 5.5 | 0 | 5.5 | 0 |
| Potassium Citrate, 1 H2O | 0 | 0 | 0 | 0 | 16.5 | 0 | 16.5 | 0 |
| Vitamin Mix V10001, AIN-76A | 10 | 40 | 0 | 0 | 10 | 40 | 10 | 40 |
| Vitamin Mix V10037, AIN-93G | 0 | 0 | 10 | 40 | 0 | 0 | 0 | 0 |
| Choline Bitartrate | 2 | 0 | 2.5 | 0 | 2 | 0 | 2 | 0 |
| Yellow Dye #5, FD&C | 0 | 0 | 0 | 0 | 0.025 | 0 | 0 | 0 |
| Red Dye#40, FD&C | 0 | 0 | 0 | 0 | 0 | 0 | 0.05 | 0 |
| Blue Dye #1, FD&C | 0 | 0 | 0 | 0 | 0.025 | 0 | 0 | 0 |
| **Total** | **1000** | **3902** | **1000** | **4000** | **1071.05** | **4084** | **1221.05** | **4084** |

#### Study 2

To further understand the role of fiber type and concentration in PDs and how they compare to GBDs, we used six additional versions of the OSD with either 100 or 200 g of cellulose (CEL), inulin (IN), or fructo-oligosaccharides (FOS) per 4084 kcals. The six experimental OSDs (100CEL, 200CEL, 100IN, 200IN, 100FOS, and 200FOS) were compared to two GBDs, LabDiet 5001 and 5002 (LabDiet, St. Louis, MO), see **Table 2**. Fiber content of 5001 and 5002 were analyzed by Covance Laboratories (Madison, WI); 5001: 18.7% total, 15.9% insoluble, 2.8% soluble, 5002: 18.2% total, 14.9% insoluble fiber, and 3.3% soluble fiber.

**Table 2:** Composition of PDs used in Study 2.

| Table 2: Composition of PDs used in Study 2 |  |  |  |  |  |  |  |  |  |  |  |  |
| --- | --- | --- | --- | --- | --- | --- | --- | --- | --- | --- | --- | --- |
| Product # | D11112222 |  | D11112223 |  | D11112224 |  | D11112225 |  | D11112226 |  | D11112227 |  |
|  | OSD+200 g Cellulose |  | OSD+200 g Inulin |  | OSD+200 g FOS |  | OSD + 100 g Cellulose |  | OSD + 100 g Inulin |  | OSD + 100 g FOS |  |
|  | 200CEL |  | 200IN |  | 200FOS |  | 100CEL |  | 100IN |  | 100FOS |  |
|  | gm% | kcal% | gm% | kcal% | gm% | kcal% | gm% | kcal% | gm% | kcal% | gm% | kcal% |
| Protein | 17.2 | 20 | 18.4 | 20 | 18.4 | 20 | 18.8 | 20 | 19.5 | 20 | 19.5 | 20 |
| Carbohydrate | 55.9 | 65 | 71.1 | 65 | 53.0 | 57 | 61.1 | 65 | 69.3 | 65 | 59.7 | 61 |
| Fat | 5.9 | 15 | 6.3 | 15 | 6.3 | 15 | 6.5 | 15 | 6.7 | 15 | 6.7 | 15 |
| Total |  | 100 |  | 100 |  | 100 |  | 100 |  | 100 |  | 100 |
| kcal/gm | 3.46 |  | 3.69 |  | 3.69 |  | 3.78 |  | 3.92 |  | 3.92 |  |
| Ingredient | gm | kcal | gm | kcal | gm | kcal | gm | kcal | gm | kcal | gm | kcal |
| Casein | 200 | 800 | 200 | 800 | 200 | 800 | 200 | 800 | 200 | 800 | 200 | 800 |
| L-Cystine | 3 | 12 | 3 | 12 | 3 | 12 | 3 | 12 | 3 | 12 | 3 | 12 |
| Corn Starch | 390.5 | 1562 | 315.5 | 1262 | 315.5 | 1262 | 390.5 | 1562 | 353 | 1412 | 353 | 1412 |
| Maltodextrin 10 | 110 | 440 | 110 | 440 | 110 | 440 | 110 | 440 | 110 | 440 | 110 | 440 |
| Dextrose | 150 | 600 | 150 | 600 | 150 | 600 | 150 | 600 | 150 | 600 | 150 | 600 |
| Cellulose | 200 | 0 | 0 | 0 | 0 | 0 | 100 | 0 | 0 | 0 | 0 | 0 |
| Inulin | 0 | 0 | 200 | 300 | 0 | 0 | 0 | 0 | 100 | 150 | 0 | 0 |
| Fructooligosaccharide | 0 | 0 | 0 | 0 | 200 | 300 | 0 | 0 | 0 | 0 | 100 | 150 |
| Soybean Oil | 70 | 630 | 70 | 630 | 70 | 630 | 70 | 630 | 70 | 630 | 70 | 630 |
| Mineral Mix S10026 | 10 | 0 | 10 | 0 | 10 | 0 | 10 | 0 | 10 | 0 | 10 | 0 |
| Dicalcium Phosphate | 13 | 0 | 13 | 0 | 13 | 0 | 13 | 0 | 13 | 0 | 13 | 0 |
| Calcium Carbonate | 5.5 | 0 | 5.5 | 0 | 5.5 | 0 | 5.5 | 0 | 5.5 | 0 | 5.5 | 0 |
| Potassium Citrate, 1 H2O | 16.5 | 0 | 16.5 | 0 | 16.5 | 0 | 16.5 | 0 | 16.5 | 0 | 16.5 | 0 |
| Vitamin Mix V10001 | 10 | 40 | 10 | 40 | 10 | 40 | 10 | 40 | 10 | 40 | 10 | 40 |
| Choline Bitartrate | 2 | 0 | 2 | 0 | 2 | 0 | 2 | 0 | 2 | 0 | 2 | 0 |
| Yellow Dye #5, FD&C | 0.05 | 0 | 0 | 0 | 0 | 0 | 0.025 | 0 | 0 | 0 | 0.01 | 0 |
| Red Dye #40, FD&C | 0 | 0 | 0.05 | 0 | 0 | 0 | 0.025 | 0 | 0.025 | 0 | 0.04 | 0 |
| Blue Dye #1, FD&C | 0 | 0 | 0 | 0 | 0.05 | 0 | 0 | 0 | 0.025 | 0 | 0 | 0 |
| Total | 1180.55 | 4084 | 1105.55 | 4084 | 1105.55 | 4084 | 1080.55 | 4084 | 1043.05 | 4084 | 1043.05 | 4084 |
| Total Fiber (%) | 16.9 |  | 18.1 |  | 18.1 |  | 9.3 |  | 9.6 |  | 9.6 |  |
| Total Insoluble (%) | 16.9 |  | 0 |  | 0 |  | 9.3 |  | 0 |  | 0 |  |
| Total Soluble (%) | 0 |  | 18.1 |  | 18.1 |  | 0 |  | 9.6 |  | 9.6 |  |

### Animals and Study Design

#### Study 1

This study was conducted at MuriGenics, Inc. (Vallejo, CA, USA). Weanling male C57Bl/6N mice (n = 75) were purchased from Charles River Laboratories. Mice were housed (*n* = 5/cage on Alpha-Dri bedding) in micro isolators on a 12hr light/dark cycle and maintained on chow (LabDiet 5001). Food and water were provided *ad libitum*. At 4 weeks of age, mouse cages were randomly assigned to one of the five treatment groups (3 cages/treatment; *n* = 15 animals/treatment). Mice were maintained on the experimental diets for 88 days. Food and water intake per cage and body weights were measured weekly throughout the study.

#### Study 2

The study was conducted at Charles River Laboratories (Wilmington, MA, USA). Male weanling C57Bl/6N mice (n = 54) from Charles River Laboratories were housed 3 per cage on Alpha-Dri bedding. Mice were fed 5001 *ad libitum* and housed in standard vivarium conditions on a 12-hr light/dark cycle. After 2 days following receipt, six mice were sacrificed for baseline data. At 4 weeks of age, initial body weight measurements were collected and mouse cages were randomly assigned to one of 8 treatment groups (2 cages/treatment, n = 6/treatment). Animals were maintained on the experimental diets for 14 days. Body weights were measured prior to euthanasia.

### Glucose Tolerance Test

#### Study 1

On day 83, an oral glucose tolerance test (OGTT) was performed after 12 weeks on the experimental diets with modification, as described previously (21). Briefly, following a 6-hour fast, baseline blood glucose measurements were taken. Immediately afterwards, mice were gavaged a 20% dextrose solution to deliver 2 g glucose/kg body weight. Blood glucose measurements were collected via the tail vein at 30 minute intervals over the course of 2 hours. Food was restored upon completion of the procedure.

### Euthanasia and Necropsy

#### Study 1

On day 88, mice were fasted for 6 hours, blood samples were collected via cardiac puncture, and euthanized. Serum was separated and stored at -80C until analysis. Following blood collection, mice were perfused with saline. Carcass, liver, and select adipose deposits (mesenteric, gonadal, inguinal, retroperitoneal) were weighed. Livers were flash frozen in liquid nitrogen and stored at -80C for triglyceride analysis.

#### Study 2

At the end of 14 days, animals were euthanized using CO2. Following CO2 asphyxiation, the cecum and colon were harvested together with their contents remaining intact. The total tissue was weighed. For the first animal in each group, an image of the colon and cecum (attached) was taken next to a standard ruler. The colon and cecum were separated from each other (contents still intact) and each were weighed. Colon and cecum contents were collected into separate vials and placed on dry ice before being stored at -70°C. The colon and cecum were cleaned with DD water, blotted dry, and weighed individually. Colonic length was recorded once the tissue was cleaned and dry.

### Serum and Triglyceride Analysis

#### Study 1

Serum from each animal was analyzed for fasting blood glucose, leptin, triglycerides, and total cholesterol at MuriGenics (Vallejo, CA, USA). Insulin and leptin were analyzed using a multiplex assay. Liver samples were assayed for triglyceride content by Vascular Strategies LLC (Wynnewood, PA, USA).

### Microbiome Analysis

#### Sequencing QC and Analysis

16S rRNA sequencing was performed on the cecum and colon content samples at the Argonne National Laboratory (Argonne, IL, USA). Briefly, total DNA was extracted from the samples, and the V3–V4 regions of the 16S rRNA were amplified using PCR and sequenced using their MG-RAST pipeline. Sequence data from this pipeline was transferred to Diversigen (Minneapolis, MN, USA) for subsequent analysis. Raw FASTQs for all samples were run through the QC pipeline at Diversigen. Briefly, this involves trimming of adapter sequences (if present) using the program *cutadapt* followed by filtering of reads to remove any with a mean Q-score of < 30. Next, FASTQ files were run through their standard *dada2* pipeline to produce the raw Amplicon Sequence Variant (ASV) count table. The raw table was then filtered to remove any ASVs at < 0.0001 % sum relative abundance across all samples. Finally, we used the filtered ASV table as input to *PICRUSt2* infer functional metagenomic content of all samples. The outputs of this were an abundance table of predicted Enzyme 1 Commission (EC) numbers, as well as an abundance table of predicted Metacyc Pathways derived from the EC abundance data.

#### Taxa and Functional Summary Plots

The filtered ASV table was used to generate summary stacked bar plots of relative abundances per-sample at the phylum and ASV levels for both cecums and colons. Comparisons between phyla and genera by diet and tissue type can be found in the supplemental tables. For visualization purposes, only the top 20 most abundant genera are plotted in the Genus-level plot, with the rest allocated to an “Other” category. At the Phylum dlevel, the ratio of Firmicutes/Bacteroidota was calculated per-sample by dividing the relative abundances for each of these. The results of these ratios were then plotted across all diet groups. To assess whether the Firmicutes/Bacteroidota ratio differed across diet groups, we used a one-way ANOVA across treatment groups followed by Tukey’s HSD. To examine broad patterns of predicted functional genomic content across diet groups, the predicted EC count table was used as an input. EC numbers in this table were then grouped according to six broad functional categories as defined in the Kyoto Encyclopedia of Genes and Genomes (KEGG) database: (1) Carbohydrate Degradation and Absorption, (2) Fructose and Mannose Metabolism, (3) Galactose Metabolism, (4) Starch and Sucrose Metabolism, (5) Fatty Acid Metabolism and (6) Fatty Acid Biosynthesis. Counts for ECs found within each of these categories were then aggregated per-category, and the resulting count data was plotted as boxplots. To test whether counts differed across diet groups within these categories, we used a one-way ANOVA across all treatment groups followed by Tukey’s HSD.

#### Alpha and Beta Diversity Analyses

Using the ASV, EC and Pathways tables, we rarefied each table to the sample with the lowest mapped counts. We then calculated three alpha diversity metrics for each of the three feature tables: the Shannon index, the Chao1 index and Observed Features. To examine all pairwise comparisons of each treatment group to every other group, we used a one-way ANOVA across all treatment groups followed by Tukey’s HSD. All results were then plotted as box plots with strip charts overlaid to show all data points. To assess differences in between-sample (i.e. beta) diversity, we calculated distance matrices for all three rarefied feature tables using the Bray-Curtis Dissimilarity metric. Next, we used Permutational Multivariate Analysis of Variance (PERMANOVA) from the R package *vegan* to assess differences in beta diversity between treatment groups. The results were then plotted using principal coordinates analysis (PCoA) plots with samples colored by treatment group. We also determined the beta diversity variability within each treatment group by calculating the distance-to-centroid for every sample within its group. We then used the same statistical methodologies used for alpha diversity to determine if beta diversity variability is different between any treatment groups. All results of statistical testing are included with individual plots.

#### Differential Abundance Analysis

Finally, we assessed whether any taxa – Phylum or Genus level or functions (Metacyc Pathways-level) differ in abundance between the treatment groups in the study. Due to the compositional nature of microbiome abundance data, we utilized the R package *ALDEx2*. Briefly, this package performs differential abundance testing of count data across samples between two experimental groups by starting with zero-estimation for any features with zero abundance in some samples but not others. Next, per-feature technical variation for each feature is estimated for each sample using Monte-Carlo sampling from a Dirichlet distribution. Each instance of Monte-Carlo sampling is then transformed using the Centered LogRatio (CLR) transformation, at which point pairwise statistical testing is performed between experimental groups using the CLR-transformed abundance values. This process is repeated for each instance, and results of statistical testing are aggregated yielding adjusted p-values (Benjamini-Hochberg corrected) for each feature in the differential abundance test of interest. We used the above procedure to test for differential abundance between every pairwise comparison of treatment groups. Individual p-value tables of the results of statistical testing for every feature in a given pairwise comparison is presented as supplementary files. For all statistical tests, an adjusted p-value of < 0.05 was used as a threshold for significance.

### Statistical Analysis

For study 1, all data were analyzed by 1-way ANOVA with GraphPad Prism. Post-hoc analyses were performed if p < 0.05 using Tukey’s HSD for comparisons among groups (both studies). For study 2, to examine all pairwise comparisons between treatment groups, 2-way ANOVA followed by Tukey’s HSD was employed using R. Microbial statistics are presented in the section above. An adjusted p-value of p < 0.05 was used as a threshold for significance for all statistical tests.

## RESULTS

### *STUDY 1:* Body Composition and Metabolic Assessment

Over time, the average body weights of OSD and 93G treatment groups were slightly higher (by mean of 2.3 g) compared with those fed the OSD+F, 76A, and 5002 diets (**Figure 1**). Terminal body weights were similar for all groups, though OSD was statistically higher than 76A (p = 0.0102), but similar to 5002 (p = 0.163) and 93G (p =0.8223) and additional fiber as cellulose in the OSD+F group blunted this effect. All the individual fat pad (mesenteric, gonadal, retroperitoneal and inguinal) weights were generally similar among groups. OSD fed mice had significantly heavier gonadal fat pads (p = 0.0312) and total fat (p = 0.0479) compared to 5002 and significantly heavier inguinal (subcutaneous) fat pads (p = 0.0036) compared to 93G **(Table 3)**; this difference was blunted by the addition of extra cellulose in the OSD+F group. The carcass weights were generally similar across groups, although in the OSD group, the carcass weight was significantly higher than in the 76A group (p = 0.019). Adiposity index (g total fat / g carcass) was also similar among groups though it tended to be higher in OSD than 5002 (p = 0.0548), but significantly greater in OSD vs. 93G (p = 0.0372) due mainly to a lower inguinal fat pad weight in the latter group. The addition of cellulose in OSD+F blunted the adiposity index and resulted in similar levels relative to all groups **(Table 3)**. The liver weights were also not significantly different among the five groups (data not shown).

**Table 3.**
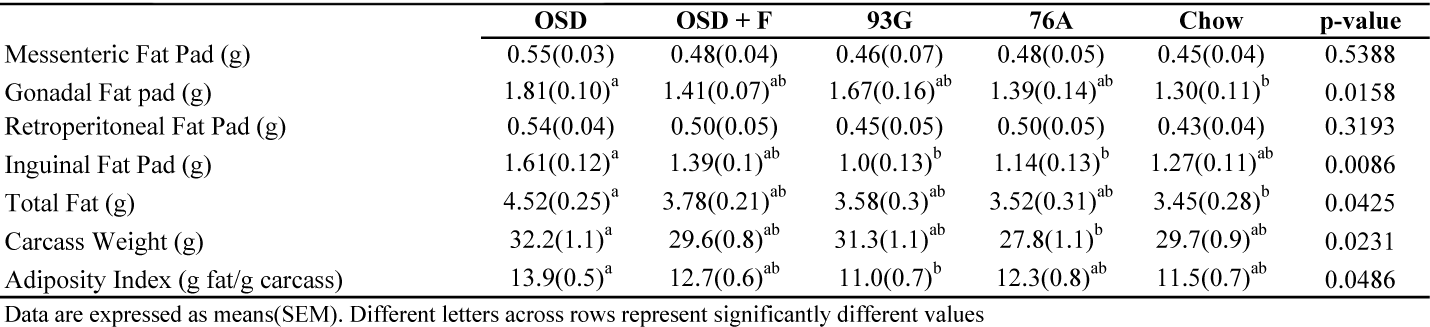
Fat Pad and Carcass Weights in Study 1

|  | OSD | OSD + F | 93G | 76A | Chow | p-value |
| --- | --- | --- | --- | --- | --- | --- |
| Mesenteric Fat Pad (g) | 0.55(0.03) | 0.48(0.04) | 0.46(0.07) | 0.48(0.05) | 0.45(0.04) | 0.5388 |
| Gonadal Fat pad (g) | 1.81(0.10)<sup>a</sup> | 1.41(0.07)<sup>ab</sup> | 1.67(0.16)<sup>ab</sup> | 1.39(0.14)<sup>ab</sup> | 1.30(0.11)<sup>b</sup> | 0.0158 |
| Retroperitoneal Fat Pad (g) | 0.54(0.04) | 0.50(0.05) | 0.45(0.05) | 0.50(0.05) | 0.43(0.04) | 0.3193 |
| Inguinal Fat Pad (g) | 1.61(0.12)<sup>a</sup> | 1.39(0.1)<sup>ab</sup> | 1.0(0.13)<sup>b</sup> | 1.14(0.13)<sup>b</sup> | 1.27(0.11)<sup>ab</sup> | 0.0086 |
| Total Fat (g) | 4.52(0.25)<sup>a</sup> | 3.78(0.21)<sup>ab</sup> | 3.58(0.3)<sup>ab</sup> | 3.52(0.31)<sup>ab</sup> | 3.45(0.28)<sup>b</sup> | 0.0425 |
| Carcass Weight (g) | 32.2(1.1)<sup>a</sup> | 29.6(0.8)<sup>ab</sup> | 31.3(1.1)<sup>ab</sup> | 27.8(1.1)<sup>b</sup> | 29.7(0.9)<sup>ab</sup> | 0.0231 |
| Adiposity Index (g fat/g carcass) | 13.9(0.5)<sup>a</sup> | 12.7(0.6)<sup>ab</sup> | 11.0(0.7)<sup>b</sup> | 12.3(0.8)<sup>ab</sup> | 11.5(0.7)<sup>ab</sup> | 0.0486 |

Data are expressed as means(SEM). Different letters across rows represent significantly different values

**Figure 1.**
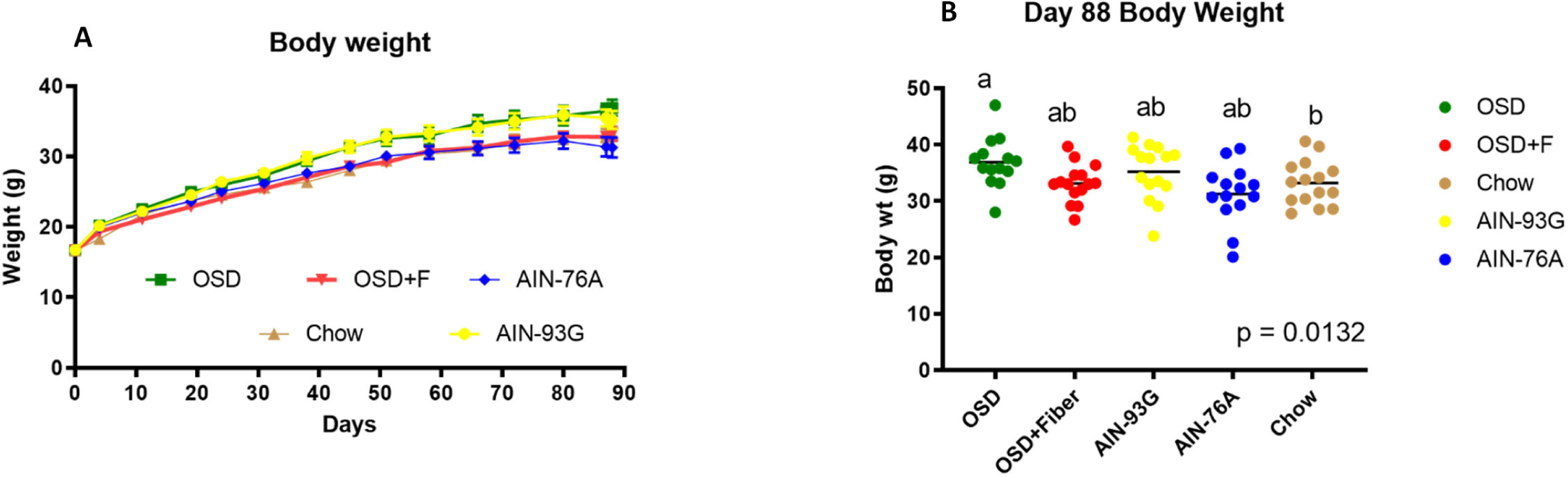
(Study 1) A. Mouse body weight measurements over an 88 day timespan expressed as means ± SEM and B. Final body weight measurements expressed as means ± SEM. Groups with different letters represent significantly different results by 1-way ANOVA with Tukey’s HSD post hoc analysis (p < 0.05). Mouse weights were recorded weekly over the 88 day metabolic phenotype study for each of the 5 dietary treatment groups (n = 15/treatment): OSD, OSD + F, 76A, 93G, and chow.

#### Serum Biochemistry

Glucose homeostasis was assessed prior to study termination with an oral glucose tolerance test (**Figure 2**). All groups had similar 6-hr fasting blood glucose levels (**Figure 2A**). Glucose tolerance was significantly reduced in 93G and 76A mice (as shown by a greater area under the curve) compared to both chow (vs. 93G, p = 0.0399, vs. 76A, p = 0.005) and OSD mice (vs. 93G, p = 0.0131; vs. 76A, p = 0.0014) (**Figure 2B**). OSD+F was intermediate and similar to all groups. Additional measurements of 6-hr fasting serum glucose and insulin measurements were made at study termination (**Table 4**). Serum cholesterol was significantly higher in all PD groups compared to chow (OSD, p=0.0037; OSD+Fiber, p=0.0005; 76A, p=0.001) and was highest in the 93G group (p < 0.0001) (**Table 4**). Serum triglycerides were similar among groups, but were significantly lower for 76A compared to chow (p = 0.0009). Serum leptin was higher in OSD and 93G fed mice compared to those fed OSD+F (OSD, p=0.0207; 93G, p=0.0057) and chow (OSD, p=0.007; 93G, p=0.0017), while 76A had a similar leptin level relative to all groups. Liver triglycerides were not different between chow, OSD+F and 76A fed mice, but OSD and 93G had higher levels compared to chow (OSD, p=0.0008; 93G, p=0.0002) and 93G had higher levels than 76A (p=0.0328).

**Table 4.** Terminal Serum and Liver Data in Study 1

|  | OSD | OSD + F | 93G | 76A | Chow | p-value |
| --- | --- | --- | --- | --- | --- | --- |
| Serum Insulin (ng/mL) | 4.6(0.6) | 3.4(0.4) | 5.2(0.6) | 3.3(0.6) | 3.5(0.5) | 0.0378 |
| Serum Glucose (mg/dL) | 267.8(16.4) | 244.1(12.9) | 254.7(8.9) | 251.0(15.9) | 236.0(11.3) | 0.5248 |
| Serum Triglyceride (mg/dL) | 96.5(4.4)<sup>ab</sup> | 92.2(2.7)<sup>ab</sup> | 92.7(3.2)<sup>ab</sup> | 80.9(0.05)<sup>b</sup> | 102.7(3.2)<sup>a</sup> | 0.0031 |
| Serum Cholesterol (mg/dL) | 166.7(10.2)<sup>a</sup> | 171.9(5.3)<sup>a</sup> | 223.5(6.2)<sup>c</sup> | 170.6(9.0)<sup>a</sup> | 129.7(2.6)<sup>b</sup> | <0.0001 |
| Serum Leptin (ng/mL) | 34.3(1.2)<sup>a</sup> | 23.5(2.6)<sup>b</sup> | 35.6(2.4)<sup>a</sup> | 26.2(3.0)<sup>ab</sup> | 22.2(2.6)<sup>b</sup> | 0.0002 |
| Liver Weight (g) | 1.55(0.10) | 1.39(0.08) | 1.36(0.08) | 1.36(0.06) | 1.57(0.04) | 0.1096 |
| Liver Triglyceride (mg/g) | 10.1(1.0)<sup>ab</sup> | 8.2(0.9)<sup>abc</sup> | 10.5(1.3)<sup>a</sup> | 6.5(1.0)<sup>bc</sup> | 4.5(0.3)<sup>c</sup> | <0.0001 |

Data are expressed as means(SEM). Different letters across rows represent significantly different values

**Figure 2:**
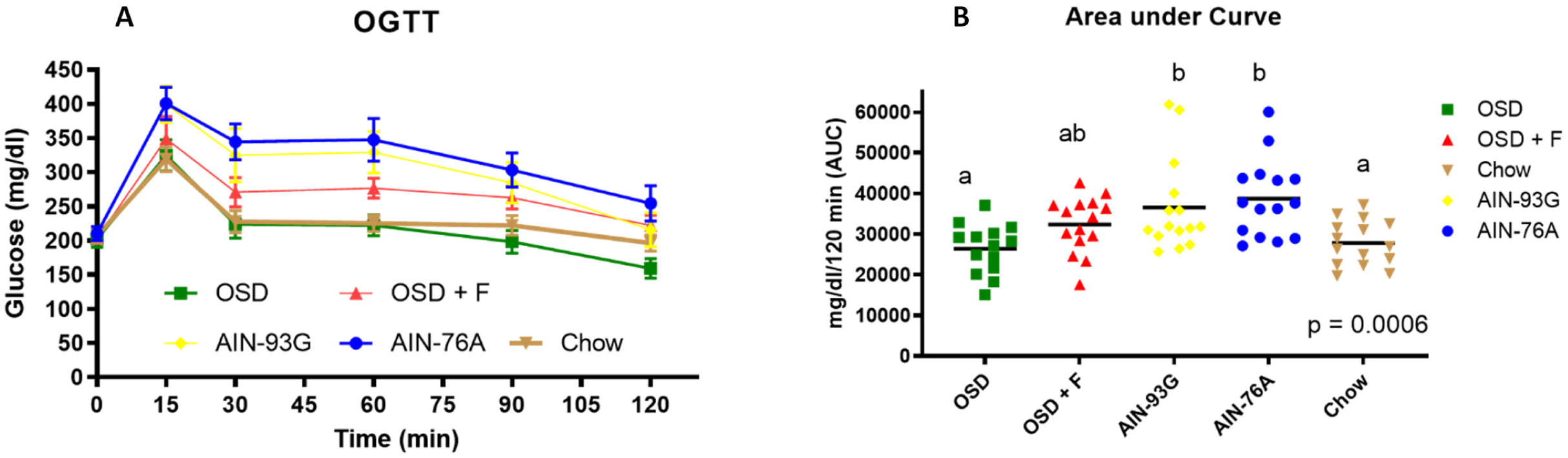
(Study 1). A. Blood glucose measurements over time following a 6-hr fast and an oral glucose load, with data points representing means ± SEM for each treatment group. B. Blood glucose area under the curve on day 83. Groups with different letters represent significantly different results by 1-way ANOVA with Tukey’s HSD post hoc analysis (p < 0.05) for each of the 5 dietary treatment groups (n = 15/treatment): OSD, OSD + F, 76A, 93G, and chow.

### *STUDY 2:* Morphological Changes After 14 Days on High-Fiber Purified Diets

The body weight of animals in all groups were similar at the end of the two week experimental period. Body weight and weight gain in only the 200FOS group were significantly lower compared to the GBD groups (5001, p=0.0498; 5002, p=0.0191) **(Table 5)**. Despite minimal differences in weight gain, rapid changes to lower intestinal morphology were observed after two weeks on certain diets. Representative pictures of cecums and colons from each group indicate that regardless of fiber amount, the cellulose supplemented PDs yielded smaller cecums and shorter colons compared to soluble fiber supplemented PDs or the GBDs **(Figure 3)**. As suggested by the differences in the photos, statistical analysis of the organ weights and lengths showed similar trends and significant differences between groups **(Table 5)**. Significantly shorter colons were observed in the 100CEL (p=0.0045) and 100FOS (p=0.0123) groups compared to the 5002 group, but both PD groups were similar to 5001; however, all other PD groups were statistically similar to both GBD groups (5001 and 5002). Cecum plus colon wall weights for the 200IN and 200FOS groups were similar to one another but significantly elevated compared to all other treatments (200IN and 200FOS vs. most groups, p<0.0001; FOS200 vs. 5002, p=0.0008); 200IN and 200FOS were similar to one another. 100CEL had significantly lower cecum plus colon weights than both GBDs (vs. 5001, p=0.0253; vs. 5002, p<0.0252) and the two inulin groups and 200FOS (100CEL vs. 100IN, p=0.0255; 100CEL vs. 200IN and 200FOS, p<0.0001), while 200CEL increased it slightly and was similar to 100IN, 100FOS, and GBD groups. The 100IN group maintained similar cecum plus colon weights as GBDs. Cecum weights of the different groups followed similar trends as cecum plus colon weights with 200IN and 200FOS being similar to one another but higher compared to all other groups (200IN vs. all groups, p<0.0001; 200FOS vs. 5001, p=0.0011; 200FOS vs. 5002, p=0.0006; 200FOS vs. 100CEL and 200CEL, p<0.0001; 200FOS vs. 100IN, p=0.0031; 200FOS vs. 100FOS, p=0.0009). Cecum weight relative to BW in 100IN tended to be higher than those of 100CEL (p=0.074), but 200IN had significantly higher cecum weights than 200CEL (p<0.0001). The 100IN group allowed for similar cecum weights as for those fed GBDs. Colon weights alone followed a similar trend and were more varied. The 200IN group tended to have higher colon weights than those fed 200CEL, but not statistically higher (p=0.0533) and the addition of more inulin or FOS also tended to increase colon weights. Notably the 100CEL group demonstrated significantly reduced colon weight compared to the GBD groups (5001, p=0.048; 5002, p=0.0186. In contrast, 200CEL maintained similar colonic weights as GBD groups.

**Table 5.**
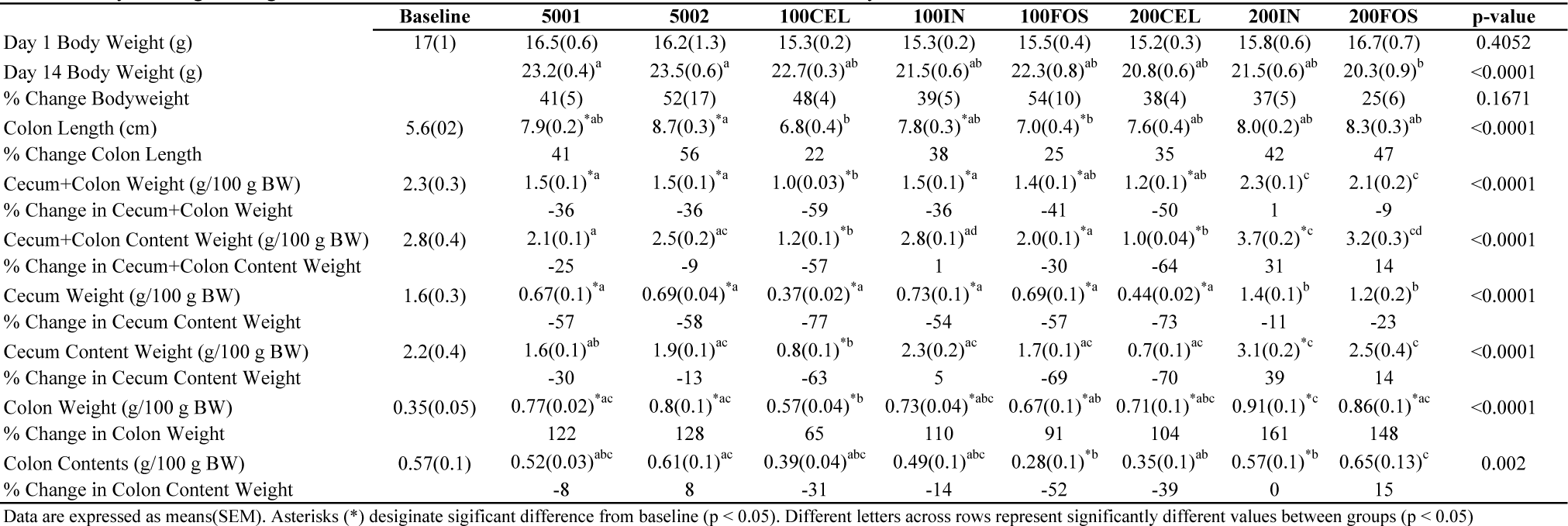
Body and Organ Weights at Baseline and After 2 Weeks on a GB or Purified OSD in Study 2

**Figure 3.**
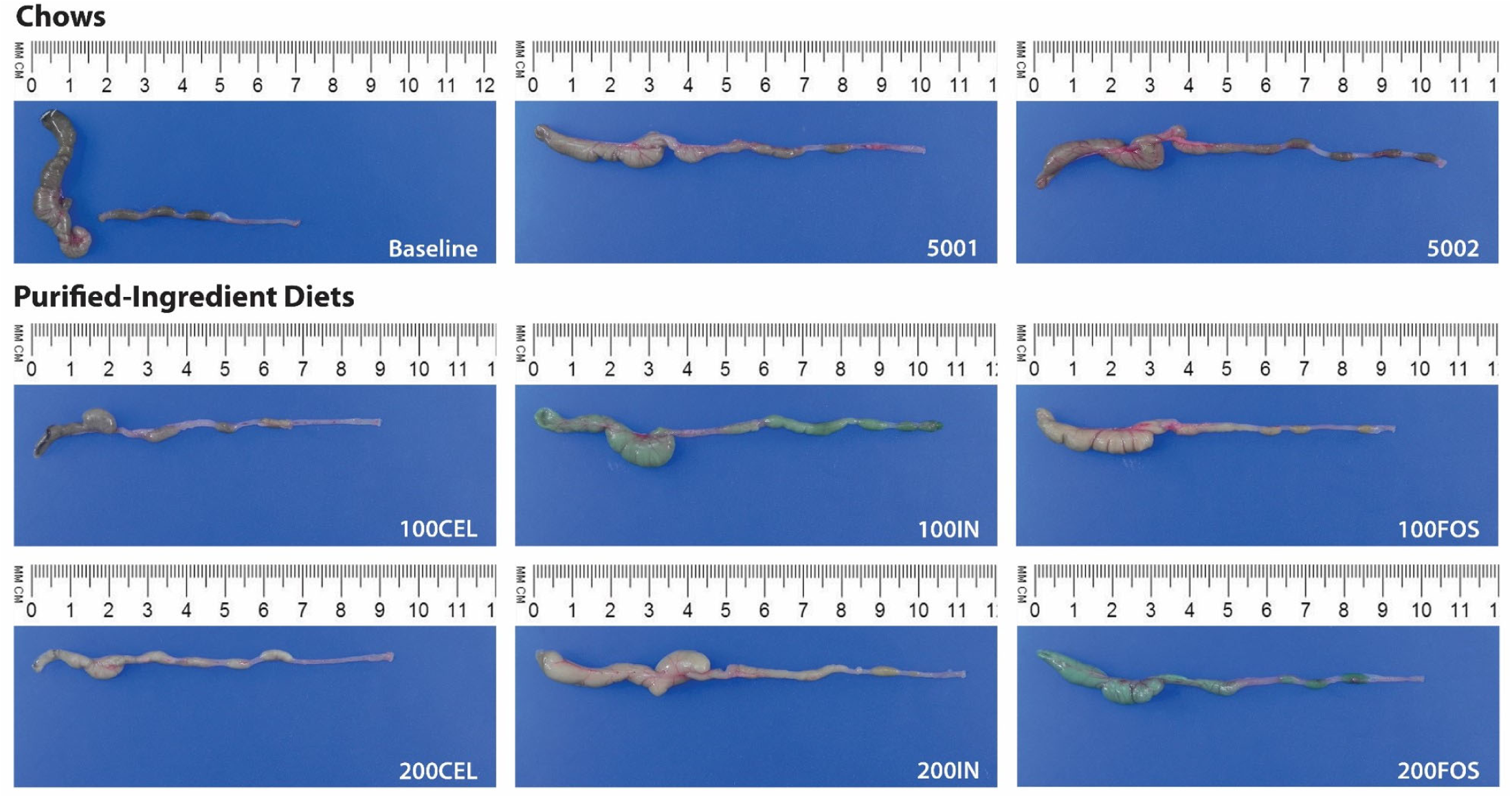
(Study 2). Images of representative cecums and colons for each dietary treatment group after 14 days on either a GBD (5001 or 5002), or high-fiber PDs (100CEL, 100IN, 100FOS, 200CEL, 200IN, 200FOS).

#### Changes to the Prominent Microbial Taxa after 14 Days on the Dietary Treatments

##### a. Alpha and Beta diversity measures

To assess the response of the microbiome to the different dietary treatments, 16S rRNA sequencing was performed on the cecum and colon contents of each animal. When examining differential abundance for both taxa and functions across treatment groups, we found a large number of features at all levels examined that differed between at least one (and usually more) pairwise comparisons of treatment groups. In the cecum samples, 168 ASVs (50 Genera, 25 Families and 7 Phyla), 1342 Ecs, and 282 metacyc pathways were differentially expressed. In the colon samples, it was 165 ASVs (51 Genera, 24 Families and 7 Phyla), 1322 Ecs, and 279 metacyc pathways. Globally, although the diets 5001 and 5002 have slightly different composition, there were no noteworthy or significant changes to report between these two groups. With respect to alpha diversity, the soluble fiber diets (IN and FOS) were similar and both soluble fibers significantly reduced species richness compared to the GBDs and cellulose based diets in cecums and colons for Chao1, Observed, and Shannon diversity indices (**Figure 4**). Dietary soluble fiber treatments significantly influenced alpha diversity metrics for both sites (p <0.001 for all analyses). The GBDs were able to support the greatest number of species in both tissue types regardless of the diversity metric used and in most cases, both the 100CEL and 200CEL groups were also able to maintain a statistically similar number of species as GBDs. Overall, fiber dose did not have a significant impact on alpha diversity. In terms of beta diversity, PCoA plots depicting the Bray-Curtis dissimilarity of ASVs for cecums and colons (**Figure 5)** indicated that treatment groups were quite distinct from one another. In fact, three significantly different clusters of microbial communities were observed in both cecums (p = 0.001) and colons (p = 0.001). Once again, dietary fiber types appeared to be the primary differentiating factor, with the three distinct clusters consisting of soluble fiber-based diets (both IN and FOS clustered together), cellulose-based diets, or GBDs. Fiber dose did not appear to influence beta diversity, as the low dose and high dose treatments for each fiber type were clustered together in both sites of sampling.

**Figure 4.**
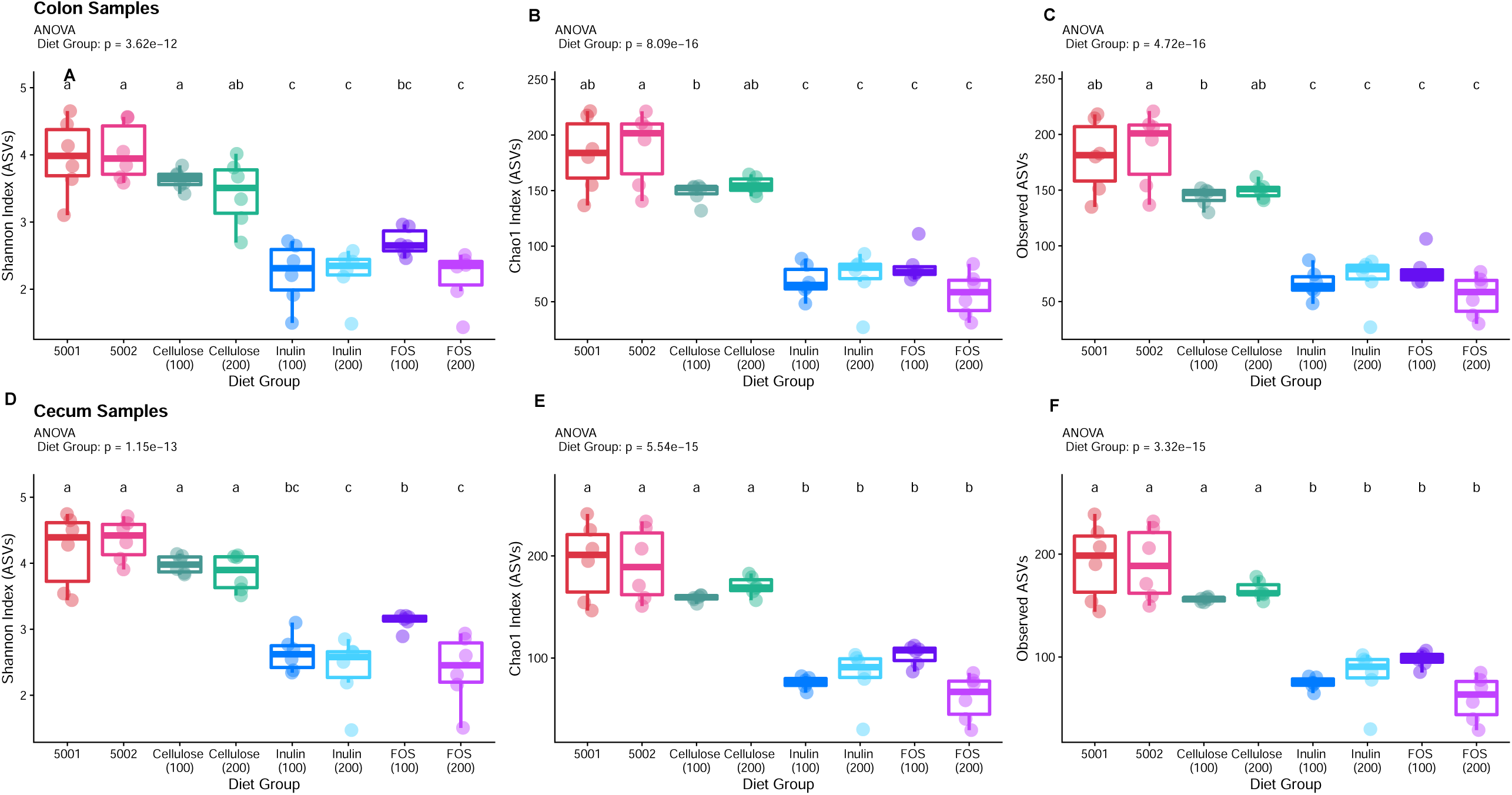
(Study 2). Alpha diversity measurements for Shannon index (A/D), Chao1 index (B/E) and Observed ASVs (C/F) by dietary treatment group for colon (A-C) and cecum (D-F) samples. Data are expressed as means ± SEM for each treatment group. Groups with different letters represent significantly different results by one-way ANOVA with Tukey’s post hoc analysis (p < 0.05) after 14 days on either a GBD (5001 or 5002), or high-fiber PDs (100CEL, 100IN, 100FOS, 200CEL, 200IN, 200FOS) with n = 7-8/group.

**Figure 5.**
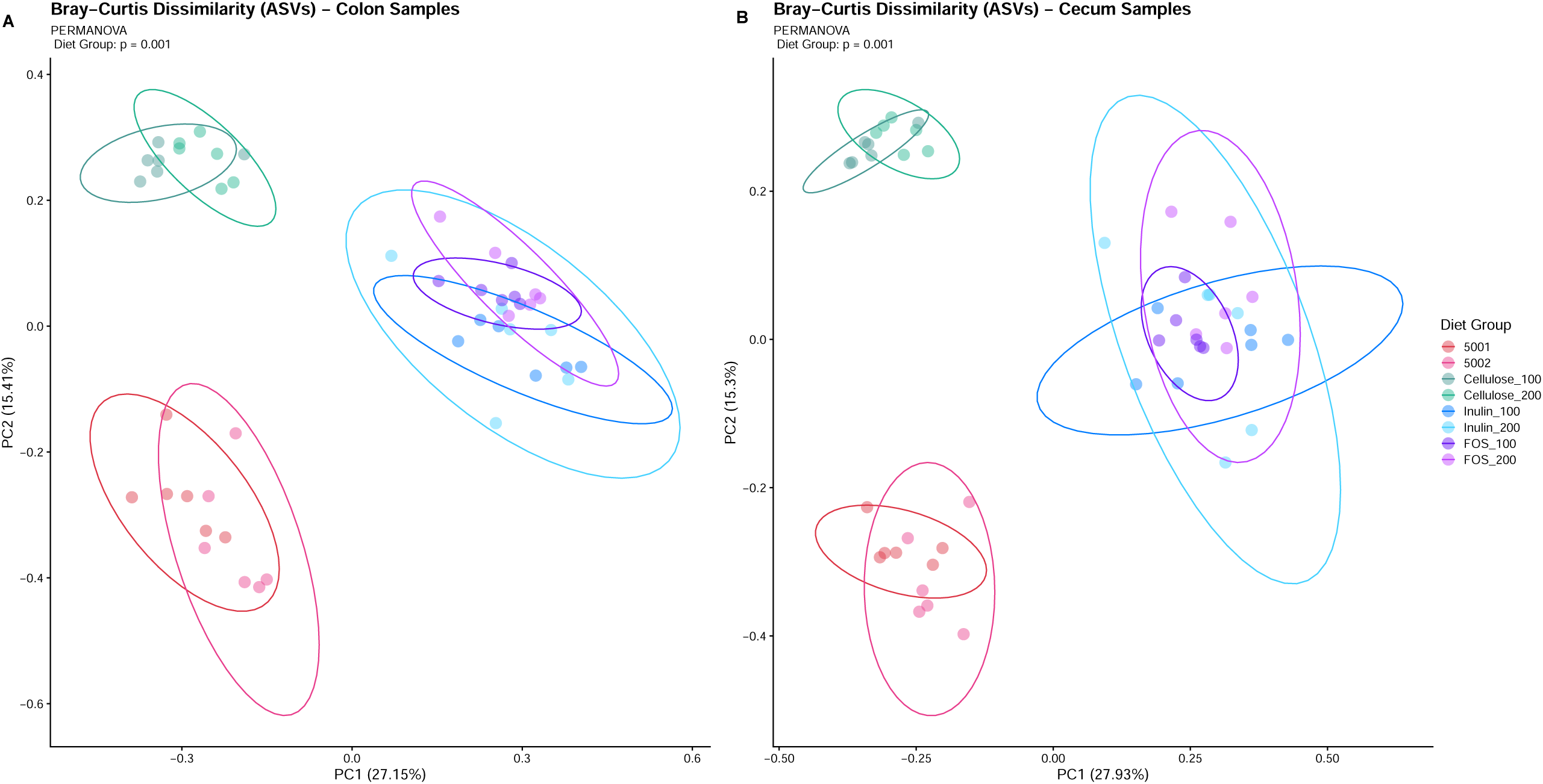
(Study 2). Bray-Curtis Dissimilarity plots for colon (A) and cecum (B) samples after 14 days on either a GBD (5001 or 5002), or high-fiber PDs (100CEL, 100IN, 100FOS, 200CEL, 200IN, 200FOS) with n = 7-8/group.

##### b. Phylum changes

When looking more closely at the relative proportions of taxa, shifts in relative abundance of certain microbes were observed at the phyla and genus level **(Figure 6)** and associated P values for individual microbial taxa differences can be found in the **Supplementary Tables 1-4**. At the phylum level, Firmicutes was the dominant phyla found in the GBD treatment groups, followed by Bacteroidota for cecum samples (**Figure 6A**). The opposite trend was observed in the colon samples, where Bacteroidota was the dominant phylum (**Figure 6B**). Interestingly, all of the soluble fiber diets significantly reduced the Firmicutes/Bacteroidota ratio in the cecum samples **(Supplementary Figure 1A**) compared to the GBDs; however, in the colon, it was not significant at P = 0.025 (except for the 200IN group) (**Supplementary Figure 1B**). The addition of cellulose to the diets led to minor phylum shifts, such as increased abundance of Deferribacterota in the cecums. This shift was significant for 100CEL (p = 0.0226) and 200CEL (p = 0.0183) groups compared to 5002 (**Supplementary Table 1**). No differences were observed between the cellulose diets and the GBDs at the phylum level in the colon samples. In contrast, major phylum-level shifts were observed in both tissue types for the mice fed the soluble-fiber diets. The soluble fiber diets generally reduced the abundance of Firmicutes in both cecum and colon, reaching significance for the 200IN, 100IN, and 200FOS groups compared to both GBDs (p values < 0.039 in each case for both tissue sites) (**Supplementary Table 1 and 2**). In the cecums, marked reductions in Firmicutes abundance alongside elevations in Verrucomicrobiota were observed, particularly for the FOS treatment groups. In fact, Verrucomicrobiota abundance was significantly higher for 100FOS compared to 5001 (p = 0.0137), while the 200FOS group demonstrated significantly lower Firmicutes abundance compared to 5001 (p = 0.012) and 5002 (p = 0.0176) (**Supplementary Table 1**). Specifically in the colon, Verrucomicrobiota abundance was generally elevated compared to the GBDs, reaching significance for 200FOS (p = 0.0133), 100FOS (p = 0.0106) and 100IN (p = 0.0245) compared to 5001, and for 100FOS (p = 0.0411) compared to 5002. Both FOS groups also demonstrated significant elevations in the phylum Actinobacteriota compared to both GBDs (**Supplementary Table 1**). Similar trends were observed in the colon samples. Colon Actinobacteriota populations were also significantly increased by both FOS treatment groups compared to both GBDs (p values < 0.026) (**Supplementary Table 2**).

**Figure 6.**
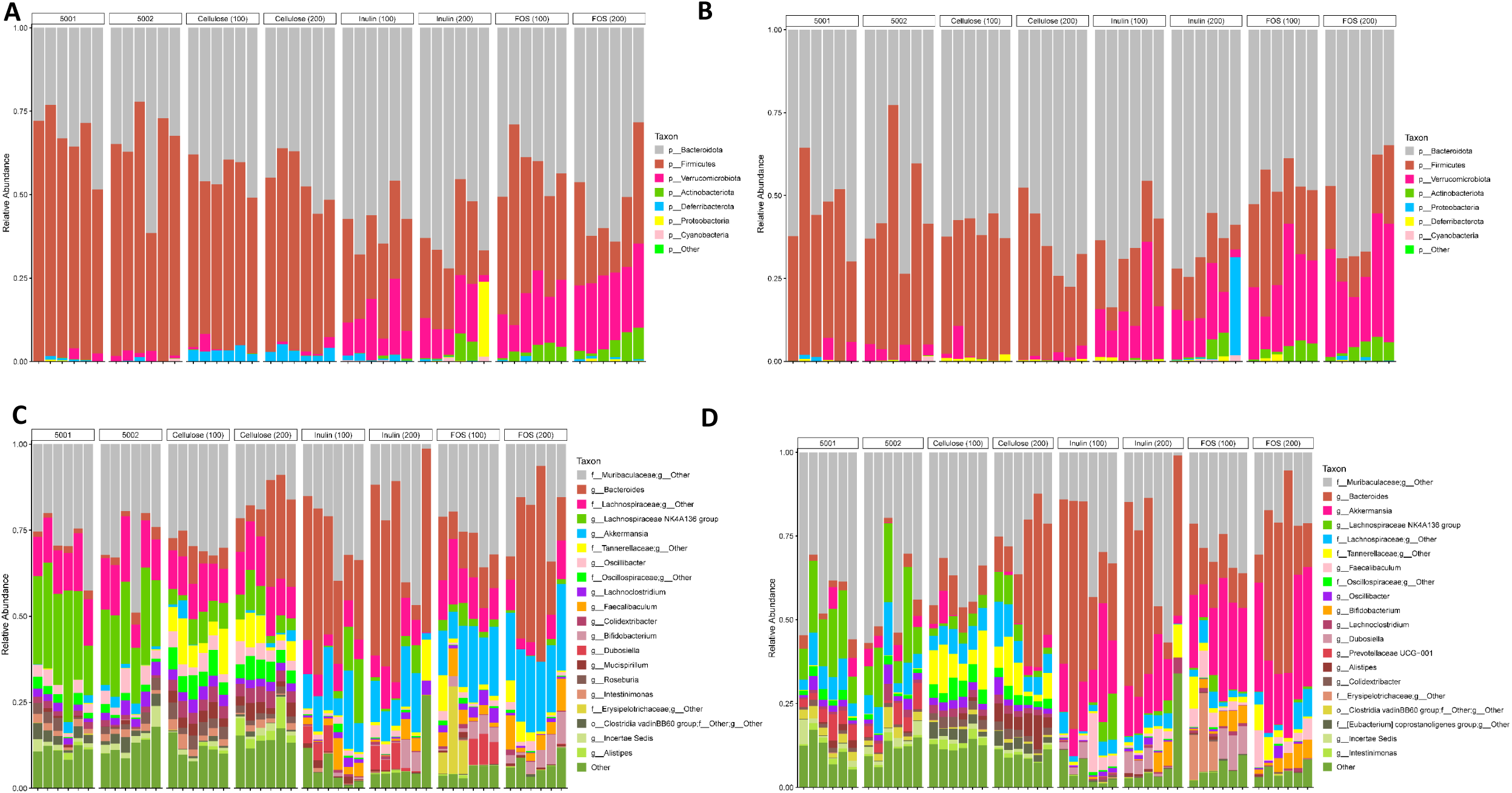
(Study 2). Composition of fiber in the diet modulates the composition of the gut microbiota. Relative abundance of dominant phyla by dietary treatment for cecums (A) and colons (B) and at the ASV level for cecums (C) and colon (D) after 14 days on either a GBD (5001 or 5002), or high-fiber PDs (100CEL, 100IN, 100FOS, 200CEL, 200IN, 200FOS) with n = 8/group.

##### c. Genus level changes

Notable shifts at the genus level were also observed due to the different dietary treatments. In the cecum samples (**Figure 6C**), prominent genera present in the GBDs included the *Lachnospiraceae NK4A136* group and an unclassified genus from the family Muribaculaceae. Interestingly, *Lachnospiraceae NK4A136* group was significantly higher in abundance for both GBDs compared to the cellulose and FOS-based diets, and tended to be higher than 200IN (5001 vs. 200IN, p=0.062, 5002 vs. 200IN, p=0.093), but not different from those fed 100IN (5001 vs. 100IN, p=0.53; 5002 vs. 100IN, p=0.57). The family Oscillospiraceae (undefined genus) was significantly increased by cellulose groups compared to both GBDs and most of the soluble fiber groups, except where 100CEL was not significantly different from 100IN (p=0.14). *Alistipes* was significantly elevated by 200CEL relative to what was found in both GBDs (vs. 5001, p=0.035; vs. 5002, p=0.009) and compared to all soluble fiber groups (p<0.02). All of the soluble fiber diets at both doses, relative to GBDs, significantly increased the relative abundance of *Bifidobacteria, Akkermansia*, and *Faecalibaculum* (GBDs vs. all soluble fiber diets, p≤0.01), while decreasing *Roseburia* (GBDs vs. 200IN or 200FOS diets, p<0.01; vs. 100IN or 100FOS, p<0.05); similar trends were also observed relative to cellulose-based diets (*Bifidobacteria, Akkermansia*, and *Faecalibaculum*, p=0.01; for *Roseburia*, p<0.02). Almost all the PDs caused a significant increase in the relative abundance of genus *Bacteroides* (p≤0.03) except for 5002 relative to 100CEL (p=0.13) and all PDs had more of the family *Tannerellaceae* (genus undefined) compared to GBDs (p≤0.01) **(Figure 6C, Supplementary Table 3)**. *Bacteroides* was significantly increased by soluble fiber diets at lower fiber doses, but not at higher doses compared to the cellulose diets at the same dose (100IN and 100FOS vs. 100CEL, p=0.01; 200IN and 200FOS vs. 200CEL, p≥0.09). Dose of fiber (regardless of type of fiber) had no significant influence on any of the above mentioned genera.

Similar trends in relative abundance at the genera level were observed in the colon samples **(Figure 6D)**. Not surprisingly, the soluble fiber based diets significantly increased the abundance of *Bifidobacteria, Akkermansia*, and *Faecalibaculum* compared to cellulose and GBDs. Unlike in the cecums, *Roseburia* abundance was significantly increased for 100IN (p = 0.0194), 200IN (p = 0.0105), and 200FOS (p = 0.011) relative to 5002; however, there were no differences compared to the 5001 group for any PDs (**Supplementary Table 6**). Similar to the cecum samples, all PD groups increased the relative abundance of the genera *Bacteroides* (except 100CEL vs. 5002, p=0.06) and *Prevotella*_UCG-001, and the Tannerellaceae family (undefined genus) compared to GBDs. The family Tannerellaceae (undefined genus) was also significantly higher for 100CEL compared to 100IN (p=0.02). *Bacteroides* was significantly increased by 100IN and 100FOS relative to 100CEL (p=0.01) and for 200FOS relative to 200CEL (p=0.045), but not changed by fiber dose. In contrast, an undefined genus in the family Muribaculaceae was significantly reduced by 100IN and 100FOS relative to 100CEL (P≤0.03) and 200FOS vs. 200CEL (P=0.012). Collectively, these differences clearly demonstrate that different fiber types support the growth of different microbes present in the GI tract and that both soluble fibers tended to support similar microbial growth.

#### Predicted Metabolic Functions of Microbiota

The predicted function of the gut microbiota with respect to metabolism in response to the dietary treatments was evaluated using the *PICRUSt* program. When examining the gene counts for specific categories of metacyc pathways for cecum (**Figure 7**) and colon (**Supplementary Figure 2**) samples, key differences between treatment groups were observed. These pathways were related to carbohydrate digestion and absorption, fructose and mannose metabolism, galactose metabolism, starch and sucrose metabolism, fatty acid metabolism, and fatty acid biosynthesis (**Figure 7**). Generally speaking, the PDs had higher gene counts than GBDs for all the six tested pathways in the cecal samples, but only the 200CEL and 200IN treatments significantly elevated the gene counts for each of these pathway groupings compared to both the GBDs. Only the cellulose group produced a significant effect of fiber dose on pathway upregulation, as the 200CEL group demonstrated higher gene counts than the 100CEL group for all the pathways except for carbohydrate degradation and absorption, which failed to reach significance. Unlike inulin, neither of the FOS diets significantly affected the metabolic pathways compared to GBDs regardless of the dose. Fewer differences were observed in terms of metabolic pathways in the colon samples (**Supplementary Figure 2**). PDs demonstrated higher gene counts related to carbohydrate degradation and absorption, with the 200CEL, 200IN, and the 200FOS groups reaching significantly higher levels compared to the GBDs. Galactose metabolism was affected by 200CEL and 200FOS relative to GBDs. In the case of fructose and mannose metabolism, starch and sucrose metabolism, fatty acid metabolism, and fatty acid biosynthesis, only the 200CEL dietary treatment produced significantly higher gene counts compared to all other groups. As in the cecum, 200CEL group increased gene counts compared to the 100CEL group in most pathways except for carbohydrate degradation and absorption, which failed to reach significance. The soluble fiber groups, regardless of dose, did not significantly modulate metabolic pathway gene counts in the colon.

**Figure 7.**
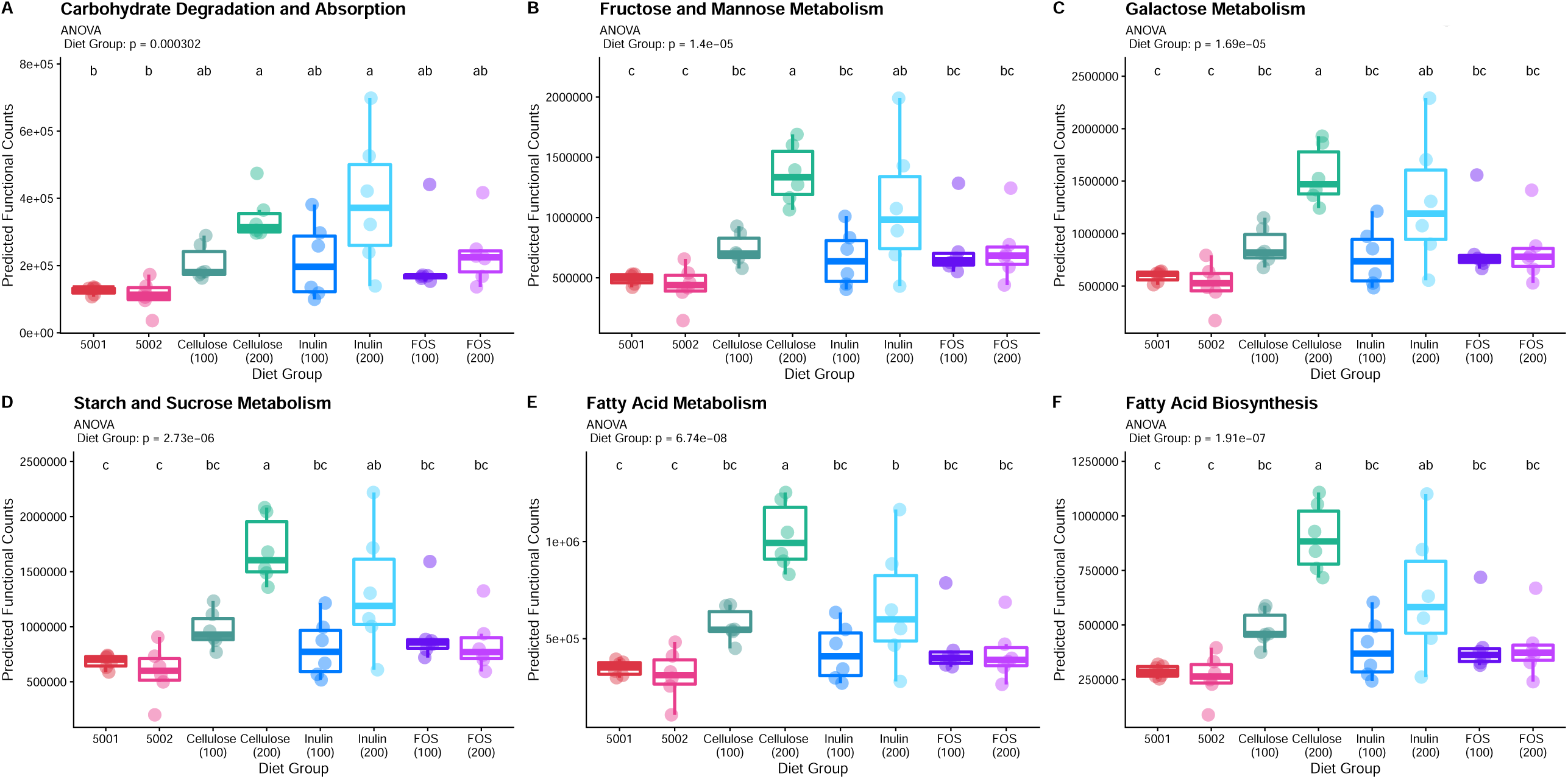
(Study 2). Predicted carbohydrate and lipid metabolism of bacterial communities in the cecum using the PICRUSt program after 14 days on either a GBD (5001 or 5002), or high-fiber PDs (100CEL, 100IN, 100FOS, 200CEL, 200IN, 200FOS) with n = 8/group. (A) Carbohydrate digestion and absorption; (B) fructose and mannose metabolism; (C) galactose metabolism; (D) starch and sucrose metabolism; (E) fatty acid metabolism; (F) fatty acid biosynthesis. Data are expressed as means ± SEM for each treatment group. Groups with different letters represent significantly different results by 1-way ANOVA with Tukey’s HSD post hoc analysis (p < 0.05).

## DISCUSSION

PDs are an essential element of nutrition research and it is important to consider the effect that background diet may have on the expressed phenotype. Since the AIN diets were designed 4 decades ago, we have learned more about how different nutrients alter metabolic profile of mice and rats and this knowledge can be applied to improve these diets for future studies. he AIN committee did allude to using a different source of carbohydrate in lieu of sucrose due to potential influences of this ingredient on metabolic disorders, they did not make any specific recommendations on adjusting the fiber levels in these diets(22). The current recommendation for 5% cellulose, an insoluble fiber, provides little fermentable dietary substrate accessible to the intestinal microbiota. Current data clearly show that this lack of soluble fiber leads to dramatic effects on gut and metabolic health in mice(8,17). We therefore wished to further understand how revised versions of the AIN diets with reduced sucrose and higher amounts of total fiber, including soluble fiber, affected metabolic and gut health compared with GBDs.

In our chronic feeding study, fiber and sucrose alterations in the two OSDs used in this study did not result in significant changes to overall adiposity index (combining all adipose depots relative to carcass weight) relative to the GBD. Leptin was elevated in mice fed either the 93G or OSD compared to those fed the GBD, suggesting that the reduced sucrose and fiber change had little impact on these parameters. However, the addition of more fiber as cellulose suppressed this effect and these mice had similar levels as those fed the GBD. While it is unknown how added cellulose led to a reduced leptin level, a trend for reduced adipose depots in these animals relative to other PDs may have partially accounted for this difference. Furthermore, there was also a slight but significant increase in liver triglycerides in the OSD fed mice compared to those fed the GBD, while those fed the higher fiber OSD diet had similar levels as those fed GBD. Overall, these changes suggested that total fiber content is more important to maintaining these static biochemical parameters. However, our data suggested that the addition of inulin was key to maintaining glucose tolerance and both OSD groups had similar glucose tolerance compared to those fed the GBD. In addition, replacement of sucrose with glucose derived carbohydrates may also have benefitted these mice fed OSD as sucrose may induce metabolic disease in rats and mice (23,24). While sucrose levels were reduced to 10% in the AIN-93G diet relative to 50% in the AIN-76A diet, it was not completely removed due to pelleting and palatability concerns by the AIN committee (25). However, even relatively lower levels of sucrose may elicit changes in glucose tolerance over a chronic feeding period, which is due to the fructose component of this carbohydrate (26).

Closer examination of how the type and amount of fiber in PDs influenced gut health (Study 2) indicated that a replacement of cellulose with soluble fiber-based PD prevented the rapid cecum and colon weight loss associated with traditional cellulose-based PDs. Both types of soluble fiber at low and high doses were capable of maintaining cecum and colon weights relative to GBD-fed animals. This phenomenon has also been observed by us previously in the context of a high-fat diet, where an inulin-based high-fat PD helped to maintain cecum and colon weight changes compared to cellulose-based high-fat PD. This effect was attributed to the microbiota as inulin’s ability to promote colon and cecum mass was completely absent in germ-free mice (17). Alongside changes to organ morphology, soluble fiber based PDs caused rapid and dramatic changes to the microbiota in contrast to GBDs, regardless of fiber content, including reduced species richness (alpha diversity) in both cecum and colon tissues. This may be in part due to the differences in fiber diversity among these diets as GBDs contain diverse fiber contents, which may support the growth of more diverse microbial species, leading to greater alpha diversity values (27). Moreover, dietary intervention studies in humans in which only one fiber type is added to the diet typically do not translate to increased alpha diversity metrics (28). Still, our results indicate that cellulose alone was able to support greater species richness in a PD compared to inulin and FOS. This could be partially because the fiber types present in GBDs are predominantly insoluble types, like cellulose (4). Previous data suggest that cellulose is important for age-related diversification of the intestinal microbiota (29) and thus our results indicate that it may be better to continue including cellulose in future designs of PD. In fact, a recent study suggested that adding inulin to a lower fat diet containing cellulose tended to increase alpha diversity relative to cellulose alone, suggesting the importance of maintaining cellulose to preserve species richness (30). Beta diversity analysis indicated that each fiber type (GBDs, cellulose based, or soluble fiber based) supported distinctly different communities of bacteria, regardless of fiber dose. These data also suggested that both GBDs supported growth of similar microbial communities, which may not be surprising given they contain similar ingredients and also insoluble and soluble fiber contents. This has been documented in other studies and together, these findings indicate that single sources of purified fibers as part of a PD, even at higher concentrations, may not support similar microbial species as the GBDs do. It is likely the case that multiple purified fibers used in combination would optimally support the microbiome and thus are necessary to achieve closer similarity to GBDs, regardless of the dose.

Although switching from GBD to PDs generally led to reduced species richness and supported different microbial species compared to GBDs, some of the changes to dominant microbial taxa could be beneficial and ultimately yield better health outcomes in rodents, especially if soluble fibers were to be added to a PD alongside cellulose in the future. The role of the Firmicutes to Bacteriodota ratio in the onset of obesity itself remains controversial (21,31). However, increased relative abundance of Bacteriodota and other major phyla have demonstrated positive health effects beyond obesity remediation. For example, a study by Rabot et al (31) indicated that an increased abundance of Bacteriodota was associated with improved glucose tolerance in a cohort of mice fed a high-fat diet. Moreover, it has been demonstrated that inclusion of prebiotic fibers, like inulin and FOS, in the diet are linked with higher circulating levels of GLP-1, a metabolic hormone with antidiabetic effects (32,33). Together, these observations could partially explain why adding soluble fiber such as inulin to PDs is associated with improved metabolic health compared to the insoluble fiber, cellulose based low-fiber, high-sucrose AIN diets as seen from results of our first study.

There is a growing body of evidence suggesting that certain microbial genera positively influence metabolic health. *Akkermansia* spp., which belong to the Verrucomicrobia phylum (both of which were elevated due to the inclusion of soluble fiber in the diet), have demonstrated the ability to maintain gut barrier integrity. As such, it is suspected that *Akkermansia* spp. may exert anti-inflammatory properties, which in turn could contribute to overall metabolic health, given that chronic inflammation is associated with insulin resistance and diabetes (34,35). Other studies have indicated that administration of FOS in rodent diets appears to increase *Akkermansia* abundance in the gut (doses as low as 0.3 gm/d, which translate roughly to 60 gm FOS/kg diet), while diet-induced obesity appears to consistently reduce abundance of this genera (36,37). Our study suggests that inulin and FOS may also increase the relative abundance of this metabolically active genus. *Bifidobacteria*, which are a subspecies of Actinobacteria have historically been categorized as health promoting microbes (38). It is believed that these bacteria play a role in stimulating host innate immunity, and may also enhance the ability of *Bacteroides* to metabolize carbohydrates (35,38). *Bifidobacteria* also appear to reduce gut permeability, similarly to *Akkermansia*, and reductions in *Bifidobacteria* populations are found in diet-induced obese rodents (39). The results from our study indicated that the addition of soluble fiber to a PD can increase the abundance of *Bifidobacteria* relative to a GBD, and both cellulose and soluble fibers may increase *Bacteroides*. Another microbial genus of interest is *Roseburia*, which is one of the most abundant genera of the Firmicutes phylum. Despite general observations that overall Firmicutes abundance is reduced in lean individuals relative to obese individuals, *Roseburia* spp. tend to be more abundant in lean individuals (40). They also generally increase in relative abundance in response to diets rich in FOS and inulin (41). However, in our study, all the soluble-fiber based diets decreased *Roseburia* relative abundance compared to both GBDs and cellulose based diets in the cecum and colon tissues. While these data are contrary to other reported findings(41), it does support the notion that multiple fiber types may be necessary to support the optimal growth of a consortium of beneficial bacteria. *Roseburia* spp. are well known for their ability to produce high concentrations of butyrate, a potent short-chain fatty acid that is known to promote colon health (42). Production of this metabolite, by *Roseburia* or other genera, would certainly be indicative of the diet supporting a healthy gut. While short-chain fatty acids were not measured in this study, previous collaborative studies suggested that replacement of cellulose with around 5% inulin in the context of a PD with low fat contents (similar to AIN and OSD) elevated levels of fecal SCFA to a similar level as those fed a GBD (17).

Further insights regarding the functional capacity of the microbiome in response to the dietary treatments were explored using the PICRUSt program. Despite the limitation that this software can only predict functional differences in the microbiota using marker-gene sequencing techniques(43), several interesting trends with respect to the treatments were observed. Firstly, it was evident from the metabolic pathways analysis (**Figure 7 and Supplementary Figure 2**) that the dietary fibers, in particular the higher dose inulin group, had a greater impact on genes related to metabolism in the cecums compared to the colons. The only group with significantly higher metabolic pathway gene counts in both cecum and colon was the 200CEL group. This could partially be explained by the fact that the cecum is the primary site of fermentation in mice, so much of the inulin may have been rapidly fermented in the cecum, leaving little to no substrate available for use by microbes in the latter portions of the gut (44). However, significant differences in carbohydrate degradation and absorption were still found for inulin and FOS PDs in the colon, suggesting that there was still enough remaining fructans from these sources for microbes residing in the latter portions of the gut. Another partial explanation for the high-dose cellulose group being the only treatment group significantly influencing colonic microbial metabolism is the fact that cellulose is poorly fermented by non-ruminant mammals such as mice and humans (45). Unlike the fast-fermenting soluble fibers, it was likely that still some undigested cellulose passed into the colon, which could be utilized by the microbes residing there. With respect to the functionality of the gut microbiome in response to the dietary treatments, it is also worth noting that there appeared to be some dose-related effects, which were not observed when looking at microbial abundance and diversity. Higher doses of cellulose and inulin tended to increase gene counts in specific pathways to a greater extent than low-dose counterparts in the cecum samples. In other words, greater substrate availability (i.e. higher dosage) translated to greater potential metabolic activity in these tissues. Taken together, all of these findings indicate that cellulose likely plays some role in the metabolic functionality of the gut microbiome, in part by maintaining microbial richness similar to GBDs, but soluble fiber may help to support normal colon physiology and also promote the growth of health beneficial microbial genera, which lead to improved gut morphology relative to cellulose alone. Overall, higher concentrations of fiber support greater microbial activity compared to the AIN recommended dose of 5% in a rodent diet.

In summary, it is clear that these changes to the formulation of traditional AIN PDs (increased amount and addition of soluble fiber with replacement of sucrose with glucose derived carbohydrate sources) provide certain improvements to metabolic health, which may have been in part due to changes in the gut microbiota profile. However, to more closely mimic gut microbiota in mice fed GBDs, the addition of multiple, diverse fiber sources will likely be required (4). Our results suggest that a mixture of soluble and insoluble fiber types, present at higher concentrations than the AIN formulations and without the addition of sucrose, may be help to maintain microbial richness similar to a GBD while also supporting greater functionality of the microbiome. However, it is difficult to say whether one type of soluble fiber is more beneficial than the other and whether a change in fiber in a PD would allow for a similar microbiome relative to all GBDs. Thus, a PD containing multiple sources of soluble fiber including the ones not used in this study (for e.g. mannans, beta-glucan, hemicelluloses, pectin etc.) may need to be developed. Future efforts should be directed towards determining the optimal ratios of soluble and insoluble fibers in PDs, as well as exploring how these changes to the gut microbiome may influence animal health in longer-term studies. It is also critical to focus not only on relative microbiome shifts and predicted functionality, but also to examine concentrations of circulating microbial metabolites such as SCFAs to better understand the potential benefits for metabolic health. Given the differences between PDs and GBDs, it should be clear that these two diet types should not be compared against each other while determining dietary effects on a given phenotype. This is particularly apparent when determining the theoretical underpinnings of how dietary effects on gut health and the microbiome drive changes in metabolic health outcomes, as suggested previously (17,46,47). Unfortunately, GBDs are frequently used as controls for many experimental studies testing effect of a PD (e.g. high-fat diet studies), which leads to misinterpretation of results. While having a matched PD allows one to compare how a given change in diet is altering the rodent phenotype, one particular concern from the research community is that rodents consuming control PDs, although lower in fat, are typically not as metabolically healthy as those fed a GBD. Thus, it is imperative that the research community focus on improving the formulation to mitigate some of the adverse changes associated with consumption of PDs. A metabolically healthy control PD would greatly help the research community to decipher nutrient related phenotypic differences in a wide range of scientific domains. While we mainly focused on sucrose levels and the type/concentration of the fiber, future studies should also examine the type and level of fat in order to optimize the formulation of a metabolically healthy PD, as recently discussed regarding the AIN series formulas (8). While we understand that the current study is limited to certain metabolic parameters and the use of prediction software to assess microbial functionality, this study reinforces the notion that lab animal diets must be formulated and selected with utmost care, as the gut microbiome is easily influenced by diet and shifts in the populations of microbes may impact study outcomes.

## List of abbreviations

PD: Purified diet
GB: Grain-based
76A: AIN-76A rodent diet
93G: AIN-93G rodent diet
CEL: Cellulose
IN: Inulin
FOS: Fructo-oligosaccharides
OSD: Open Standard Diet
EC: Enzyme Commission
OGTT: Oral glucose tolerance test
PERMANOVA: Permutational Analysis of Variance
ANOVA: Analysis of Variance

## ACKNOWLEDGEMENTS

The authors would like to thank Dr. Kelly Swanson for his assistance with analyzing the fiber content of the GBDs. We would also like to thank Alex La Reau at Diversigen for his assistance with microbiome data analysis.

## AUTHOR CONTRIBUTIONS

MP designed the studies and the diets, facilitated the completion of the studies with CROs, and analyzed the data from Study 1. LG and SR wrote the manuscript and interpreted the data from Study 2. All authors contributed to revisions and final submission of the manuscript.

## TABLES

**Supplementary Table 1:** P-values for individual microbial taxa differences (pairwise comparisons, Tukey HSD) at the phylum level in the cecum samples.

**Supplementary Table 2:** P-values for individual microbial taxa differences (pairwise comparisons, Tukey HSD) at the phylum level in the colon samples.

**Supplementary Table 3:** P-values for individual microbial taxa differences (pairwise comparisons, Tukey HSD) at the genus level in the cecum samples.

**Supplementary Table 4:** P-values for individual microbial taxa differences (pairwise comparisons, Tukey HSD) at the genus level in the colon samples.

## FIGURE LEGENDS

**Supplementary Figure 1: (**Study 2). Composition of fiber in the diet modulates the composition of the gut microbiota. Changes in Firmicutes/Bacteroidetes ratio in different groups dietary treatment for cecums (A) and colons (B) after 14 days on either a GBD (5001 or 5002), or high-fiber PDs (100CEL, 100IN, 100FOS, 200CEL, 200IN, 200FOS) with n = 8/group.

**Supplementary Figure 2**. (Study) Predicted carbohydrate and lipid metabolism of bacterial communities in the colon using the PICRUSt program after 14 days on either a GBD (5001 or 5002), or high-fiber PDs (100CEL, 100IN, 100FOS, 200CEL, 200IN, 200FOS) with n = 6/group. (A) Carbohydrate digestion and absorption; (B) fructose and mannose metabolism; (C) galactose metabolism; (D) starch and sucrose metabolism; (E) fatty acid metabolism; (F) fatty acid biosynthesis. Data are expressed as means ± SEM for each treatment group. Groups with different letters represent significantly different results by 1-way ANOVA with Tukey’s HSD post hoc analysis (p < 0.05).

## Notes

**Conflict of Interest and Funding Disclosure:** Research Diets, Inc. provided the funding for this research work. SR and MP are employees of Research Diets Inc., the manufacturer of the purified ingredient diets utilized in these studies.

### Competing Interest Statement

SR and MP are employees of Research Diets Inc., the manufacturer of the purified ingredient diets utilized in these studies.

