## Supplemental Tables 1 and 2 for "Addition of soluble fiber in low fat purified diets improves gut and metabolic health compared to traditional AIN diets"

**Supplementary Table 1. P-values for individual microbial taxa differences (pairwise comparisons, Tukey HSD) at the phylum level in the cecum samples.**

| Taxon | 5001__5002 | 5001__200CEL | 5001__200IN | 5001__200FOS | 5001__100CEL | 5001__100IN | 5001__100FOS | 002__200CEL | 5002__200IN | 5002__200FOS | 5002__200CEL | 5002__100IN | 5002__100FOS |
|---|---|---|---|---|---|---|---|---|---|---|---|---|---|
| k__Bacteria;p__Actinobacteriota | 0.8828 | 0.4846 | 0.5881 | 0.0153 | 0.7718 | 0.8292 | 0.0115 | 0.7264 | 0.5454 | 0.0252 | 0.8386 | 0.8034 | 0.0318 |
| k__Bacteria;p__Bacteroidota | 0.8541 | 0.7900 | 0.0368 | 0.0622 | 0.8157 | 0.1308 | 0.5009 | 0.8521 | 0.2060 | 0.1791 | 0.8544 | 0.2617 | 0.5047 |
| k__Bacteria;p__Cyanobacteria | 0.6600 | NA | 0.4854 | NA | NA | 0.0174 | 0.5631 | 0.4747 | 0.8111 | 0.1984 | 0.4494 | 0.2332 | 0.3801 |
| k__Bacteria;p__Deferribacterota | 0.4843 | 0.0634 | 0.0969 | 0.1766 | 0.0911 | 0.7998 | 0.2829 | 0.0183 | 0.8465 | 0.6193 | 0.0226 | 0.2866 | 0.8844 |
| k__Bacteria;p__Firmicutes | 0.7916 | 0.4388 | 0.0125 | 0.0120 | 0.5667 | 0.0123 | 0.0919 | 0.7347 | 0.0124 | 0.0176 | 0.7266 | 0.0234 | 0.1089 |
| k__Bacteria;p__Other | 0.6699 | 0.7437 | NA | 0.4309 | 0.5969 | 0.5539 | 0.6311 | 0.7828 | 0.1560 | 0.3261 | 0.8169 | 0.3811 | 0.4496 |
| k__Bacteria;p__Proteobacteria | 0.6678 | 0.4709 | 0.3699 | 0.8155 | 0.5835 | 0.2465 | 0.1900 | 0.7729 | 0.0195 | 0.1609 | 0.7417 | 0.4618 | NA |
| k__Bacteria;p__Verrucomicrobiota | 0.5524 | 0.5840 | 0.2424 | 0.0651 | 0.9031 | 0.1205 | 0.0137 | 0.7744 | 0.6069 | 0.1273 | 0.2690 | 0.3588 | 0.0809 |

| Taxon | 200CEL__200IN | 00CEL__200FOS | 00CEL__100CEL | 00CEL__100IN | 00CEL__100FOS | 200IN__200FOS | 00IN__100CEL | 200IN__100IN | 200IN__100FOS | 200FOS__100CEL | 200FOS__100IN | 200FOS__100FOS | 100CEL__100IN | 100CEL__100FOS | 100IN__100FOS |
|---|---|---|---|---|---|---|---|---|---|---|---|---|---|---|---|
| k__Bacteria;p__Actinobacteriota | 0.1478 | 0.0055 | 0.8645 | 0.3792 | 0.0068 | 0.4525 | 0.2676 | 0.6483 | 0.0904 | 0.0075 | 0.0279 | 0.6434 | 0.6720 | 0.0072 | 0.0157 |
| k__Bacteria;p__Bacteroidota | 0.0261 | 0.0241 | 0.8339 | 0.0489 | 0.2933 | 0.8093 | 0.0872 | 0.7432 | 0.1955 | 0.0762 | 0.7713 | 0.3798 | 0.1484 | 0.5944 | 0.3078 |
| k__Bacteria;p__Cyanobacteria | 0.2053 | NA | NA | 0.0065 | 0.6454 | 0.3590 | 0.2308 | 0.7288 | 0.2913 | NA | 0.0161 | 0.5737 | 0.0090 | 0.6490 | 0.0112 |
| k__Bacteria;p__Deferribacterota | 0.0064 | 0.0059 | 0.8543 | 0.0081 | 0.0070 | 0.6708 | 0.0080 | 0.2299 | 0.6382 | 0.0065 | 0.3309 | 0.8906 | 0.0100 | 0.0089 | 0.3826 |
| k__Bacteria;p__Firmicutes | 0.0066 | 0.0070 | 0.8523 | 0.0076 | 0.2139 | 0.8229 | 0.0091 | 0.5702 | 0.0208 | 0.0095 | 0.5139 | 0.0965 | 0.0123 | 0.4179 | 0.0515 |
| k__Bacteria;p__Other | 0.1860 | 0.4035 | 0.8125 | 0.4302 | 0.5716 | 0.6471 | 0.0587 | 0.5339 | 0.3143 | 0.1558 | 0.7288 | 0.6998 | 0.2186 | 0.3530 | 0.6321 |
| k__Bacteria;p__Proteobacteria | 0.0082 | 0.0751 | 0.8297 | 0.5428 | 0.6135 | 0.6475 | 0.0104 | 0.0180 | 0.0105 | 0.0970 | 0.0809 | 0.1197 | 0.4787 | 0.4686 | 0.6218 |
| k__Bacteria;p__Verrucomicrobiota | 0.1619 | 0.0286 | 0.7585 | 0.0881 | 0.0129 | 0.3890 | 0.0225 | 0.7141 | 0.0309 | 0.0067 | 0.3185 | 0.3959 | 0.0124 | 0.0071 | 0.0370 |

**Supplementary Table 2: P-values for individual microbial taxa differences (pairwise comparisons, Tukey HSD) at the phylum level in the colon samples.**

| Taxon | 5001__5002 | 5001__200CEL | 5001__200IN | 5001__200FOS | 5001__100CEL | 5001__100IN | 5001__100FOS | 002__200CEL | 5002__200IN | 5002__200FOS | 5002__200CEL | 5002__100IN | 5002__100FOS |
|---|---|---|---|---|---|---|---|---|---|---|---|---|---|
| k__Bacteria;p__Actinobacteriota | 0.8540 | 0.6796 | 0.3899 | 0.0113 | 0.8612 | 0.8008 | 0.0099 | 0.9061 | 0.5678 | 0.0228 | 0.8357 | 0.8093 | 0.0257 |
| k__Bacteria;p__Bacteroidota | 0.7946 | 0.4170 | 0.1195 | 0.2967 | 0.6780 | 0.3250 | 0.6383 | 0.8488 | 0.0591 | 0.1641 | 0.8336 | 0.1898 | 0.3380 |
| k__Bacteria;p__Cyanobacteria | 0.8358 | 0.3090 | 0.5855 | 0.1711 | 0.4432 | 0.0255 | 0.2344 | 0.5892 | 0.6855 | 0.2559 | 0.6379 | 0.1865 | 0.3277 |
| k__Bacteria;p__Deferribacterota | 0.8850 | 0.1748 | 0.4215 | 0.4490 | 0.1204 | 0.8120 | 0.7264 | 0.4872 | 0.5973 | 0.8211 | 0.1776 | 0.6755 | 0.8386 |
| k__Bacteria;p__Firmicutes | 0.8352 | 0.7849 | 0.0171 | 0.0388 | 0.8573 | 0.0287 | 0.4055 | 0.8555 | 0.0106 | 0.0348 | 0.8266 | 0.0351 | 0.2286 |
| k__Bacteria;p__Other | 0.8620 | 0.7885 | 0.2769 | 0.3044 | 0.8184 | 0.4438 | 0.3875 | 0.8041 | 0.2718 | 0.2950 | 0.7256 | 0.5042 | 0.3729 |
| k__Bacteria;p__Proteobacteria | 0.2206 | 0.0996 | 0.5347 | 0.5657 | 0.3105 | 0.0502 | 0.0474 | NA | 0.0136 | 0.4513 | 0.7497 | 0.6222 | NA |
| k__Bacteria;p__Verrucomicrobiota | 0.3892 | 0.3251 | 0.1760 | 0.0133 | 0.7239 | 0.0245 | 0.0106 | 0.8748 | 0.7510 | 0.0692 | 0.5565 | 0.3219 | 0.0411 |

| Taxon | 200CEL__200IN | 00CEL__200FOS | 00CEL__100CEL | 00CEL__100IN | 00CEL__100FOS | 200IN__200FOS | 00IN__100CEL | 200IN__100IN | 200IN__100FOS | 200FOS__100CEL | 200FOS__100IN | 200FOS__100FOS | 100CEL__100IN | 100CEL__100FOS | 100IN__100FOS |
|---|---|---|---|---|---|---|---|---|---|---|---|---|---|---|---|
| k__Bacteria;p__Actinobacteriota | 0.4365 | 0.0081 | 0.8514 | 0.7507 | 0.0096 | 0.2243 | 0.2861 | 0.4389 | 0.0605 | 0.0076 | 0.0159 | 0.7231 | 0.8123 | 0.0118 | 0.0110 |
| k__Bacteria;p__Bacteroidota | 0.0156 | 0.0320 | 0.8437 | 0.0491 | 0.1245 | 0.6154 | 0.0328 | 0.4967 | 0.0578 | 0.0672 | 0.7220 | 0.4674 | 0.0906 | 0.2939 | 0.4554 |
| k__Bacteria;p__Cyanobacteria | 0.2158 | NA | NA | 0.0094 | NA | 0.3083 | 0.2163 | 0.7601 | 0.1924 | NA | 0.0141 | NA | 0.0110 | NA | 0.0092 |
| k__Bacteria;p__Deferribacterota | 0.0458 | 0.0077 | 0.6988 | 0.2117 | 0.2220 | 0.7148 | 0.0206 | 0.3978 | 0.5270 | 0.0075 | 0.4678 | 0.8730 | 0.0597 | 0.1158 | 0.7162 |
| k__Bacteria;p__Firmicutes | 0.0071 | 0.0094 | 0.8879 | 0.0121 | 0.1286 | 0.5559 | 0.0118 | 0.2959 | 0.0070 | 0.0155 | 0.7755 | 0.1995 | 0.0137 | 0.2282 | 0.0674 |
| k__Bacteria;p__Other | 0.1202 | 0.1137 | 0.9090 | 0.2210 | 0.1504 | NA | 0.0967 | 0.5292 | NA | 0.1175 | 0.6160 | NA | 0.2097 | 0.1857 | 0.5961 |
| k__Bacteria;p__Proteobacteria | 0.0073 | 0.2371 | 0.7089 | 0.6596 | NA | 0.6001 | 0.0215 | 0.0231 | 0.0066 | 0.5112 | 0.4041 | 0.4106 | 0.4968 | 0.3600 | 0.6443 |
| k__Bacteria;p__Verrucomicrobiota | 0.2927 | 0.0199 | 0.7379 | 0.0659 | 0.0111 | 0.0950 | 0.1031 | 0.2557 | 0.0112 | 0.0144 | 0.3502 | 0.5380 | 0.0476 | 0.0135 | 0.0969 |

Highlighted cells Significant at P<0.05
