## Supplemental Table 3 for "Addition of soluble fiber in low fat purified diets improves gut and metabolic health compared to traditional AIN diets"

### Supplementary Table 3: P-values for individual microbial taxa differences (pairwise comparisons, Tukey HSD) at the genus level in the cecum samples.

| Taxon | 5001__5002 | 5001__200CEL | 5001__200IN | 5001__200FOS | 5001__100CEL | 5001__100IN | 5001__100FOS | 5002__200CEL | 5002__200IN | 5002__200FOS | 5002__200CEL | 5002__100IN | 5002__100FOS | 200CEL__200IN | 200CEL__200FOS | 200CEL__100CEL | 200CEL__100IN | 200CEL__100FOS | 200IN__200FOS | 200IN__100CEL | 200IN__100IN | 200IN__100FOS | 200FOS__100CEL | 200FOS__100IN | 200FOS__100FOS | 100CEL__100IN | 100CEL__100FOS | 100IN__100FOS |
|---|---|---|---|---|---|---|---|---|---|---|---|---|---|---|---|---|---|---|---|---|---|---|---|---|---|---|---|---|
| k__Bacteria;p__Actinobacteriota;c__Actinobacteria;o__Bifidobacteriales;f__Bifidobacteriaceae;g__Bifidobacterium | 0.7249 | 0.7114 | 0.0093 | 0.0088 | 0.4637 | 0.0082 | 0.0082 | 0.7430 | 0.0100 | 0.0083 | 0.8471 | 0.0090 | 0.0077 | 0.0106 | 0.0104 | 0.8466 | 0.0102 | 0.0117 | 0.4408 | 0.0103 | 0.8985 | 0.7518 | 0.0098 | 0.0584 | 0.3001 | 0.0109 | 0.0121 | 0.1382 |
| k__Bacteria;p__Actinobacteriota;c__Actinobacteria;o__Bifidobacteriales;f__Bifidobacteriaceae;g__Other | NA | NA | NA | 0.2241 | NA | NA | 0.4260 | NA | NA | 0.1758 | NA | NA | 0.4696 | NA | 0.1128 | NA | NA | 0.3561 | 0.4305 | NA | NA | 0.7745 | 0.1604 | 0.5508 | 0.6035 | NA | 0.4222 | 0.8225 |
| k__Bacteria;p__Actinobacteriota;c__Coriobacteriia;o__Coriobacteriales;f__Atopobiaceae;g__Coriobacteriaceae UCG-002 | NA | NA | 0.2202 | NA | NA | NA | NA | NA | 0.2996 | NA | NA | NA | NA | 0.1708 | NA | NA | NA | NA | 0.5623 | 0.1547 | 0.6691 | 0.4242 | NA | NA | NA | NA | NA | NA |
| k__Bacteria;p__Actinobacteriota;c__Coriobacteriia;o__Coriobacteriales;f__Eggerthellaceae;g__Enterorhabdus | 0.9561 | 0.5381 | 0.1033 | 0.4611 | 0.8571 | 0.4137 | 0.4660 | 0.7868 | 0.2204 | 0.6433 | 0.8722 | 0.7179 | 0.5952 | 0.1522 | 0.6408 | 0.6935 | 0.7598 | 0.7121 | 0.6064 | 0.0254 | 0.7174 | 0.6790 | 0.3479 | 0.8919 | 0.7784 | 0.3738 | 0.3105 | 0.8688 |
| k__Bacteria;p__Actinobacteriota;c__Coriobacteriia;o__Coriobacteriales;f__Eggerthellaceae;g__Other | NA | 0.6403 | NA | NA | NA | NA | NA | 0.7251 | NA | NA | NA | NA | NA | 0.6939 | 0.5510 | 0.8067 | 0.6094 | 0.6877 | NA | NA | NA | NA | NA | NA | NA | NA | NA | NA |
| k__Bacteria;p__Bacteroidota;c__Bacteroidia;o__Bacteroidales;f__Bacteroidaceae;g__Bacteroides | 0.8647 | 0.0092 | 0.0093 | 0.0088 | 0.0317 | 0.0082 | 0.0082 | 0.0101 | 0.0100 | 0.0083 | 0.1266 | 0.0090 | 0.0077 | 0.2154 | 0.0901 | 0.2483 | 0.0106 | 0.3234 | 0.8493 | 0.0147 | 0.8624 | 0.7423 | 0.0099 | 0.9812 | 0.2868 | 0.0109 | 0.0123 | 0.0292 |
| k__Bacteria;p__Bacteroidota;c__Bacteroidia;o__Bacteroidales;f__Muribaculaceae;g__Muribaculum | 0.4382 | 1.0000 | 0.0130 | 0.1055 | 0.5639 | 0.9536 | 0.2667 | 0.9933 | 0.4518 | 0.0303 | 0.0851 | 0.7247 | 0.7011 | 0.9713 | 0.7225 | 0.8381 | 0.9163 | 0.6680 | 0.1145 | 0.0135 | 0.7363 | 0.9600 | 0.8825 | 0.7321 | 0.2162 | 0.6065 | 0.1486 | 0.6836 |
| k__Bacteria;p__Bacteroidota;c__Bacteroidia;o__Bacteroidales;f__Muribaculaceae;g__Other | 0.9607 | 0.5085 | 0.6746 | 0.0444 | 0.6524 | 0.0308 | 0.0372 | 0.5825 | 0.6580 | 0.0621 | 0.7993 | 0.0479 | 0.1344 | 0.3906 | 0.0243 | 0.5296 | 0.0136 | 0.0119 | 0.7264 | 0.8611 | 0.7270 | 0.8885 | 0.0597 | 0.8569 | 0.3833 | 0.0305 | 0.0202 | 0.5092 |
| k__Bacteria;p__Bacteroidota;c__Bacteroidia;o__Bacteroidales;f__Other;g__Other | 0.8423 | 0.4570 | 0.7167 | 0.4149 | 0.7600 | 0.6707 | 0.7006 | 0.8443 | 0.6558 | 0.6840 | 0.8179 | 0.7503 | 0.7270 | 0.3565 | 0.8652 | 0.8934 | 0.4832 | 0.5883 | 0.6103 | 0.7439 | NA | 0.8460 | 0.6677 | 0.6709 | 0.5601 | 0.8192 | 0.8340 | 0.7680 |
| k__Bacteria;p__Bacteroidota;c__Bacteroidia;o__Bacteroidales;f__Prevotellaceae;g__Other | 0.8633 | 0.4617 | 0.6781 | 0.6254 | 0.6222 | 0.6403 | 0.6535 | NA | NA | NA | NA | NA | NA | NA | NA | NA | NA | NA | NA | NA | NA | NA | NA | NA | NA | NA | NA | NA |
| k__Bacteria;p__Bacteroidota;c__Bacteroidia;o__Bacteroidales;f__Prevotellaceae;g__Prevotellaceae NK3B31 group | 0.9582 | 0.2291 | 0.4738 | 0.5969 | 0.3240 | 0.6783 | 0.3950 | 0.2297 | 0.4151 | 0.5313 | 0.2592 | 0.6322 | 0.3272 | NA | NA | NA | 0.2496 | NA | NA | NA | 0.7802 | NA | NA | 0.7974 | NA | 0.3439 | NA | 0.5199 |
| k__Bacteria;p__Bacteroidota;c__Bacteroidia;o__Bacteroidales;f__Prevotellaceae;g__Prevotellaceae UCG-001 | 0.6436 | 0.0092 | 0.0093 | 0.0088 | 0.0111 | 0.0082 | 0.0082 | 0.0090 | 0.0100 | 0.0083 | 0.0113 | 0.0090 | 0.0077 | 0.7542 | 0.5258 | 0.8576 | 0.5516 | 0.7536 | 0.7598 | 0.6380 | 0.8051 | 0.7613 | 0.4300 | 0.8561 | 0.4507 | 0.3403 | 0.6620 | 0.4245 |
| k__Bacteria;p__Bacteroidota;c__Bacteroidia;o__Bacteroidales;f__Rikenellaceae;g__Alistipes | 0.6716 | 0.0349 | 0.8795 | 0.5136 | 0.9071 | 0.0989 | 0.9073 | 0.0090 | 0.7162 | 0.8930 | 0.7580 | 0.3373 | 0.4461 | 0.0109 | 0.0128 | 0.1692 | 0.0102 | 0.0174 | 0.8623 | 0.8700 | 0.7146 | 0.9413 | 0.8451 | 0.5878 | 0.7897 | 0.3771 | 0.8978 | 0.4009 |
| k__Bacteria;p__Bacteroidota;c__Bacteroidia;o__Bacteroidales;f__Tannerellaceae;g__Other | 0.9717 | 0.0092 | 0.0093 | 0.0088 | 0.0111 | 0.0082 | 0.0082 | 0.0090 | 0.0100 | 0.0083 | 0.0113 | 0.0090 | 0.0077 | 0.5455 | 0.7480 | 0.9352 | 0.2960 | 0.8243 | 0.6827 | 0.1523 | 0.9175 | 0.8202 | 0.8172 | 0.4203 | 0.5722 | 0.0159 | 0.2580 | 0.7143 |
| k__Bacteria;p__Bacteroidota;c__Bacteroidia;o__Bacteroidales;f__Tannerellaceae;g__Parabacteroides | NA | NA | 0.5272 | NA | NA | NA | NA | NA | 0.5333 | NA | NA | NA | NA | 0.3656 | NA | NA | NA | NA | 0.8294 | 0.4452 | 0.8669 | 0.7382 | NA | NA | NA | NA | NA | NA |
| k__Bacteria;p__Bacteroidota;c__Bacteroidia;o__Other;f__Other;g__Other | NA | 0.6962 | NA | NA | 0.7284 | NA | NA | 0.7434 | NA | NA | 0.7270 | NA | NA | 0.6598 | 0.5370 | 0.8474 | 0.5377 | 0.6677 | NA | 0.6837 | NA | NA | 0.5903 | NA | NA | 0.6573 | 0.7374 | NA |
| k__Bacteria;p__Bacteroidota;c__Other;o__Other;f__Other;g__Other | NA | 0.6700 | NA | NA | NA | NA | NA | 0.7213 | NA | NA | NA | NA | NA | 0.6580 | 0.5975 | 0.8405 | 0.5690 | 0.7104 | NA | NA | NA | NA | NA | NA | NA | NA | NA | NA |
| k__Bacteria;p__Cyanobacteria;c__Vampirivibrionia;o__Gastranaerophilales;f__Other;g__Other | 0.7947 | NA | 0.1329 | NA | NA | 0.0082 | 0.5789 | 0.5431 | 0.3426 | 0.6921 | 0.5582 | 0.0503 | 0.6871 | 0.1131 | NA | NA | 0.0102 | 0.3792 | 0.4721 | 0.1187 | 0.8740 | 0.4400 | NA | 0.0519 | 0.6606 | 0.0109 | 0.4576 | 0.0280 |
| k__Bacteria;p__Deferribacterota;c__Deferribacteres;o__Deferribacterales;f__Deferribacteraceae;g__Mucispirillum | 0.7032 | 0.0098 | 0.9113 | 0.2668 | 0.0111 | 0.0133 | 0.9387 | 0.0093 | 0.4356 | 0.0551 | 0.0113 | 0.0095 | 0.3892 | 0.0133 | 0.3880 | 0.9342 | 0.8077 | 0.0592 | 0.5363 | 0.0103 | 0.2656 | 0.9811 | 0.4961 | 0.7805 | 0.3343 | 0.7595 | 0.0515 | 0.1602 |
| k__Bacteria;p__Firmicutes;c__Bacilli;o__Acholeplasmatales;f__Acholeplasmataceae;g__Anaeroplasma | 0.9553 | 0.0092 | 0.0093 | 0.0088 | 0.0569 | 0.0082 | 0.0082 | 0.0090 | 0.0100 | 0.0083 | 0.0400 | 0.0090 | 0.0077 | 0.7474 | 0.7789 | 0.8930 | 0.8153 | 0.8255 | NA | 0.5653 | NA | 0.7829 | 0.7493 | NA | 0.7425 | 0.6771 | 0.8523 | 0.8089 |
| k__Bacteria;p__Firmicutes;c__Bacilli;o__Erysipelotrichales;f__Erysipelatoclostridiaceae;g__Candidatus Stoquefichus | 0.7334 | 0.0652 | NA | NA | 0.6785 | NA | NA | 0.1710 | 0.6734 | 0.6823 | 0.7243 | 0.7491 | 0.6358 | 0.1047 | 0.1293 | 0.4886 | 0.1397 | 0.0839 | NA | 0.6984 | NA | NA | 0.6741 | NA | NA | 0.6652 | 0.7279 | NA |
| k__Bacteria;p__Firmicutes;c__Bacilli;o__Erysipelotrichales;f__Erysipelatoclostridiaceae;g__Erysipelatoclostridium | NA | NA | 0.1746 | NA | NA | NA | NA | NA | 0.2356 | NA | NA | NA | NA | 0.1698 | NA | NA | NA | NA | 0.5711 | 0.1598 | 0.6621 | 0.4585 | NA | NA | NA | NA | NA | NA |
| k__Bacteria;p__Firmicutes;c__Bacilli;o__Erysipelotrichales;f__Erysipelotrichaceae;g__[Clostridium] innocuum group | NA | 0.4532 | NA | NA | 0.6820 | NA | NA | 0.5857 | NA | NA | 0.7065 | NA | NA | 0.5978 | 0.7249 | 0.7594 | 0.7258 | 0.5849 | NA | 0.7337 | NA | NA | 0.6081 | NA | NA | 0.6374 | 0.6978 | NA |
| k__Bacteria;p__Firmicutes;c__Bacilli;o__Erysipelotrichales;f__Erysipelotrichaceae;g__Allobaculum | NA | NA | NA | NA | NA | 0.2721 | NA | NA | NA | NA | NA | 0.3118 | NA | NA | NA | NA | 0.2271 | NA | NA | NA | 0.7245 | NA | NA | 0.6989 | NA | 0.2342 | NA | 0.4161 |
| k__Bacteria;p__Firmicutes;c__Bacilli;o__Erysipelotrichales;f__Erysipelotrichaceae;g__Dubosiella | NA | NA | 0.0336 | 0.3623 | 0.7000 | 0.0706 | 0.1302 | NA | 0.0488 | 0.3809 | 0.7789 | 0.0984 | 0.1270 | 0.0286 | 0.2643 | 0.7565 | 0.0646 | 0.1005 | 0.2141 | 0.0371 | 0.9716 | 0.9746 | 0.5128 | 0.5329 | 0.4596 | 0.1182 | 0.1618 | 0.9493 |
| k__Bacteria;p__Firmicutes;c__Bacilli;o__Erysipelotrichales;f__Erysipelotrichaceae;g__Faecalibaculum | 0.7266 | 0.6811 | 0.0093 | 0.0088 | 0.1983 | 0.0082 | 0.0082 | 0.7675 | 0.0100 | 0.0083 | 0.8007 | 0.0090 | 0.0077 | 0.0105 | 0.0104 | 0.6892 | 0.0102 | 0.0117 | 0.3569 | 0.0104 | 0.7283 | 0.6433 | 0.0098 | 0.6731 | 0.2574 | 0.0109 | 0.0121 | 0.7910 |
| k__Bacteria;p__Firmicutes;c__Bacilli;o__Erysipelotrichales;f__Erysipelotrichaceae;g__Faecalitalea | NA | NA | 0.5343 | 0.2886 | NA | 0.1474 | NA | NA | 0.5839 | 0.3337 | NA | 0.1722 | NA | 0.4205 | 0.1869 | NA | 0.0880 | NA | 0.7048 | 0.4979 | 0.7788 | 0.8085 | 0.2704 | 0.9293 | 0.4232 | 0.1523 | NA | 0.3211 |
| k__Bacteria;p__Firmicutes;c__Bacilli;o__Erysipelotrichales;f__Erysipelotrichaceae;g__Other | 0.3572 | 0.1344 | 0.1301 | 0.9302 | 0.7609 | 0.6704 | 0.3903 | 0.6731 | NA | 0.2166 | 0.0829 | 0.3288 | 0.0079 | 0.6758 | 0.4348 | 0.3430 | 0.6448 | 0.0123 | 0.5117 | 0.0991 | 0.7411 | 0.0733 | 0.9317 | 0.8321 | 0.3774 | 0.7830 | 0.2063 | 0.2784 |
| k__Bacteria;p__Firmicutes;c__Bacilli;o__Erysipelotrichales;f__Erysipelotrichaceae;g__Turicibacter | NA | 0.6447 | 0.3591 | NA | 0.4414 | NA | NA | 0.7593 | 0.4224 | NA | 0.5238 | NA | NA | 0.5450 | 0.7205 | 0.7706 | 0.7543 | 0.7534 | 0.7463 | 0.8525 | 0.8251 | 0.6371 | 0.7677 | NA | NA | 0.7098 | 0.5572 | NA |
| k__Bacteria;p__Firmicutes;c__Bacilli;o__Lactobacillales;f__Enterococcaceae;g__Enterococcus | NA | 0.7009 | 0.5553 | 0.0497 | NA | NA | NA | 0.7478 | 0.5986 | 0.0425 | NA | NA | NA | 0.5567 | 0.0442 | 0.8421 | 0.5715 | 0.7222 | 0.4626 | 0.4561 | 0.8573 | 0.7790 | 0.0418 | 0.2080 | 0.1473 | NA | NA | NA |
| k__Bacteria;p__Firmicutes;c__Bacilli;o__Lactobacillales;f__Lactobacillaceae;g__Lactobacillus | 0.1988 | 0.0234 | 0.4444 | 0.7831 | 0.1231 | 0.8663 | 0.5193 | 0.5870 | 0.0531 | 0.0130 | 0.2420 | 0.0235 | 0.0248 | 0.1241 | 0.0178 | 0.7947 | 0.0562 | 0.0514 | 0.3837 | 0.4114 | 0.7130 | 0.6254 | 0.0751 | 0.9661 | 0.7417 | 0.1617 | 0.1113 | 0.7551 |
| k__Bacteria;p__Firmicutes;c__Bacilli;o__Lactobacillales;f__Other;g__Other | NA | NA | NA | NA | NA | NA | 0.5212 | NA | NA | NA | NA | NA | 0.5847 | NA | NA | NA | NA | 0.4073 | NA | NA | NA | 0.8226 | NA | NA | 0.6841 | NA | 0.5472 | 0.8034 |
| k__Bacteria;p__Firmicutes;c__Bacilli;o__Lactobacillales;f__Streptococcaceae;g__Lactococcus | NA | 0.0092 | 0.0093 | 0.0088 | 0.0111 | 0.0082 | 0.0082 | 0.0090 | 0.0100 | 0.0083 | 0.0113 | 0.0097 | 0.0077 | 0.6760 | 0.6999 | 0.8873 | 0.3154 | 0.5598 | 0.6992 | 0.6307 | 0.8540 | 0.9231 | 0.7258 | 0.3311 | 0.2061 | 0.2813 | 0.3842 | 0.8984 |
| k__Bacteria;p__Firmicutes;c__Bacilli;o__RF39;f__Other;g__Other | 0.9304 | 0.9379 | 0.9080 | 0.3563 | 0.3620 | 0.8923 | 0.9797 | 0.8876 | 0.7330 | 0.1919 | 0.6890 | 0.9424 | 0.8228 | 0.9472 | 0.2633 | 0.6702 | 0.9598 | 0.9845 | 0.5130 | 0.3788 | 0.9487 | 0.9750 | 0.1653 | 0.6661 | 0.4476 | 0.7176 | 0.6168 | 0.8880 |
| k__Bacteria;p__Firmicutes;c__Bacilli;o__Staphylococcales;f__Staphylococcaceae;g__Staphylococcus | NA | 0.6786 | NA | NA | 0.6837 | NA | NA | 0.7666 | NA | NA | 0.6817 | NA | NA | 0.7680 | 0.7392 | 0.8639 | 0.7747 | 0.7989 | NA | 0.6050 | NA | NA | 0.5171 | NA | NA | 0.5486 | 0.6915 | NA |
| k__Bacteria;p__Firmicutes;c__Clostridia;o__Christensenellales;f__Christensenellaceae;g__Other | NA | 0.7007 | NA | NA | NA | NA | NA | 0.7460 | NA | NA | NA | NA | NA | 0.6822 | 0.5945 | 0.8182 | 0.6462 | 0.7030 | NA | NA | NA | NA | NA | NA | NA | NA | NA | NA |
| k__Bacteria;p__Firmicutes;c__Clostridia;o__Clostridia UCG-014;f__Other;g__Other | 0.9129 | 0.4759 | 0.8261 | 0.1506 | 0.1030 | 0.3975 | 0.5505 | 0.3571 | 0.9001 | 0.0484 | 0.0418 | 0.5202 | 0.9025 | 0.6603 | 0.3181 | 0.5161 | 0.3337 | 0.4794 | 0.5614 | 0.1993 | 0.7516 | 0.9659 | 0.6615 | 0.3079 | 0.4466 | 0.0802 | 0.2142 | 0.5773 |
| k__Bacteria;p__Firmicutes;c__Clostridia;o__Clostridia vadinBB60 group;f__Other;g__Other | 0.9638 | 0.0379 | 0.0093 | 0.0957 | 0.0311 | 0.0094 | 0.0082 | 0.1070 | 0.0145 | 0.0869 | 0.0762 | 0.0258 | 0.0102 | 0.3199 | 0.5279 | 0.9496 | 0.4276 | 0.3988 | 0.9211 | 0.2122 | 0.9651 | 0.9650 | 0.4446 | 0.9605 | 0.8548 | 0.3380 | 0.2425 | 0.9175 |
| k__Bacteria;p__Firmicutes;c__Clostridia;o__Clostridiales;f__Clostridiaceae;g__Candidatus Arthromitus | 0.9674 | 0.1033 | 0.2375 | 0.3036 | 0.1434 | 0.3033 | 0.1745 | 0.0866 | 0.2281 | 0.2663 | 0.1181 | 0.2700 | 0.1372 | NA | NA | NA | NA | NA | NA | NA | NA | NA | NA | NA | NA | NA | NA | NA |
| k__Bacteria;p__Firmicutes;c__Clostridia;o__Clostridiales;f__Clostridiaceae;g__Clostridium sensu stricto 1 | 0.8597 | 0.0390 | 0.3686 | 0.6613 | 0.0181 | 0.6905 | 0.6879 | 0.0090 | 0.2578 | NA | 0.0113 | NA | NA | 0.3408 | 0.0136 | 0.3545 | 0.0111 | 0.0117 | 0.5945 | 0.0464 | 0.6746 | 0.4689 | 0.0109 | NA | NA | 0.0116 | 0.0122 | NA |
| k__Bacteria;p__Firmicutes;c__Clostridia;o__Lachnospirales;f__Lachnospiraceae;g__[Eubacterium] fissicatena group | NA | NA | NA | NA | 0.4552 | NA | NA | NA | NA | NA | 0.5039 | NA | NA | NA | NA | 0.5326 | NA | NA | NA | 0.5363 | NA | NA | 0.6920 | NA | NA | 0.6616 | 0.5306 | NA |
| k__Bacteria;p__Firmicutes;c__Clostridia;o__Lachnospirales;f__Lachnospiraceae;g__[Eubacterium] oxidoreducens group | 0.8500 | 0.6830 | 0.7069 | 0.6553 | 0.6161 | 0.6635 | 0.7077 | 0.7354 | NA | NA | NA | NA | NA | 0.6829 | 0.4774 | 0.8102 | 0.5594 | 0.7144 | NA | NA | NA | NA | NA | NA | NA | NA | NA | NA |
| k__Bacteria;p__Firmicutes;c__Clostridia;o__Lachnospirales;f__Lachnospiraceae;g__[Eubacterium] xylanophilum group | 0.8431 | 0.0092 | 0.0093 | 0.0088 | 0.0111 | 0.0082 | 0.0082 | 0.0090 | 0.0100 | 0.0083 | 0.0113 | 0.0090 | 0.0077 | NA | NA | NA | NA | NA | NA | NA | NA | NA | NA | NA | NA | NA | NA | NA |
| k__Bacteria;p__Firmicutes;c__Clostridia;o__Lachnospirales;f__Lachnospiraceae;g__[Ruminococcus] torques group | NA | NA | NA | NA | NA | NA | 0.5994 | NA | NA | NA | NA | NA | 0.5695 | NA | NA | NA | NA | 0.4344 | NA | NA | NA | 0.7974 | NA | NA | 0.7336 | NA | 0.4985 | 0.7970 |
| k__Bacteria;p__Firmicutes;c__Clostridia;o__Lachnospirales;f__Lachnospiraceae;g__A2 | 0.9339 | 0.8235 | 0.0269 | 0.0088 | 0.9104 | 0.0082 | 0.1489 | 0.7385 | 0.0260 | 0.0083 | 0.8217 | 0.0090 | 0.1216 | 0.0374 | 0.0104 | 0.8711 | 0.0102 | 0.1451 | 0.7402 | 0.0225 | 0.8757 | 0.2937 | 0.0098 | 0.6303 | 0.2058 | 0.0109 | 0.1509 | 0.0289 |
| k__Bacteria;p__Firmicutes;c__Clostridia;o__Lachnospirales;f__Lachnospiraceae;g__Acetatifactor | 0.5217 | 0.0092 | 0.2173 | 0.0798 | 0.0137 | 0.0732 | 0.3171 | 0.0090 | 0.0795 | 0.0566 | 0.0581 | 0.0644 | 0.1100 | 0.0105 | 0.0104 | 0.3519 | 0.0102 | 0.0117 | 0.8149 | 0.0104 | 0.8594 | 0.8531 | 0.0098 | NA | 0.7620 | 0.0109 | 0.0148 | 0.7925 |
| k__Bacteria;p__Firmicutes;c__Clostridia;o__Lachnospirales;f__Lachnospiraceae;g__Anaerostipes | 0.8092 | 0.2795 | 0.0539 | 0.0088 | 0.4663 | 0.3457 | 0.0082 | 0.6474 | 0.0652 | 0.0083 | 0.6603 | 0.5084 | 0.0077 | 0.0830 | 0.0104 | 0.9341 | 0.7756 | 0.0117 | 0.1504 | 0.0799 | 0.4257 | 0.7902 | 0.0098 | 0.0512 | 0.1423 | 0.8459 | 0.0158 | 0.0368 |
| k__Bacteria;p__Firmicutes;c__Clostridia;o__Lachnospirales;f__Lachnospiraceae;g__ASF356 | 0.9551 | 0.8406 | 0.1110 | 0.8204 | 0.9387 | 0.0091 | 0.0082 | 0.8969 | 0.1137 | 0.7743 | 0.9483 | 0.0132 | 0.0081 | 0.1254 | 0.7666 | 0.9503 | 0.0260 | 0.0121 | 0.9284 | 0.1384 | 0.8865 | 0.7785 | 0.7367 | 0.9821 | 0.7370 | 0.0455 | 0.0149 | 0.4997 |
| k__Bacteria;p__Firmicutes;c__Clostridia;o__Lachnospirales;f__Lachnospiraceae;g__Blautia | NA | NA | 0.4401 | 0.1227 | NA | 0.3850 | NA | NA | 0.4661 | 0.1077 | NA | 0.3790 | NA | 0.3356 | 0.0751 | NA | 0.2434 | NA | 0.7291 | 0.4141 | 0.9081 | 0.7182 | 0.0937 | 0.4739 | 0.2201 | 0.2934 | NA | 0.5762 |
| k__Bacteria;p__Firmicutes;c__Clostridia;o__Lachnospirales;f__Lachnospiraceae;g__Frisingicoccus | NA | NA | 0.5370 | NA | NA | NA | NA | NA | 0.5513 | NA | NA | NA | NA | 0.4033 | NA | NA | NA | NA | 0.8106 | 0.4856 | 0.8586 | 0.7041 | NA | NA | NA | NA | NA | NA |
| k__Bacteria;p__Firmicutes;c__Clostridia;o__Lachnospirales;f__Lachnospiraceae;g__GCA-900066575 | 0.9584 | 0.1486 | 0.5675 | 0.0994 | 0.3912 | 0.6085 | 0.6120 | 0.3111 | 0.6021 | 0.1078 | 0.5798 | 0.6757 | 0.5615 | 0.0848 | 0.0238 | 0.7316 | 0.1680 | 0.9162 | 0.5722 | 0.2445 | 0.9688 | 0.5707 | 0.0494 | 0.6251 | 0.1671 | 0.3939 | 0.8718 | 0.5603 |
| k__Bacteria;p__Firmicutes;c__Clostridia;o__Lachnospirales;f__Lachnospiraceae;g__Lachnoclostridium | 0.3200 | 0.1501 | 0.6099 | 0.6804 | 0.0387 | 0.4191 | 0.0163 | 0.7795 | 0.5676 | 0.8258 | 0.7137 | 0.7862 | 0.1318 | 0.5306 | 0.6905 | 0.7266 | 0.9048 | 0.1139 | 0.9070 | 0.5113 | 0.7685 | 0.5483 | 0.5552 | 0.8085 | 0.2864 | 0.9018 | 0.2155 | 0.7465 |
| k__Bacteria;p__Firmicutes;c__Clostridia;o__Lachnospirales;f__Lachnospiraceae;g__Lachnospiraceae FCS020 group | 0.7137 | 0.1726 | 0.1458 | 0.4083 | 0.3983 | 0.5740 | 0.4178 | 0.9469 | 0.8386 | 0.5443 | 0.9582 | 0.7989 | 0.9372 | 0.8434 | 0.5577 | 0.7612 | 0.3516 | 0.9388 | 0.7964 | 0.5690 | 0.7242 | 0.9488 | 0.5913 | 0.8402 | 0.7468 | 0.3361 | 0.9796 | 0.9010 |
| k__Bacteria;p__Firmicutes;c__Clostridia;o__Lachnospirales;f__Lachnospiraceae;g__Lachnospiraceae NK4A136 group | 0.9399 | 0.0195 | 0.0624 | 0.0088 | 0.0118 | 0.5260 | 0.0102 | 0.0386 | 0.0932 | 0.0084 | 0.0153 | 0.5781 | 0.0118 | 0.4770 | 0.1124 | 0.9173 | 0.8529 | 0.9338 | 0.7835 | 0.6051 | 0.9405 | 0.6992 | 0.1367 | 0.6707 | 0.1869 | 0.8981 | 0.9216 | 0.9646 |
| k__Bacteria;p__Firmicutes;c__Clostridia;o__Lachnospirales;f__Lachnospiraceae;g__Lachnospiraceae NK4B4 group | 0.8254 | 0.5234 | 0.7253 | 0.6432 | 0.6075 | 0.6582 | 0.7112 | NA | NA | NA | NA | NA | NA | NA | NA | NA | NA | NA | NA | NA | NA | NA | NA | NA | NA | NA | NA | NA |
| k__Bacteria;p__Firmicutes;c__Clostridia;o__Lachnospirales;f__Lachnospiraceae;g__Lachnospiraceae UCG-001 | 0.9653 | 0.0100 | 0.7635 | 0.0153 | 0.0111 | 0.1227 | 0.0084 | 0.0531 | 0.8582 | 0.0683 | 0.0552 | 0.3497 | 0.0515 | 0.2531 | 0.4581 | 0.8345 | 0.4289 | 0.7340 | 0.6039 | 0.2217 | 0.8050 | 0.4828 | NA | 0.7912 | NA | 0.3542 | NA | 0.5761 |
| k__Bacteria;p__Firmicutes;c__Clostridia;o__Lachnospirales;f__Lachnospiraceae;g__Lachnospiraceae UCG-004 | 0.6935 | 0.3727 | 0.3335 | 0.3888 | 0.6646 | 0.4062 | 0.2306 | 0.8317 | 0.7813 | 0.9126 | 0.8877 | 0.8998 | 0.5093 | 0.7058 | 0.9147 | 0.7911 | 0.9439 | 0.8397 | 0.8285 | 0.5797 | 0.8925 | 0.9272 | 0.7998 | 0.9071 | 0.8603 | 0.8194 | 0.3937 | 0.9379 |
| k__Bacteria;p__Firmicutes;c__Clostridia;o__Lachnospirales;f__Lachnospiraceae;g__Lachnospiraceae UCG-006 | 0.9607 | 0.0093 | 0.1343 | 0.4580 | 0.0129 | 0.0824 | 0.4722 | 0.0090 | 0.4712 | 0.8264 | 0.0158 | 0.4861 | 0.3475 | 0.0105 | 0.0105 | 0.7163 | 0.0102 | 0.2214 | 0.6888 | 0.0103 | 0.8641 | 0.2603 | 0.0120 | 0.6682 | 0.3336 | 0.0109 | 0.5634 | 0.1768 |
| k__Bacteria;p__Firmicutes;c__Clostridia;o__Lachnospirales;f__Lachnospiraceae;g__Lachnospiraceae UCG-010 | 0.7101 | 0.5136 | 0.0661 | 0.1010 | 0.1399 | 0.0914 | 0.0596 | 0.7415 | 0.3218 | 0.4509 | 0.5552 | 0.3934 | 0.2346 | 0.0354 | 0.0808 | 0.4189 | 0.0710 | 0.0274 | NA | 0.6229 | NA | NA | 0.7638 | NA | NA | 0.7453 | 0.5883 | NA |
| k__Bacteria;p__Firmicutes;c__Clostridia;o__Lachnospirales;f__Lachnospiraceae;g__Marvinbryantia | 0.8142 | 0.1393 | 0.8506 | 0.6319 | 0.1544 | 0.6827 | 0.3207 | 0.1181 | 0.6294 | 0.6709 | 0.1276 | 0.3704 | 0.1564 | 0.0124 | 0.0105 | 0.9092 | 0.5953 | 0.9221 | 0.7667 | 0.0134 | 0.8269 | 0.4070 | 0.0105 | 0.6490 | 0.2012 | 0.5877 | 0.9107 | 0.9270 |
| k__Bacteria;p__Firmicutes;c__Clostridia;o__Lachnospirales;f__Lachnospiraceae;g__Murimonas | NA | NA | NA | 0.2947 | NA | NA | NA | NA | NA | 0.3716 | NA | NA | NA | NA | 0.2255 | NA | NA | NA | 0.6581 | NA | NA | NA | 0.2919 | 0.7572 | 0.4603 | NA | NA | NA |
| k__Bacteria;p__Firmicutes;c__Clostridia;o__Lachnospirales;f__Lachnospiraceae;g__Other | 0.9301 | 0.6254 | 0.6718 | 0.3444 | 0.7870 | 0.0521 | 0.0259 | 0.7872 | 0.7239 | 0.3300 | 0.7845 | 0.0608 | 0.0441 | 0.8901 | 0.7054 | 0.7661 | 0.2874 | 0.1171 | 0.9238 | 0.6415 | 0.8387 | 0.7258 | 0.3031 | 0.8543 | 0.7510 | 0.0539 | 0.0281 | 0.8993 |
| k__Bacteria;p__Firmicutes;c__Clostridia;o__Lachnospirales;f__Lachnospiraceae;g__Roseburia | 0.9663 | 0.7339 | 0.0094 | 0.0091 | 0.0976 | 0.0230 | 0.0420 | 0.6967 | 0.0101 | 0.0086 | 0.0952 | 0.0153 | 0.0309 | 0.0105 | 0.0104 | 0.4251 | 0.0102 | 0.0184 | NA | 0.0103 | 0.8106 | 0.8181 | 0.0098 | 0.8235 | 0.7403 | 0.0109 | 0.0132 | 0.7642 |
| k__Bacteria;p__Firmicutes;c__Clostridia;o__Lachnospirales;f__Lachnospiraceae;g__Tuzzerella | NA | NA | NA | 0.4092 | 0.4418 | NA | NA | NA | NA | 0.4454 | 0.4874 | NA | NA | NA | 0.2850 | 0.5307 | NA | NA | 0.7085 | 0.5654 | NA | NA | 0.7175 | 0.7690 | 0.4761 | 0.6492 | 0.5128 | NA |
| k__Bacteria;p__Firmicutes;c__Clostridia;o__Lachnospirales;f__Lachnospiraceae;g__Tyzzerella | 0.8138 | 0.0092 | 0.0093 | 0.0089 | 0.0111 | 0.0083 | 0.0082 | 0.0090 | 0.0103 | 0.0089 | 0.0113 | 0.0091 | 0.0077 | NA | NA | NA | NA | NA | NA | NA | NA | NA | NA | NA | NA | NA | NA | NA |
| k__Bacteria;p__Firmicutes;c__Clostridia;o__Lachnospirales;f__Lachnospiraceae;g__UC5-1-2E3 | 0.7897 | NA | NA | NA | NA | NA | NA | 0.5145 | 0.7284 | 0.6588 | 0.5547 | 0.6650 | 0.6559 | NA | NA | NA | NA | NA | NA | NA | NA | NA | NA | NA | NA | NA | NA | NA |
| k__Bacteria;p__Firmicutes;c__Clostridia;o__Monoglobales;f__Monoglobaceae;g__Monoglobus | 0.8931 | 0.8276 | 0.6763 | 0.7238 | 0.8375 | 0.7395 | 0.7583 | 0.8231 | 0.6286 | 0.7097 | 0.7986 | 0.7647 | 0.7424 | 0.7517 | 0.7527 | 0.9014 | 0.7934 | 0.8393 | NA | 0.7102 | 0.7695 | 0.7797 | 0.7731 | 0.7750 | 0.7309 | 0.8039 | 0.7886 | 0.8380 |
| k__Bacteria;p__Firmicutes;c__Clostridia;o__Oscillospirales;f__[Eubacterium] coprostanoligenes group;g__Other | 0.4691 | 0.0092 | 0.3223 | 0.7737 | 0.0111 | 0.8837 | 0.0411 | 0.0090 | 0.1310 | 0.1986 | 0.0113 | 0.8345 | 0.3814 | 0.0105 | 0.0104 | 0.9053 | 0.0119 | 0.0337 | 0.7295 | 0.0103 | 0.7369 | 0.0868 | 0.0098 | 0.8385 | 0.1405 | 0.0113 | 0.0343 | 0.5926 |
| k__Bacteria;p__Firmicutes;c__Clostridia;o__Oscillospirales;f__Butyricicoccaceae;g__Butyricicoccus | 0.9549 | 0.0298 | 0.6584 | 0.0097 | 0.0939 | 0.6328 | 0.0360 | 0.0701 | 0.7916 | 0.0106 | 0.1240 | 0.7500 | 0.0361 | 0.3977 | 0.1476 | 0.7120 | 0.7058 | 0.3776 | 0.2013 | 0.6312 | 0.9499 | 0.2919 | 0.0174 | 0.3742 | 0.7195 | 0.8877 | 0.1523 | 0.3582 |
| k__Bacteria;p__Firmicutes;c__Clostridia;o__Oscillospirales;f__Oscillospiraceae;g__Colidextribacter | 0.8765 | 0.0122 | 0.5704 | 0.4645 | 0.0155 | 0.8820 | 0.5202 | 0.0090 | 0.7759 | 0.7267 | 0.0119 | 0.4342 | 0.6327 | 0.0119 | 0.0622 | 0.8180 | 0.0349 | 0.0201 | 0.9133 | 0.0144 | 0.7836 | 0.9473 | 0.1048 | 0.7888 | 0.8060 | 0.0665 | 0.0303 | 0.6948 |
| k__Bacteria;p__Firmicutes;c__Clostridia;o__Oscillospirales;f__Oscillospiraceae;g__Flavonifractor | NA | NA | 0.4276 | 0.1330 | NA | 0.1565 | 0.1596 | NA | 0.4586 | 0.1199 | NA | 0.1980 | 0.1471 | 0.3491 | 0.0739 | NA | 0.1221 | 0.1070 | 0.6773 | 0.3317 | 0.8699 | 0.8679 | 0.1159 | 0.8360 | 0.5284 | 0.1399 | 0.1203 | 0.9225 |
| k__Bacteria;p__Firmicutes;c__Clostridia;o__Oscillospirales;f__Oscillospiraceae;g__Intestinimonas | 0.9373 | 0.7602 | 0.0096 | 0.0146 | 0.3178 | 0.3814 | 0.0082 | 0.9293 | 0.0146 | 0.0246 | 0.2212 | 0.5381 | 0.0097 | 0.0145 | 0.0333 | 0.6338 | 0.5249 | 0.0141 | 0.7227 | 0.0103 | 0.6931 | 0.8103 | 0.0099 | 0.7826 | 0.7875 | 0.1524 | 0.0121 | 0.6144 |
| k__Bacteria;p__Firmicutes;c__Clostridia;o__Oscillospirales;f__Oscillospiraceae;g__Oscillibacter | 0.8878 | 0.3656 | 0.7825 | 0.7988 | 0.4237 | 0.0980 | 0.8836 | 0.7289 | 0.7444 | 0.8414 | 0.7627 | 0.2556 | 0.7448 | 0.4825 | 0.7752 | 0.9301 | 0.7863 | 0.5145 | 0.8279 | 0.5921 | 0.7441 | 0.9471 | 0.9227 | 0.8699 | 0.7730 | 0.6086 | 0.6325 | 0.2729 |
| k__Bacteria;p__Firmicutes;c__Clostridia;o__Oscillospirales;f__Oscillospiraceae;g__Other | 0.7267 | 0.0191 | 0.1226 | 0.3992 | 0.0179 | 0.7528 | 0.5433 | 0.0094 | 0.3811 | 0.8199 | 0.0118 | 0.8732 | 0.7121 | 0.0107 | 0.0145 | 0.9270 | 0.1279 | 0.0386 | 0.8802 | 0.0104 | 0.7596 | 0.9140 | 0.0169 | 0.8073 | 0.9004 | 0.1357 | 0.0431 | 0.8098 |
| k__Bacteria;p__Firmicutes;c__Clostridia;o__Oscillospirales;f__Oscillospiraceae;g__Pseudoflavonifractor | NA | NA | NA | NA | 0.7076 | NA | NA | NA | NA | NA | 0.7191 | NA | NA | NA | NA | 0.7439 | NA | NA | NA | 0.6893 | NA | NA | 0.6579 | NA | NA | 0.6221 | 0.7256 | NA |
| k__Bacteria;p__Firmicutes;c__Clostridia;o__Oscillospirales;f__Oscillospiraceae;g__UCG-003 | 0.8765 | 0.6960 | 0.7137 | 0.6241 | 0.6091 | 0.6922 | 0.6916 | 0.7496 | NA | NA | NA | NA | NA | 0.6455 | 0.5897 | 0.8412 | 0.5822 | 0.7294 | NA | NA | NA | NA | NA | NA | NA | NA | NA | NA |
| k__Bacteria;p__Firmicutes;c__Clostridia;o__Oscillospirales;f__Oscillospiraceae;g__UCG-005 | 0.2131 | 0.0294 | 0.0651 | 0.2509 | 0.1047 | 0.0092 | 0.2732 | 0.8552 | 0.6821 | 0.4368 | 0.5438 | 0.7369 | 0.0109 | 0.8449 | 0.7284 | 0.8185 | 0.7507 | 0.0135 | 0.7177 | 0.5512 | 0.8574 | 0.0990 | 0.8940 | 0.5782 | 0.2722 | 0.3524 | 0.0270 | 0.0279 |
| k__Bacteria;p__Firmicutes;c__Clostridia;o__Oscillospirales;f__Other;g__Other | NA | 0.6741 | NA | NA | 0.6750 | NA | NA | 0.7653 | NA | NA | 0.6992 | NA | NA | 0.7588 | 0.7122 | 0.8866 | 0.7374 | 0.7985 | NA | 0.6798 | NA | NA | 0.6218 | NA | NA | 0.6217 | 0.7240 | NA |
| k__Bacteria;p__Firmicutes;c__Clostridia;o__Oscillospirales;f__Ruminococcaceae;g__[Eubacterium] siraeum group | 0.9569 | 0.0092 | 0.0093 | 0.0088 | 0.0111 | 0.0082 | 0.0082 | 0.0090 | 0.0100 | 0.0083 | 0.0113 | 0.0091 | 0.0077 | NA | NA | NA | NA | NA | NA | NA | NA | NA | NA | NA | NA | NA | NA | NA |
| k__Bacteria;p__Firmicutes;c__Clostridia;o__Oscillospirales;f__Ruminococcaceae;g__Anaerotruncus | 0.7843 | 0.6033 | 0.1079 | 0.8654 | 0.6596 | 0.7725 | 0.2497 | 0.8624 | 0.0328 | 0.9274 | 0.9024 | 0.5372 | 0.1207 | 0.1008 | 0.6376 | 0.9008 | 0.5057 | 0.1699 | 0.5394 | 0.0693 | 0.7312 | 0.9221 | 0.9502 | 0.8875 | 0.5930 | 0.6729 | 0.2253 | 0.6474 |
| k__Bacteria;p__Firmicutes;c__Clostridia;o__Oscillospirales;f__Ruminococcaceae;g__Candidatus Soleaferrea | NA | 0.6479 | NA | NA | NA | NA | NA | 0.7625 | NA | NA | NA | NA | NA | 0.7869 | 0.7960 | 0.7884 | 0.7999 | 0.8166 | NA | NA | NA | NA | NA | NA | NA | NA | NA | NA |
| k__Bacteria;p__Firmicutes;c__Clostridia;o__Oscillospirales;f__Ruminococcaceae;g__Harryflintia | NA | 0.4288 | NA | NA | NA | NA | NA | 0.5416 | NA | NA | NA | NA | NA | 0.5903 | 0.6736 | 0.6236 | 0.6878 | 0.5887 | NA | NA | NA | NA | NA | NA | NA | NA | NA | NA |
| k__Bacteria;p__Firmicutes;c__Clostridia;o__Oscillospirales;f__Ruminococcaceae;g__Incertae Sedis | 0.5621 | 0.8914 | 0.0098 | 0.0127 | 0.8373 | 0.0084 | 0.0311 | 0.3447 | 0.0100 | 0.0083 | 0.1104 | 0.0090 | 0.0077 | 0.0110 | 0.0143 | 0.8384 | 0.0105 | 0.0426 | 0.8341 | 0.0140 | 0.8185 | 0.2970 | 0.0169 | 0.8249 | 0.1806 | 0.0120 | 0.0994 | 0.0895 |
| k__Bacteria;p__Firmicutes;c__Clostridia;o__Oscillospirales;f__Ruminococcaceae;g__Other | 0.2916 | 0.0206 | 0.8287 | 0.5887 | 0.0111 | 0.8144 | 0.6749 | 0.3114 | 0.3368 | 0.3496 | 0.0328 | 0.4832 | 0.0740 | 0.0624 | 0.0696 | 0.7395 | 0.0808 | 0.0283 | 0.8546 | 0.0194 | 0.9310 | 0.8770 | 0.0221 | 0.7604 | 0.9087 | 0.0347 | 0.0126 | 0.5777 |
| k__Bacteria;p__Firmicutes;c__Clostridia;o__Oscillospirales;f__Ruminococcaceae;g__Paludicola | NA | NA | NA | NA | 0.6794 | NA | NA | NA | NA | NA | 0.7064 | NA | NA | NA | NA | 0.7240 | NA | NA | NA | 0.6614 | NA | NA | 0.5991 | NA | NA | 0.5929 | 0.6989 | NA |
| k__Bacteria;p__Firmicutes;c__Clostridia;o__Oscillospirales;f__Ruminococcaceae;g__Ruminococcus | NA | NA | NA | NA | NA | NA | NA | NA | NA | NA | NA | NA | NA | NA | NA | NA | NA | NA | NA | NA | NA | NA | NA | NA | NA | NA | NA | NA |
| k__Bacteria;p__Firmicutes;c__Clostridia;o__Oscillospirales;f__UCG-010;g__Other | 0.8062 | 0.4179 | 0.6557 | 0.7410 | 0.4908 | 0.7541 | 0.6316 | NA | NA | NA | NA | NA | NA | NA | NA | NA | NA | NA | NA | NA | NA | NA | NA | NA | NA | NA | NA | NA |
| k__Bacteria;p__Firmicutes;c__Clostridia;o__Other;f__Other;g__Other | 0.8455 | 0.7063 | 0.6259 | 0.4864 | 0.7378 | 0.6148 | 0.6786 | 0.7393 | 0.5688 | 0.3956 | 0.7195 | NA | 0.5947 | 0.5331 | 0.3928 | 0.7852 | 0.5530 | 0.5956 | 0.7750 | 0.7943 | 0.8347 | 0.8582 | 0.7080 | 0.7673 | 0.7219 | 0.7710 | 0.7921 | 0.8123 |
| k__Bacteria;p__Firmicutes;c__Clostridia;o__Peptococcales;f__Peptococcaceae;g__Other | 0.9572 | 0.0093 | 0.3415 | 0.8012 | 0.6052 | 0.7079 | 0.5086 | 0.0090 | 0.3515 | 0.8344 | 0.6709 | 0.7308 | 0.3790 | 0.2597 | 0.1329 | 0.2380 | 0.9948 | 0.1488 | 0.6153 | 0.7404 | 0.9739 | 0.4989 | 0.8127 | 0.8109 | 0.5725 | 0.7085 | 0.5852 | 0.4800 |
| k__Bacteria;p__Firmicutes;c__Clostridia;o__Peptococcales;f__Peptococcaceae;g__Peptococcus | 0.9475 | 0.0094 | 0.8048 | 0.6777 | 0.0111 | 0.6538 | 0.7653 | 0.0100 | 0.8020 | 0.6967 | 0.0115 | 0.6538 | 0.7321 | 0.0124 | 0.0140 | 0.8239 | 0.0184 | 0.0182 | 0.8038 | 0.0110 | 0.8591 | 0.8710 | 0.0106 | 0.8507 | 0.7860 | 0.0145 | 0.0146 | 0.8436 |
| k__Bacteria;p__Firmicutes;c__Clostridia;o__Peptostreptococcales-Tissierellales;f__Anaerovoracaceae;g__[Eubacterium] brachy group | NA | NA | NA | NA | 0.4172 | NA | NA | NA | NA | NA | 0.4932 | NA | NA | NA | NA | 0.5626 | NA | NA | NA | 0.5338 | NA | NA | 0.7498 | NA | NA | 0.6891 | 0.5561 | NA |
| k__Bacteria;p__Firmicutes;c__Clostridia;o__Peptostreptococcales-Tissierellales;f__Anaerovoracaceae;g__[Eubacterium] nodatum group | NA | NA | NA | NA | 0.6984 | NA | NA | NA | NA | NA | 0.7224 | NA | NA | NA | NA | 0.7499 | NA | NA | NA | 0.6858 | NA | NA | 0.6392 | NA | NA | 0.6484 | 0.7435 | NA |
| k__Bacteria;p__Firmicutes;c__Clostridia;o__Peptostreptococcales-Tissierellales;f__Anaerovoracaceae;g__Family XIII AD3011 group | 0.6958 | NA | NA | NA | NA | NA | NA | 0.3215 | 0.6738 | 0.7177 | 0.3873 | 0.7101 | 0.5441 | NA | NA | NA | NA | NA | NA | NA | NA | NA | NA | NA | NA | NA | NA | NA |
| k__Bacteria;p__Firmicutes;c__Clostridia;o__Peptostreptococcales-Tissierellales;f__Peptostreptococcaceae;g__Clostridioides | NA | NA | 0.5538 | NA | NA | NA | NA | NA | 0.6021 | NA | NA | NA | NA | 0.4370 | NA | NA | NA | NA | 0.7879 | 0.5019 | 0.8732 | 0.7761 | NA | NA | NA | NA | NA | NA |
| k__Bacteria;p__Firmicutes;c__Clostridia;o__Peptostreptococcales-Tissierellales;f__Peptostreptococcaceae;g__Romboutsia | NA | 0.2291 | NA | NA | 0.0776 | NA | NA | 0.2922 | NA | NA | 0.0757 | NA | NA | 0.3334 | 0.4547 | 0.8284 | 0.4682 | 0.3121 | NA | 0.0975 | NA | NA | 0.1218 | NA | NA | 0.1176 | 0.0864 | NA |
| k__Bacteria;p__Firmicutes;c__Other;o__Other;f__Other;g__Other | 0.8483 | 0.5232 | 0.6538 | 0.6641 | 0.6112 | 0.6723 | 0.7110 | 0.4197 | 0.5909 | NA | NA | NA | NA | 0.6696 | 0.6165 | 0.5163 | 0.6543 | 0.4962 | 0.7903 | 0.4375 | 0.8476 | 0.6917 | NA | NA | NA | NA | NA | NA |
| k__Bacteria;p__Other;c__Other;o__Other;f__Other;g__Other | 0.8048 | 0.7278 | NA | 0.3704 | 0.4275 | 0.4303 | 0.5597 | 0.8076 | 0.7275 | 0.5937 | 0.6805 | 0.5855 | 0.7224 | 0.7744 | 0.6215 | 0.8181 | 0.6970 | 0.8131 | 0.6106 | 0.5906 | 0.7808 | 0.7949 | 0.8372 | 0.8394 | 0.5984 | 0.8387 | 0.7819 | 0.7365 |
| k__Bacteria;p__Proteobacteria;c__Alphaproteobacteria;o__Rhodospirillales;f__Other;g__Other | 0.7954 | 0.3655 | 0.7682 | 0.7509 | 0.5167 | 0.7366 | 0.5913 | NA | 0.6137 | NA | NA | NA | NA | 0.4190 | NA | NA | NA | NA | 0.7901 | 0.5030 | 0.8696 | 0.8052 | NA | NA | NA | NA | NA | NA |
| k__Bacteria;p__Proteobacteria;c__Gammaproteobacteria;o__Burkholderiales;f__Sutterellaceae;g__Parasutterella | NA | NA | NA | NA | NA | 0.4258 | NA | NA | NA | NA | NA | 0.4582 | NA | NA | NA | NA | 0.2625 | NA | NA | NA | 0.7914 | NA | NA | 0.7806 | NA | 0.3179 | NA | 0.6066 |
| k__Bacteria;p__Proteobacteria;c__Gammaproteobacteria;o__Enterobacterales;f__Enterobacteriaceae;g__Aquamonas | NA | NA | 0.5126 | NA | NA | NA | NA | NA | 0.5060 | NA | NA | NA | NA | 0.3429 | NA | NA | NA | NA | 0.7823 | 0.3980 | 0.8703 | 0.7278 | NA | NA | NA | NA | NA | NA |
| k__Bacteria;p__Proteobacteria;c__Gammaproteobacteria;o__Enterobacterales;f__Enterobacteriaceae;g__Escherichia-Shigella | NA | NA | 0.0093 | 0.0407 | 0.7542 | NA | NA | NA | 0.0100 | 0.0477 | 0.6967 | NA | NA | 0.0105 | 0.0346 | 0.7502 | NA | NA | 0.7551 | 0.0106 | 0.1290 | 0.0705 | 0.0681 | 0.2396 | 0.1467 | 0.6541 | 0.7393 | NA |
| k__Bacteria;p__Proteobacteria;c__Gammaproteobacteria;o__Enterobacterales;f__Enterobacteriaceae;g__Other | NA | NA | 0.4798 | NA | NA | NA | NA | NA | 0.5402 | NA | NA | NA | NA | 0.3875 | NA | NA | NA | NA | 0.7984 | 0.3894 | 0.8582 | 0.6887 | NA | NA | NA | NA | NA | NA |
| k__Bacteria;p__Proteobacteria;c__Gammaproteobacteria;o__Enterobacterales;f__Morganellaceae;g__Proteus | NA | NA | NA | 0.4078 | NA | NA | NA | NA | NA | 0.4567 | NA | NA | NA | NA | 0.2827 | NA | NA | NA | 0.6911 | NA | NA | NA | 0.3937 | 0.8052 | 0.5690 | NA | NA | NA |
| k__Bacteria;p__Proteobacteria;c__Gammaproteobacteria;o__Enterobacterales;f__Other;g__Other | NA | NA | NA | 0.3798 | NA | NA | NA | NA | NA | 0.4706 | NA | NA | NA | NA | 0.3020 | NA | NA | NA | 0.7268 | NA | NA | NA | 0.3214 | 0.7753 | 0.5086 | NA | NA | NA |
| k__Bacteria;p__Proteobacteria;c__Gammaproteobacteria;o__Pseudomonadales;f__Pseudomonadaceae;g__Pseudomonas | 0.8346 | 0.6822 | 0.6564 | 0.5634 | 0.5827 | 0.5499 | 0.6922 | 0.7420 | NA | NA | NA | NA | NA | 0.6296 | 0.5338 | 0.8260 | 0.5352 | 0.7116 | NA | NA | NA | NA | NA | NA | NA | NA | NA | NA |
| k__Bacteria;p__Verrucomicrobiota;c__Verrucomicrobiae;o__Verrucomicrobiales;f__Akkermansiaceae;g__Akkermansia | 0.7914 | 0.5226 | 0.0093 | 0.0088 | 0.7792 | 0.0082 | 0.0082 | 0.7807 | 0.0100 | 0.0083 | 0.4064 | 0.0090 | 0.0077 | 0.0105 | 0.0104 | 0.6640 | 0.0102 | 0.0117 | 0.1858 | 0.0103 | 0.8397 | 0.8761 | 0.0098 | 0.4232 | 0.1230 | 0.0109 | 0.0121 | 0.9091 |

Highlighted cells Significant at P<0.05

Comparison with baseline also included.
