## Supplemental Table 4 for "Addition of soluble fiber in low fat purified diets improves gut and metabolic health compared to traditional AIN diets"

### Supplementary Table 4: P-values for individual microbial taxa differences (pairwise comparisons, Tukey HSD) at the genus level in the colon samples.

| Taxon | 5001__5002 | 5001__200CEL | 5001__200IN | 5001__200FOS | 5001__100CEL | 5001__100IN | 5001__100FOS | 5002__200CEL | 5002__200IN | 5002__200FOS | 5002__200CEL | 5002__100IN | 5002__100FOS | 200CEL__200IN | 200CEL__200FOS | 200CEL__100CEL | 200CEL__100IN | 200CEL__100FOS | 200IN__200FOS | 200IN__100CEL | 200IN__100IN | 200IN__100FOS | 200FOS__100CEL | 200FOS__100IN | 200FOS__100FOS | 100CEL__100IN | 100CEL__100FOS | 100IN__100FOS |
|---|---|---|---|---|---|---|---|---|---|---|---|---|---|---|---|---|---|---|---|---|---|---|---|---|---|---|---|---|
| k__Bacteria;p__Actinobacteriota;c__Actinobacteria;o__Bifidobacteriales;f__Bifidobacteriaceae;g__Bifidobacterium | 0.81736 | NA | 0.03242 | 0.00962 | 0.67715 | 0.01050 | 0.00943 | 0.47614 | 0.03406 | 0.00867 | 0.70946 | 0.00933 | 0.00858 | 0.02565 | 0.01075 | 0.76208 | 0.01044 | 0.01472 | 0.36210 | 0.03339 | 0.87457 | 0.43600 | 0.01020 | 0.04698 | 0.59169 | 0.01048 | 0.01110 | 0.09350 |
| k__Bacteria;p__Actinobacteriota;c__Actinobacteria;o__Bifidobacteriales;f__Bifidobacteriaceae;g__Other | NA | NA | NA | 0.31170 | NA | NA | 0.47517 | NA | NA | 0.29304 | NA | NA | 0.44554 | NA | 0.22354 | NA | NA | 0.32162 | 0.71009 | NA | NA | 0.80444 | 0.31126 | 0.79453 | 0.75338 | NA | 0.40715 | 0.81064 |
| k__Bacteria;p__Actinobacteriota;c__Coriobacteriia;o__Coriobacteriales;f__Atopobiaceae;g__Coriobacteriaceae UCG-002 | NA | NA | 0.17125 | NA | NA | NA | NA | NA | 0.17351 | NA | NA | NA | NA | 0.13336 | NA | NA | NA | NA | 0.58678 | 0.16926 | 0.61172 | 0.42361 | NA | NA | NA | NA | NA | NA |
| k__Bacteria;p__Actinobacteriota;c__Coriobacteriia;o__Coriobacteriales;f__Eggerthellaceae;g__Enterorhabdus | 0.92248 | 0.78556 | 0.03760 | 0.27145 | 0.87419 | 0.06339 | 0.17643 | 0.87830 | 0.02458 | 0.22177 | 0.74951 | 0.04181 | 0.09066 | 0.02494 | 0.24967 | 0.73976 | 0.04206 | 0.09203 | 0.72738 | 0.02134 | NA | 0.68995 | 0.23989 | 0.82774 | 0.80591 | 0.02959 | 0.14632 | 0.72236 |
| k__Bacteria;p__Actinobacteriota;c__Coriobacteriia;o__Coriobacteriales;f__Eggerthellaceae;g__Other | NA | NA | NA | 0.38919 | NA | NA | NA | NA | NA | 0.31929 | NA | NA | NA | NA | 0.24471 | NA | NA | NA | 0.67249 | NA | NA | NA | 0.29825 | 0.83537 | 0.65282 | NA | NA | NA |
| k__Bacteria;p__Bacteroidota;c__Bacteroidia;o__Bacteroidales;f__Bacteroidaceae;g__Bacteroides | 0.89853 | 0.01133 | 0.00984 | 0.00962 | 0.04058 | 0.01050 | 0.00943 | 0.01078 | 0.00898 | 0.00867 | 0.06090 | 0.00933 | 0.00858 | 0.19450 | 0.04490 | 0.23256 | 0.01089 | 0.17877 | 0.81522 | 0.01627 | 0.70109 | 0.70799 | 0.01020 | 0.93053 | 0.40088 | 0.01048 | 0.01110 | 0.06729 |
| k__Bacteria;p__Bacteroidota;c__Bacteroidia;o__Bacteroidales;f__Muribaculaceae;g__Muribaculum | 0.76965 | 0.99837 | 0.27859 | 0.01543 | 0.26777 | 0.85897 | 0.41798 | 0.94557 | 0.84418 | 0.01178 | 0.06741 | 0.57453 | 0.66138 | 0.97345 | 0.73189 | 0.83049 | 0.92139 | 0.53105 | 0.05842 | 0.03293 | 0.71339 | 0.89491 | 0.80117 | 0.68968 | 0.28485 | 0.59095 | 0.13964 | 0.58608 |
| k__Bacteria;p__Bacteroidota;c__Bacteroidia;o__Bacteroidales;f__Muribaculaceae;g__Other | 0.94844 | 0.24212 | 0.86941 | 0.05703 | 0.81489 | 0.05915 | 0.02585 | 0.40265 | 0.83186 | 0.13273 | 0.94727 | 0.13222 | 0.07810 | 0.47564 | 0.01217 | 0.41376 | 0.01208 | 0.01481 | 0.68755 | 0.87526 | 0.64229 | 0.69771 | 0.03524 | 0.90864 | 0.81519 | 0.03029 | 0.01195 | 0.84795 |
| k__Bacteria;p__Bacteroidota;c__Bacteroidia;o__Bacteroidales;f__Other;g__Other | 0.90668 | 0.71358 | 0.47652 | 0.55275 | 0.77615 | 0.61061 | 0.64008 | 0.78677 | 0.55895 | 0.53661 | 0.74807 | 0.67749 | 0.63416 | 0.73174 | 0.80422 | 0.83686 | 0.76790 | 0.82710 | 0.83462 | 0.51088 | 0.76370 | 0.76774 | 0.50419 | 0.79695 | 0.74949 | 0.61417 | 0.61914 | 0.78976 |
| k__Bacteria;p__Bacteroidota;c__Bacteroidia;o__Bacteroidales;f__Porphyromonadaceae;g__Other | NA | 0.69607 | NA | NA | NA | NA | NA | 0.71695 | NA | NA | NA | NA | NA | 0.58846 | 0.49217 | 0.79678 | 0.56097 | 0.55632 | NA | NA | NA | NA | NA | NA | NA | NA | NA | NA |
| k__Bacteria;p__Bacteroidota;c__Bacteroidia;o__Bacteroidales;f__Prevotellaceae;g__Prevotellaceae NK3B31 group | 0.92512 | 0.26255 | 0.58738 | 0.63668 | 0.37020 | 0.64920 | 0.55823 | 0.12572 | 0.29009 | 0.34490 | 0.12629 | 0.39899 | 0.22605 | NA | NA | NA | NA | NA | NA | NA | NA | NA | NA | NA | NA | NA | NA | NA |
| k__Bacteria;p__Bacteroidota;c__Bacteroidia;o__Bacteroidales;f__Prevotellaceae;g__Prevotellaceae UCG-001 | 0.91676 | 0.01005 | 0.00979 | 0.00962 | 0.01180 | 0.01050 | 0.00943 | 0.01011 | 0.00898 | 0.00871 | 0.01290 | 0.00933 | 0.00858 | 0.72305 | 0.54323 | 0.85799 | 0.59161 | 0.67192 | 0.66285 | 0.66476 | 0.75237 | NA | 0.51266 | 0.86264 | 0.61867 | 0.56069 | 0.70721 | 0.63037 |
| k__Bacteria;p__Bacteroidota;c__Bacteroidia;o__Bacteroidales;f__Rikenellaceae;g__Alistipes | 0.85047 | 0.02210 | 0.82669 | 0.66831 | 0.75599 | 0.34080 | 0.94295 | 0.01011 | 0.69903 | 0.84958 | 0.77647 | 0.39717 | 0.25360 | 0.01622 | 0.01433 | 0.13279 | 0.01044 | 0.02366 | 0.84289 | 0.88895 | 0.64131 | 0.92765 | 0.85159 | 0.74972 | 0.67999 | 0.44370 | 0.73023 | 0.17812 |
| k__Bacteria;p__Bacteroidota;c__Bacteroidia;o__Bacteroidales;f__Tannerellaceae;g__Other | 0.96403 | 0.01005 | 0.00979 | 0.00962 | 0.01180 | 0.01050 | 0.00943 | 0.01083 | 0.00898 | 0.00867 | 0.01290 | 0.00933 | 0.00863 | 0.78258 | 0.50217 | 0.81429 | 0.52872 | 0.93484 | 0.60020 | 0.14722 | 0.83981 | 0.69095 | 0.91049 | 0.42262 | 0.62627 | 0.02027 | 0.64248 | 0.54852 |
| k__Bacteria;p__Bacteroidota;c__Bacteroidia;o__Other;f__Other;g__Other | NA | NA | NA | NA | 0.61511 | NA | NA | NA | NA | NA | 0.69511 | NA | NA | NA | NA | 0.67561 | NA | NA | NA | 0.74861 | NA | NA | 0.81366 | NA | NA | 0.77587 | 0.74352 | NA |
| k__Bacteria;p__Bacteroidota;c__Other;o__Other;f__Other;g__Other | 0.88433 | 0.49308 | 0.72291 | 0.67155 | 0.57694 | 0.67181 | 0.69807 | NA | NA | NA | NA | NA | NA | NA | NA | NA | NA | NA | NA | NA | NA | NA | NA | NA | NA | NA | NA | NA |
| k__Bacteria;p__Cyanobacteria;c__Cyanobacteriia;o__Chloroplast;f__Other;g__Other | 0.76642 | 0.28828 | 0.66749 | 0.73066 | 0.32610 | 0.76264 | 0.60483 | NA | NA | NA | NA | NA | NA | NA | NA | NA | NA | NA | NA | NA | NA | NA | NA | NA | NA | NA | NA | NA |
| k__Bacteria;p__Cyanobacteria;c__Vampirivibrionia;o__Gastranaerophilales;f__Other;g__Other | 0.83700 | NA | 0.12033 | NA | NA | 0.01050 | NA | 0.50084 | 0.22612 | 0.62155 | 0.58961 | 0.06759 | 0.66551 | 0.09595 | NA | NA | 0.01044 | NA | 0.46479 | 0.09592 | 0.91053 | 0.37631 | NA | 0.04355 | NA | 0.01048 | NA | 0.05063 |
| k__Bacteria;p__Deferribacterota;c__Deferribacteres;o__Deferribacterales;f__Deferribacteraceae;g__Mucispirillum | 0.90917 | 0.19080 | 0.82523 | 0.17072 | 0.05926 | 0.02602 | 0.63495 | 0.12800 | 0.55238 | 0.09975 | 0.03721 | 0.01562 | 0.43142 | 0.93286 | 0.75506 | 0.56397 | 0.10439 | 0.82559 | 0.74671 | 0.57545 | 0.60671 | 0.95290 | 0.86135 | 0.73324 | 0.76031 | 0.56972 | 0.61669 | 0.48272 |
| k__Bacteria;p__Firmicutes;c__Bacilli;o__Acholeplasmatales;f__Acholeplasmataceae;g__Anaeroplasma | 0.94416 | 0.01005 | 0.00979 | 0.00962 | 0.04018 | 0.01050 | 0.00943 | 0.01011 | 0.00898 | 0.00867 | 0.03557 | 0.00933 | 0.00858 | 0.79352 | 0.79931 | 0.89403 | 0.83107 | 0.81418 | NA | 0.68410 | NA | 0.78818 | 0.78140 | NA | 0.75377 | 0.75270 | 0.77827 | 0.78189 |
| k__Bacteria;p__Firmicutes;c__Bacilli;o__Erysipelotrichales;f__Erysipelatoclostridiaceae;g__Candidatus Stoquefichus | 0.89729 | 0.01056 | 0.63557 | 0.51803 | 0.51879 | 0.53382 | 0.65883 | 0.01065 | NA | NA | 0.48101 | NA | NA | 0.01574 | 0.02847 | 0.36006 | 0.02637 | 0.01873 | NA | 0.65681 | NA | NA | 0.75159 | NA | NA | 0.72565 | 0.66200 | NA |
| k__Bacteria;p__Firmicutes;c__Bacilli;o__Erysipelotrichales;f__Erysipelatoclostridiaceae;g__Erysipelatoclostridium | NA | NA | 0.14751 | NA | NA | NA | NA | NA | 0.18020 | NA | NA | NA | NA | 0.12504 | NA | NA | NA | NA | 0.56680 | 0.14397 | 0.64910 | 0.44115 | NA | NA | NA | NA | NA | NA |
| k__Bacteria;p__Firmicutes;c__Bacilli;o__Erysipelotrichales;f__Erysipelotrichaceae;g__[Clostridium] innocuum group | NA | 0.38601 | NA | NA | 0.71444 | NA | NA | 0.45312 | NA | NA | 0.72553 | NA | NA | 0.66752 | 0.72663 | 0.68046 | 0.76624 | 0.66715 | NA | 0.62507 | NA | NA | 0.56058 | NA | NA | 0.57372 | 0.67270 | NA |
| k__Bacteria;p__Firmicutes;c__Bacilli;o__Erysipelotrichales;f__Erysipelotrichaceae;g__Allobaculum | NA | NA | NA | NA | NA | 0.22319 | NA | NA | NA | NA | NA | 0.29714 | NA | NA | NA | NA | 0.19655 | NA | NA | NA | 0.68548 | NA | NA | 0.71924 | NA | 0.24610 | NA | 0.53496 |
| k__Bacteria;p__Firmicutes;c__Bacilli;o__Erysipelotrichales;f__Erysipelotrichaceae;g__Dubosiella | NA | NA | 0.00979 | 0.26222 | 0.66264 | 0.18378 | 0.23094 | NA | 0.00898 | 0.22385 | 0.76265 | 0.16159 | 0.22106 | 0.00976 | 0.15599 | 0.74914 | 0.15864 | 0.13102 | 0.06241 | 0.00942 | 0.84459 | 0.99963 | 0.31054 | 0.81803 | 0.77583 | 0.23466 | 0.26970 | 0.82251 |
| k__Bacteria;p__Firmicutes;c__Bacilli;o__Erysipelotrichales;f__Erysipelotrichaceae;g__Faecalibaculum | 0.90046 | 0.54697 | 0.03580 | 0.00962 | 0.59033 | 0.01050 | 0.00943 | 0.51612 | 0.03386 | 0.00867 | 0.57520 | 0.00933 | 0.00858 | 0.02859 | 0.01075 | NA | 0.01044 | 0.01472 | 0.30745 | 0.02455 | 0.63416 | 0.38786 | 0.01020 | 0.86819 | 0.63811 | 0.01048 | 0.01110 | 0.95945 |
| k__Bacteria;p__Firmicutes;c__Bacilli;o__Erysipelotrichales;f__Erysipelotrichaceae;g__Faecalitalea | NA | NA | 0.44993 | 0.29547 | NA | 0.33043 | NA | NA | 0.47031 | 0.28121 | NA | 0.33367 | NA | 0.35577 | 0.18440 | NA | 0.28911 | NA | 0.74094 | 0.44442 | 0.82801 | 0.76599 | 0.22874 | 0.85770 | 0.61455 | 0.29848 | NA | 0.61058 |
| k__Bacteria;p__Firmicutes;c__Bacilli;o__Erysipelotrichales;f__Erysipelotrichaceae;g__Other | 0.40440 | 0.11442 | 0.28135 | 0.90260 | 0.64351 | 0.42042 | 0.40056 | 0.69903 | 0.63177 | 0.13168 | 0.60828 | 0.49078 | 0.03902 | 0.81446 | 0.19525 | 0.81116 | 0.73811 | 0.06503 | 0.48691 | 0.80185 | 0.79930 | 0.31759 | 0.58313 | 0.66398 | 0.64208 | 0.84648 | 0.20802 | 0.33914 |
| k__Bacteria;p__Firmicutes;c__Bacilli;o__Erysipelotrichales;f__Erysipelotrichaceae;g__Turicibacter | NA | 0.26864 | 0.27229 | NA | 0.42763 | NA | NA | 0.37325 | 0.30240 | NA | 0.48675 | NA | NA | 0.72084 | 0.73066 | 0.89059 | 0.72065 | 0.58944 | 0.71469 | 0.81952 | 0.74382 | 0.61045 | 0.74305 | NA | NA | 0.74463 | 0.66703 | NA |
| k__Bacteria;p__Firmicutes;c__Bacilli;o__Lactobacillales;f__Enterococcaceae;g__Enterococcus | NA | 0.68859 | 0.49813 | 0.09032 | NA | NA | NA | 0.70175 | 0.50846 | 0.07342 | NA | NA | NA | 0.56891 | 0.10723 | 0.81969 | 0.53554 | 0.58434 | 0.53921 | 0.37085 | 0.82152 | 0.73881 | 0.10354 | 0.45508 | 0.31822 | NA | NA | NA |
| k__Bacteria;p__Firmicutes;c__Bacilli;o__Lactobacillales;f__Lactobacillaceae;g__Lactobacillus | 0.27825 | 0.03449 | 0.10386 | 0.77303 | 0.30412 | 0.65245 | 0.52612 | 0.46925 | 0.41788 | 0.05446 | 0.24412 | 0.19519 | 0.02505 | 0.94219 | 0.10411 | 0.72950 | 0.69753 | 0.03076 | 0.40878 | 0.58403 | 0.79673 | 0.25784 | 0.57469 | 0.91111 | 0.62673 | 0.82454 | 0.15818 | 0.44073 |
| k__Bacteria;p__Firmicutes;c__Bacilli;o__Lactobacillales;f__Other;g__Other | 0.89118 | 0.51962 | 0.69392 | 0.64810 | 0.73831 | 0.65588 | 0.66618 | NA | NA | NA | 0.69908 | NA | NA | NA | NA | 0.77872 | NA | NA | NA | 0.64474 | NA | NA | 0.47880 | NA | NA | 0.58019 | 0.64011 | NA |
| k__Bacteria;p__Firmicutes;c__Bacilli;o__Lactobacillales;f__Streptococcaceae;g__Lactococcus | NA | 0.01005 | 0.00979 | 0.03601 | 0.01180 | 0.01050 | 0.00943 | 0.01011 | 0.00898 | 0.02983 | 0.01290 | 0.00933 | 0.00858 | 0.10535 | 0.30278 | 0.93050 | 0.10584 | 0.77186 | 0.86915 | 0.11880 | 0.89311 | 0.59592 | 0.30376 | 0.82101 | 0.81591 | 0.09469 | 0.71329 | 0.50936 |
| k__Bacteria;p__Firmicutes;c__Bacilli;o__RF39;f__Other;g__Other | 0.94403 | 0.90542 | 0.68819 | 0.43441 | 0.47808 | 0.94993 | 0.96289 | 0.85857 | 0.63115 | 0.19418 | 0.64609 | 0.91358 | 0.90588 | 0.73782 | 0.23976 | 0.68544 | 0.95953 | 0.97256 | 0.71891 | 0.23579 | 0.88848 | 0.87308 | 0.16907 | 0.81388 | 0.47002 | 0.62406 | 0.54871 | 0.97294 |
| k__Bacteria;p__Firmicutes;c__Bacilli;o__Staphylococcales;f__Staphylococcaceae;g__Staphylococcus | NA | 0.68335 | NA | NA | NA | NA | NA | 0.73436 | NA | NA | NA | NA | NA | 0.83318 | 0.78207 | 0.80064 | 0.82857 | 0.82413 | NA | NA | NA | NA | NA | NA | NA | NA | NA | NA |
| k__Bacteria;p__Firmicutes;c__Clostridia;o__Christensenellales;f__Christensenellaceae;g__Other | NA | 0.71731 | NA | NA | NA | NA | NA | 0.73620 | NA | NA | NA | NA | NA | 0.67442 | 0.54000 | 0.77933 | 0.62737 | 0.67045 | NA | NA | NA | NA | NA | NA | NA | NA | NA | NA |
| k__Bacteria;p__Firmicutes;c__Clostridia;o__Clostridia UCG-014;f__Other;g__Other | 0.93240 | 0.44142 | 0.87947 | 0.02820 | 0.02637 | 0.51958 | 0.64103 | 0.34405 | 0.83153 | 0.03025 | 0.02997 | 0.58128 | 0.82955 | 0.77598 | 0.15110 | 0.29195 | 0.45772 | 0.45242 | 0.46338 | 0.17825 | 0.67047 | 0.86645 | 0.72812 | 0.34486 | 0.35457 | 0.10049 | 0.11558 | 0.65363 |
| k__Bacteria;p__Firmicutes;c__Clostridia;o__Clostridia vadinBB60 group;f__Other;g__Other | 0.93466 | 0.01862 | 0.00979 | 0.04165 | 0.02349 | 0.01574 | 0.00943 | 0.02947 | 0.00898 | 0.04010 | 0.03923 | 0.03298 | 0.00858 | 0.13518 | 0.46094 | 0.80060 | 0.47580 | 0.10396 | 0.94507 | 0.06889 | 0.89026 | 0.87657 | 0.31439 | 0.93164 | 0.91414 | 0.37730 | 0.04394 | 0.90735 |
| k__Bacteria;p__Firmicutes;c__Clostridia;o__Clostridiales;f__Clostridiaceae;g__Candidatus Arthromitus | 0.88724 | 0.17864 | 0.24075 | 0.29560 | 0.23774 | 0.30554 | 0.24006 | 0.15976 | 0.18393 | 0.23275 | 0.16768 | 0.28499 | 0.17894 | 0.76769 | 0.72152 | 0.88920 | 0.70336 | 0.75916 | NA | 0.69072 | NA | NA | 0.64117 | NA | NA | 0.65837 | 0.68658 | NA |
| k__Bacteria;p__Firmicutes;c__Clostridia;o__Clostridiales;f__Clostridiaceae;g__Clostridium sensu stricto 1 | 0.87071 | 0.04434 | 0.42160 | 0.66992 | 0.01806 | 0.66618 | 0.69522 | 0.01011 | 0.19842 | NA | 0.01290 | NA | NA | 0.24311 | 0.01335 | 0.26870 | 0.01151 | 0.01530 | 0.56218 | 0.03053 | 0.62810 | 0.48207 | 0.01046 | NA | NA | 0.01058 | 0.01110 | NA |
| k__Bacteria;p__Firmicutes;c__Clostridia;o__Clostridiales;f__Other;g__Other | NA | NA | NA | NA | 0.67027 | NA | NA | NA | NA | NA | 0.70873 | NA | NA | NA | NA | 0.79504 | NA | NA | NA | 0.68980 | NA | NA | 0.62084 | NA | NA | 0.62288 | 0.63315 | NA |
| k__Bacteria;p__Firmicutes;c__Clostridia;o__Hungateiclostridiaceae;f__Ruminiclostridium;g__Other | 0.83772 | NA | NA | NA | NA | NA | NA | 0.50135 | 0.69948 | 0.59943 | 0.57176 | 0.62553 | 0.65171 | NA | NA | NA | NA | NA | NA | NA | NA | NA | NA | NA | NA | NA | NA | NA |
| k__Bacteria;p__Firmicutes;c__Clostridia;o__Lachnospirales;f__Lachnospiraceae;g__[Eubacterium] fissicatena group | NA | NA | NA | NA | 0.42739 | NA | NA | NA | NA | NA | 0.46226 | NA | NA | NA | NA | 0.53274 | NA | NA | NA | 0.61907 | NA | NA | 0.74130 | NA | NA | 0.75818 | 0.62378 | NA |
| k__Bacteria;p__Firmicutes;c__Clostridia;o__Lachnospirales;f__Lachnospiraceae;g__[Eubacterium] oxidoreducens group | NA | NA | NA | NA | NA | NA | NA | NA | NA | NA | NA | NA | NA | NA | NA | NA | NA | NA | NA | NA | NA | NA | NA | NA | NA | NA | NA | NA |
| k__Bacteria;p__Firmicutes;c__Clostridia;o__Lachnospirales;f__Lachnospiraceae;g__[Eubacterium] xylanophilum group | 0.93114 | 0.01005 | 0.00979 | 0.00962 | 0.01180 | 0.01050 | 0.00943 | 0.01011 | 0.00898 | 0.00867 | 0.01290 | 0.00937 | 0.00858 | NA | NA | NA | NA | NA | NA | NA | NA | NA | NA | NA | NA | NA | NA | NA |
| k__Bacteria;p__Firmicutes;c__Clostridia;o__Lachnospirales;f__Lachnospiraceae;g__[Ruminococcus] torques group | NA | NA | NA | NA | NA | NA | 0.40447 | NA | NA | NA | NA | NA | 0.44521 | NA | NA | NA | NA | 0.28961 | NA | NA | NA | 0.77188 | NA | NA | 0.77164 | NA | 0.34089 | 0.80129 |
| k__Bacteria;p__Firmicutes;c__Clostridia;o__Lachnospirales;f__Lachnospiraceae;g__A2 | 0.89940 | 0.82277 | 0.00979 | 0.01223 | 0.88265 | 0.01050 | 0.04920 | 0.83328 | 0.00904 | 0.01116 | 0.81867 | 0.00933 | 0.05541 | 0.00976 | 0.01080 | 0.94086 | 0.01044 | 0.04201 | 0.57004 | 0.00942 | NA | 0.39771 | 0.01020 | 0.73036 | 0.74612 | 0.01048 | 0.03790 | 0.39650 |
| k__Bacteria;p__Firmicutes;c__Clostridia;o__Lachnospirales;f__Lachnospiraceae;g__Acetatifactor | 0.68682 | 0.01005 | 0.31966 | 0.17358 | 0.01251 | 0.10275 | 0.17300 | 0.01400 | 0.09810 | 0.02943 | 0.04130 | 0.01676 | 0.02177 | 0.00976 | 0.01075 | 0.69975 | 0.01044 | 0.01472 | 0.79535 | 0.01220 | 0.81833 | 0.83632 | 0.01026 | 0.79708 | 0.77846 | 0.01048 | 0.01110 | 0.76766 |
| k__Bacteria;p__Firmicutes;c__Clostridia;o__Lachnospirales;f__Lachnospiraceae;g__Anaerostipes | NA | 0.44940 | 0.11823 | 0.00962 | 0.07632 | 0.37663 | 0.00943 | 0.55885 | 0.10342 | 0.00867 | 0.07949 | 0.37793 | 0.00858 | 0.22424 | 0.01075 | 0.67161 | 0.79921 | 0.01472 | 0.07069 | 0.40299 | 0.60193 | 0.50293 | 0.01020 | 0.04338 | 0.21583 | 0.62217 | 0.01110 | 0.06541 |
| k__Bacteria;p__Firmicutes;c__Clostridia;o__Lachnospirales;f__Lachnospiraceae;g__ASF356 | 0.94531 | 0.08880 | 0.09960 | 0.69219 | 0.17048 | 0.43595 | 0.01172 | 0.09852 | 0.08226 | 0.50803 | 0.13909 | 0.32099 | 0.00902 | 0.34204 | 0.98063 | 0.95646 | 0.77415 | 0.11719 | 0.83128 | 0.43302 | 0.79882 | 0.86812 | 0.95939 | 0.93805 | 0.93625 | 0.85523 | 0.24524 | 0.89865 |
| k__Bacteria;p__Firmicutes;c__Clostridia;o__Lachnospirales;f__Lachnospiraceae;g__Blautia | NA | NA | 0.38073 | 0.03730 | NA | 0.30214 | NA | NA | 0.40128 | 0.03683 | NA | 0.25572 | NA | 0.27890 | 0.02997 | NA | 0.19559 | NA | 0.53597 | 0.25994 | 0.86490 | 0.66308 | 0.02888 | 0.35097 | 0.21486 | 0.23122 | NA | 0.60933 |
| k__Bacteria;p__Firmicutes;c__Clostridia;o__Lachnospirales;f__Lachnospiraceae;g__Frisingicoccus | NA | NA | 0.45100 | NA | NA | NA | NA | NA | 0.43022 | NA | NA | NA | NA | 0.32807 | NA | NA | NA | NA | 0.79998 | 0.35782 | 0.81374 | 0.70902 | NA | NA | NA | NA | NA | NA |
| k__Bacteria;p__Firmicutes;c__Clostridia;o__Lachnospirales;f__Lachnospiraceae;g__GCA-900066575 | 0.93333 | 0.07048 | 0.56827 | 0.21409 | 0.03677 | 0.80830 | 0.87480 | 0.18313 | 0.67373 | 0.19197 | 0.10492 | 0.86929 | 0.78530 | 0.15215 | 0.03935 | 0.92508 | 0.39099 | 0.56920 | 0.60249 | 0.13419 | 0.91996 | 0.80392 | 0.02652 | 0.52311 | 0.38285 | 0.37419 | 0.49018 | 0.89651 |
| k__Bacteria;p__Firmicutes;c__Clostridia;o__Lachnospirales;f__Lachnospiraceae;g__Lachnoclostridium | 0.74185 | 0.01436 | 0.85283 | 0.10873 | 0.01180 | 0.16768 | 0.00943 | 0.65310 | 0.71302 | 0.52473 | 0.40890 | 0.64338 | 0.07759 | 0.33581 | 0.91858 | 0.84992 | 0.91042 | 0.16580 | 0.65130 | 0.18140 | 0.70211 | 0.35314 | 0.80724 | 0.94580 | 0.81337 | 0.69177 | 0.15856 | 0.58466 |
| k__Bacteria;p__Firmicutes;c__Clostridia;o__Lachnospirales;f__Lachnospiraceae;g__Lachnospiraceae FCS020 group | 0.84412 | 0.41673 | 0.08965 | 0.52335 | 0.53504 | 0.70905 | 0.31273 | 0.93985 | 0.52501 | 0.75626 | 0.96699 | 0.81490 | 0.76012 | 0.15767 | 0.80503 | 0.80185 | 0.84123 | 0.74975 | 0.94504 | 0.02830 | 0.41030 | 0.84301 | 0.67963 | 0.84243 | 0.94338 | 0.77767 | 0.43628 | 0.59789 |
| k__Bacteria;p__Firmicutes;c__Clostridia;o__Lachnospirales;f__Lachnospiraceae;g__Lachnospiraceae NK4A136 group | 0.92854 | 0.06477 | 0.03051 | 0.00962 | 0.01180 | 0.58873 | 0.05888 | 0.11359 | 0.04110 | 0.00943 | 0.01648 | 0.57267 | 0.03981 | 0.48857 | 0.17936 | 0.92904 | 0.85660 | 0.89918 | 0.76983 | 0.52894 | 0.95668 | 0.74013 | 0.16002 | 0.78445 | 0.52217 | 0.83056 | 0.67805 | 0.96941 |
| k__Bacteria;p__Firmicutes;c__Clostridia;o__Lachnospirales;f__Lachnospiraceae;g__Lachnospiraceae NK4B4 group | 0.89605 | 0.39361 | 0.70140 | 0.70930 | 0.50090 | 0.72650 | 0.73481 | 0.49116 | 0.66855 | 0.60588 | 0.59272 | 0.60492 | 0.67829 | NA | NA | NA | NA | NA | NA | NA | NA | NA | NA | NA | NA | NA | NA | NA |
| k__Bacteria;p__Firmicutes;c__Clostridia;o__Lachnospirales;f__Lachnospiraceae;g__Lachnospiraceae UCG-001 | 0.96644 | 0.13324 | 0.91364 | 0.35422 | 0.17229 | 0.70984 | 0.30911 | 0.11102 | 0.91510 | 0.26407 | 0.15168 | 0.63195 | 0.21366 | 0.12975 | NA | NA | 0.25122 | NA | 0.55318 | 0.15231 | 0.77106 | 0.45346 | NA | 0.82002 | NA | 0.31740 | NA | 0.67107 |
| k__Bacteria;p__Firmicutes;c__Clostridia;o__Lachnospirales;f__Lachnospiraceae;g__Lachnospiraceae UCG-004 | 0.47330 | 0.06815 | 0.20651 | 0.14071 | 0.38572 | 0.35972 | 0.43814 | 0.59360 | 0.69587 | 0.61465 | 0.93665 | 0.65605 | 0.52280 | 0.86129 | 0.88728 | 0.71691 | 0.53181 | 0.39204 | 0.92630 | 0.52301 | 0.80560 | 0.58181 | 0.52775 | 0.75655 | 0.53934 | 0.87660 | 0.82084 | 0.78487 |
| k__Bacteria;p__Firmicutes;c__Clostridia;o__Lachnospirales;f__Lachnospiraceae;g__Lachnospiraceae UCG-006 | 0.88357 | 0.01005 | 0.56849 | 0.82982 | 0.01426 | 0.64783 | 0.83438 | 0.01053 | 0.26568 | 0.68143 | 0.04565 | 0.41340 | 0.85073 | 0.00976 | 0.03673 | 0.75760 | 0.01044 | 0.12758 | 0.61821 | 0.00971 | NA | 0.71063 | 0.05943 | 0.79139 | 0.84076 | 0.01072 | 0.19314 | 0.76158 |
| k__Bacteria;p__Firmicutes;c__Clostridia;o__Lachnospirales;f__Lachnospiraceae;g__Lachnospiraceae UCG-010 | 0.88751 | 0.76171 | 0.66397 | 0.70025 | 0.69039 | 0.77889 | 0.60395 | 0.77102 | 0.35998 | 0.48763 | 0.90749 | 0.72785 | 0.32429 | 0.29687 | 0.45151 | 0.83397 | 0.73578 | 0.31714 | NA | 0.39072 | 0.73483 | NA | 0.55087 | 0.80986 | NA | 0.72661 | 0.34940 | 0.67449 |
| k__Bacteria;p__Firmicutes;c__Clostridia;o__Lachnospirales;f__Lachnospiraceae;g__Marvinbryantia | 0.85822 | 0.10961 | 0.04225 | 0.02887 | 0.45377 | 0.65268 | 0.39161 | 0.08627 | 0.48367 | 0.40790 | 0.17306 | 0.87015 | 0.16229 | 0.00999 | 0.01092 | 0.43017 | 0.60801 | 0.93055 | 0.78714 | 0.01145 | 0.74384 | 0.23594 | 0.01076 | 0.74671 | 0.23404 | 0.60698 | 0.57772 | 0.82148 |
| k__Bacteria;p__Firmicutes;c__Clostridia;o__Lachnospirales;f__Lachnospiraceae;g__Other | 0.89015 | 0.65379 | 0.73742 | 0.26857 | 0.69889 | 0.23196 | 0.32191 | 0.91507 | 0.87305 | 0.38397 | 0.87632 | 0.35036 | 0.54833 | 0.90676 | 0.49006 | 0.91715 | 0.56503 | 0.71221 | 0.68500 | 0.83588 | 0.67671 | 0.88805 | 0.33715 | 0.94893 | 0.62211 | 0.38224 | 0.43643 | 0.65669 |
| k__Bacteria;p__Firmicutes;c__Clostridia;o__Lachnospirales;f__Lachnospiraceae;g__Roseburia | 0.90503 | 0.09790 | 0.06443 | 0.07688 | 0.11315 | 0.13994 | 0.28619 | 0.27950 | 0.01049 | 0.01085 | 0.25984 | 0.01947 | 0.08640 | 0.00976 | 0.01075 | 0.89765 | 0.01044 | 0.06606 | NA | 0.00942 | 0.69307 | 0.76922 | 0.01020 | 0.82075 | 0.82750 | 0.01048 | 0.05364 | 0.77104 |
| k__Bacteria;p__Firmicutes;c__Clostridia;o__Lachnospirales;f__Lachnospiraceae;g__Sellimonas | NA | NA | 0.41493 | NA | NA | NA | NA | NA | 0.45723 | NA | NA | NA | NA | 0.33689 | NA | NA | NA | NA | 0.78855 | 0.36598 | 0.80247 | 0.75441 | NA | NA | NA | NA | NA | NA |
| k__Bacteria;p__Firmicutes;c__Clostridia;o__Lachnospirales;f__Lachnospiraceae;g__Tuzzerella | NA | NA | NA | NA | 0.42436 | NA | NA | NA | NA | NA | 0.44399 | NA | NA | NA | NA | 0.55564 | NA | NA | NA | 0.57303 | NA | NA | 0.70104 | NA | NA | 0.67511 | 0.63514 | NA |
| k__Bacteria;p__Firmicutes;c__Clostridia;o__Lachnospirales;f__Lachnospiraceae;g__Tyzzerella | 0.89646 | 0.01005 | 0.00986 | 0.01009 | 0.01180 | 0.01090 | 0.00943 | 0.01011 | 0.00915 | 0.00925 | 0.01290 | 0.00942 | 0.00858 | NA | NA | NA | NA | NA | NA | NA | NA | NA | NA | NA | NA | NA | NA | NA |
| k__Bacteria;p__Firmicutes;c__Clostridia;o__Lachnospirales;f__Other;g__Other | NA | NA | NA | NA | NA | NA | 0.43041 | NA | NA | NA | NA | NA | 0.38602 | NA | NA | NA | NA | 0.30547 | NA | NA | NA | 0.75367 | NA | NA | 0.77314 | NA | 0.35960 | 0.73760 |
| k__Bacteria;p__Firmicutes;c__Clostridia;o__Monoglobales;f__Monoglobaceae;g__Monoglobus | 0.90622 | 0.87932 | 0.60203 | 0.70732 | 0.79756 | 0.68759 | 0.83026 | 0.76595 | 0.51324 | 0.59299 | 0.90083 | 0.65235 | 0.84433 | 0.70828 | 0.75807 | 0.89652 | 0.78030 | 0.80939 | NA | 0.40958 | NA | 0.71964 | 0.54968 | NA | 0.77853 | 0.53688 | 0.86392 | 0.79558 |
| k__Bacteria;p__Firmicutes;c__Clostridia;o__Oscillospirales;f__[Eubacterium] coprostanoligenes group;g__Other | 0.93172 | 0.01005 | 0.31793 | 0.79854 | 0.01180 | 0.89446 | 0.08283 | 0.01011 | 0.08652 | 0.50654 | 0.01290 | 0.90988 | 0.15326 | 0.00976 | 0.01075 | 0.91230 | 0.01478 | 0.04859 | 0.71848 | 0.00942 | 0.65999 | 0.08304 | 0.01020 | 0.84682 | 0.21625 | 0.01114 | 0.03344 | 0.52017 |
| k__Bacteria;p__Firmicutes;c__Clostridia;o__Oscillospirales;f__Butyricicoccaceae;g__Butyricicoccus | 0.93556 | 0.02055 | 0.62147 | 0.01384 | 0.11614 | 0.27842 | 0.02176 | 0.12517 | 0.89938 | 0.07268 | 0.19079 | 0.57162 | 0.08291 | 0.36968 | 0.36331 | 0.48609 | 0.85860 | 0.52959 | 0.35609 | 0.64440 | 0.76336 | 0.37915 | 0.11866 | 0.75999 | 0.76784 | 0.78027 | 0.25054 | 0.66828 |
| k__Bacteria;p__Firmicutes;c__Clostridia;o__Oscillospirales;f__Oscillospiraceae;g__Colidextribacter | 0.91949 | 0.10863 | 0.17116 | 0.54873 | 0.01909 | 0.84304 | 0.26350 | 0.03561 | 0.45078 | 0.77570 | 0.01389 | 0.46553 | 0.43951 | 0.01580 | 0.10126 | 0.76251 | 0.41628 | 0.05768 | 0.92417 | 0.00967 | 0.55001 | 0.92552 | 0.01966 | 0.72131 | 0.82029 | 0.19012 | 0.01397 | 0.32232 |
| k__Bacteria;p__Firmicutes;c__Clostridia;o__Oscillospirales;f__Oscillospiraceae;g__Flavonifractor | NA | NA | 0.24801 | 0.10636 | NA | 0.26291 | 0.31715 | NA | 0.25858 | 0.10752 | NA | 0.29864 | 0.32398 | 0.17348 | 0.07907 | NA | 0.17173 | 0.22970 | 0.81707 | 0.22237 | 0.89500 | 0.86481 | 0.10337 | 0.85678 | 0.65118 | 0.24226 | 0.32485 | 0.80959 |
| k__Bacteria;p__Firmicutes;c__Clostridia;o__Oscillospirales;f__Oscillospiraceae;g__Intestinimonas | 0.90988 | 0.33374 | 0.00979 | 0.04933 | 0.08693 | 0.27129 | 0.01269 | 0.65001 | 0.00916 | 0.07946 | 0.05499 | 0.47093 | 0.02081 | 0.01146 | 0.12731 | 0.17179 | 0.58601 | 0.04446 | 0.65622 | 0.00942 | 0.64184 | 0.92911 | 0.01301 | 0.87439 | 0.67549 | 0.02585 | 0.01110 | 0.56790 |
| k__Bacteria;p__Firmicutes;c__Clostridia;o__Oscillospirales;f__Oscillospiraceae;g__Oscillibacter | 0.81220 | 0.88982 | 0.82676 | 0.85904 | 0.18905 | 0.12886 | 0.20588 | 0.60177 | 0.82431 | 0.89885 | 0.85022 | 0.44408 | 0.05524 | 0.93425 | 0.95398 | 0.70833 | 0.37531 | 0.17854 | 0.92595 | 0.63792 | 0.62281 | 0.45712 | 0.77601 | 0.80421 | 0.67012 | 0.74432 | 0.02543 | 0.05157 |
| k__Bacteria;p__Firmicutes;c__Clostridia;o__Oscillospirales;f__Oscillospiraceae;g__Other | 0.73098 | 0.16513 | 0.02273 | 0.17423 | 0.02079 | 0.86991 | 0.12433 | 0.02700 | 0.07267 | 0.42973 | 0.01354 | 0.81675 | 0.35413 | 0.00998 | 0.08539 | 0.47641 | 0.42271 | 0.03358 | 0.96532 | 0.00942 | 0.70421 | 0.94273 | 0.02653 | 0.83042 | 0.93522 | 0.05978 | 0.01188 | 0.76341 |
| k__Bacteria;p__Firmicutes;c__Clostridia;o__Oscillospirales;f__Oscillospiraceae;g__Pseudoflavonifractor | 0.90100 | 0.53165 | 0.69726 | 0.57152 | 0.62615 | 0.61789 | 0.68878 | NA | NA | NA | NA | NA | NA | NA | NA | NA | NA | NA | NA | NA | NA | NA | NA | NA | NA | NA | NA | NA |
| k__Bacteria;p__Firmicutes;c__Clostridia;o__Oscillospirales;f__Oscillospiraceae;g__UCG-005 | 0.18766 | 0.01559 | 0.09044 | 0.38153 | 0.02565 | 0.06460 | 0.84257 | 0.80912 | 0.39374 | 0.83016 | 0.78593 | 0.75049 | 0.12111 | 0.86565 | 0.72542 | 0.82908 | 0.82743 | 0.08068 | 0.71422 | 0.38612 | 0.82130 | 0.33025 | 0.86709 | 0.87567 | 0.57064 | 0.72405 | 0.17762 | 0.36349 |
| k__Bacteria;p__Firmicutes;c__Clostridia;o__Oscillospirales;f__Other;g__Other | NA | NA | NA | NA | 0.58743 | NA | NA | NA | NA | NA | 0.69270 | NA | NA | NA | NA | 0.70574 | NA | NA | NA | 0.72240 | NA | NA | 0.74029 | NA | NA | 0.73005 | 0.72892 | NA |
| k__Bacteria;p__Firmicutes;c__Clostridia;o__Oscillospirales;f__Ruminococcaceae;g__[Eubacterium] siraeum group | 0.91607 | 0.01005 | 0.00997 | 0.01013 | 0.01180 | 0.01063 | 0.00946 | 0.01011 | 0.00898 | 0.00867 | 0.01290 | 0.00933 | 0.00858 | NA | NA | NA | NA | NA | NA | NA | NA | NA | NA | NA | NA | NA | NA | NA |
| k__Bacteria;p__Firmicutes;c__Clostridia;o__Oscillospirales;f__Ruminococcaceae;g__Anaerotruncus | 0.77370 | 0.24056 | 0.18184 | 0.59958 | 0.56197 | 0.88365 | 0.18989 | 0.53159 | 0.03451 | 0.86983 | 0.90298 | 0.67266 | 0.02978 | 0.02662 | 0.74545 | 0.76689 | 0.34382 | 0.03206 | 0.48384 | 0.03012 | 0.61602 | 0.80259 | 0.93022 | 0.87451 | 0.37724 | 0.63442 | 0.03813 | 0.38856 |
| k__Bacteria;p__Firmicutes;c__Clostridia;o__Oscillospirales;f__Ruminococcaceae;g__Candidatus Soleaferrea | NA | 0.63144 | NA | NA | NA | NA | NA | 0.66905 | NA | NA | NA | NA | NA | 0.79923 | 0.80370 | 0.74698 | 0.82869 | 0.79043 | NA | NA | NA | NA | NA | NA | NA | NA | NA | NA |
| k__Bacteria;p__Firmicutes;c__Clostridia;o__Oscillospirales;f__Ruminococcaceae;g__Harryflintia | NA | 0.38395 | NA | NA | NA | NA | NA | 0.46766 | NA | NA | NA | NA | NA | 0.62770 | 0.71539 | 0.58347 | 0.74222 | 0.60199 | NA | NA | NA | NA | NA | NA | NA | NA | NA | NA |
| k__Bacteria;p__Firmicutes;c__Clostridia;o__Oscillospirales;f__Ruminococcaceae;g__Incertae Sedis | 0.90485 | 0.62639 | 0.00979 | 0.01100 | 0.64484 | 0.01590 | 0.02165 | 0.14234 | 0.00898 | 0.00867 | 0.14702 | 0.00933 | 0.00858 | 0.01132 | 0.01636 | 0.94681 | 0.01785 | 0.04333 | 0.83910 | 0.00957 | 0.84694 | 0.71164 | 0.01316 | 0.83557 | 0.75534 | 0.01870 | 0.03681 | 0.76383 |
| k__Bacteria;p__Firmicutes;c__Clostridia;o__Oscillospirales;f__Ruminococcaceae;g__Other | 0.28771 | 0.06022 | 0.88785 | 0.89129 | 0.01180 | 0.73236 | 0.51408 | 0.69814 | 0.34388 | 0.49004 | 0.01843 | 0.62485 | 0.03456 | 0.14338 | 0.28887 | 0.70679 | 0.33595 | 0.04687 | 0.93468 | 0.01147 | 0.85838 | 0.79756 | 0.07541 | 0.92698 | 0.74058 | 0.01658 | 0.01110 | 0.45620 |
| k__Bacteria;p__Firmicutes;c__Clostridia;o__Oscillospirales;f__Ruminococcaceae;g__Paludicola | NA | 0.49537 | NA | NA | NA | NA | NA | 0.55246 | NA | NA | NA | NA | NA | 0.72920 | 0.70859 | 0.62826 | 0.78456 | 0.74989 | NA | NA | NA | NA | NA | NA | NA | NA | NA | NA |
| k__Bacteria;p__Firmicutes;c__Clostridia;o__Oscillospirales;f__Ruminococcaceae;g__Ruminococcus | NA | 0.64594 | NA | NA | NA | NA | NA | 0.68746 | NA | NA | NA | NA | NA | 0.56624 | 0.50709 | 0.80720 | 0.54950 | 0.59530 | NA | NA | NA | NA | NA | NA | NA | NA | NA | NA |
| k__Bacteria;p__Firmicutes;c__Clostridia;o__Oscillospirales;f__UCG-010;g__Other | 0.74086 | 0.27041 | 0.56543 | 0.64379 | 0.31323 | 0.62306 | 0.54950 | NA | NA | NA | NA | NA | NA | NA | NA | NA | NA | NA | NA | NA | NA | NA | NA | NA | NA | NA | NA | NA |
| k__Bacteria;p__Firmicutes;c__Clostridia;o__Other;f__Other;g__Other | 0.84012 | 0.41534 | 0.72649 | 0.78590 | 0.60640 | 0.77681 | 0.78865 | NA | NA | NA | 0.66226 | 0.35095 | 0.45239 | NA | NA | 0.75773 | 0.29005 | 0.31271 | NA | 0.68728 | 0.76366 | 0.78739 | 0.60442 | 0.81910 | 0.80308 | 0.50606 | 0.53127 | 0.80750 |
| k__Bacteria;p__Firmicutes;c__Clostridia;o__Peptococcales;f__Peptococcaceae;g__Other | 0.87705 | 0.01033 | 0.19905 | 0.88256 | 0.44151 | 0.71818 | 0.63554 | 0.01584 | 0.36828 | 0.89443 | 0.63229 | 0.71812 | 0.55362 | 0.45903 | 0.32725 | 0.38309 | 1.00000 | 0.14044 | 0.71574 | 0.76290 | 0.95633 | 0.48547 | 0.91717 | 0.87855 | 0.71985 | 0.71550 | 0.65733 | 0.66818 |
| k__Bacteria;p__Firmicutes;c__Clostridia;o__Peptococcales;f__Peptococcaceae;g__Peptococcus | 0.93840 | 0.01024 | 0.41446 | 0.55099 | 0.01192 | 0.55713 | 0.37583 | 0.01329 | 0.55326 | 0.63481 | 0.01380 | 0.69314 | 0.51992 | 0.01084 | 0.01295 | 0.85112 | 0.01138 | 0.01480 | NA | 0.00960 | NA | NA | 0.01059 | NA | NA | 0.01058 | 0.01123 | NA |
| k__Bacteria;p__Firmicutes;c__Clostridia;o__Peptostreptococcales-Tissierellales;f__Anaerovoracaceae;g__[Eubacterium] brachy group | NA | NA | NA | NA | 0.61361 | NA | NA | NA | NA | NA | 0.67829 | NA | NA | NA | NA | 0.69250 | NA | NA | NA | 0.76586 | NA | NA | 0.72009 | NA | NA | 0.72603 | 0.73842 | NA |
| k__Bacteria;p__Firmicutes;c__Clostridia;o__Peptostreptococcales-Tissierellales;f__Anaerovoracaceae;g__[Eubacterium] nodatum group | NA | NA | NA | NA | 0.46573 | NA | NA | NA | NA | NA | 0.51788 | NA | NA | NA | NA | 0.53121 | NA | NA | NA | 0.69235 | NA | NA | 0.81742 | NA | NA | 0.78651 | 0.72282 | NA |
| k__Bacteria;p__Firmicutes;c__Clostridia;o__Peptostreptococcales-Tissierellales;f__Anaerovoracaceae;g__Family XIII AD3011 group | 0.78304 | NA | NA | NA | NA | NA | NA | 0.33596 | 0.72719 | 0.71842 | 0.41051 | 0.75248 | 0.64440 | NA | NA | NA | NA | NA | NA | NA | NA | NA | NA | NA | NA | NA | NA | NA |
| k__Bacteria;p__Firmicutes;c__Clostridia;o__Peptostreptococcales-Tissierellales;f__Other;g__Other | NA | 0.70747 | NA | NA | NA | NA | NA | 0.71025 | NA | NA | NA | NA | NA | 0.60730 | 0.50415 | 0.81143 | 0.57264 | 0.62282 | NA | NA | NA | NA | NA | NA | NA | NA | NA | NA |
| k__Bacteria;p__Firmicutes;c__Clostridia;o__Peptostreptococcales-Tissierellales;f__Peptostreptococcaceae;g__Asaccharospora | NA | NA | NA | NA | NA | NA | NA | NA | NA | NA | NA | NA | NA | NA | NA | NA | NA | NA | NA | NA | NA | NA | NA | NA | NA | NA | NA | NA |
| k__Bacteria;p__Firmicutes;c__Clostridia;o__Peptostreptococcales-Tissierellales;f__Peptostreptococcaceae;g__Clostridioides | NA | NA | 0.45205 | NA | NA | NA | NA | NA | 0.51517 | NA | NA | NA | NA | 0.36549 | NA | NA | NA | NA | 0.83098 | 0.42499 | 0.82355 | 0.77504 | NA | NA | NA | NA | NA | NA |
| k__Bacteria;p__Firmicutes;c__Clostridia;o__Peptostreptococcales-Tissierellales;f__Peptostreptococcaceae;g__Romboutsia | NA | 0.39148 | NA | NA | 0.06091 | NA | NA | 0.47488 | NA | NA | 0.06788 | NA | NA | 0.61634 | 0.70078 | 0.91178 | 0.75081 | 0.58056 | NA | 0.09366 | NA | NA | 0.10042 | NA | NA | 0.10451 | 0.08439 | NA |
| k__Bacteria;p__Firmicutes;c__Other;o__Other;f__Other;g__Other | NA | 0.46418 | 0.40820 | NA | 0.66919 | NA | NA | 0.54002 | 0.46049 | NA | 0.73108 | NA | NA | 0.85447 | 0.81418 | 0.74224 | 0.79114 | 0.72040 | 0.78307 | 0.51068 | 0.83389 | 0.76249 | 0.62776 | NA | NA | 0.60379 | 0.66180 | NA |
| k__Bacteria;p__Other;c__Other;o__Other;f__Other;g__Other | 0.91595 | 0.84805 | 0.75534 | 0.78760 | 0.82188 | 0.78609 | 0.74952 | 0.72606 | 0.70018 | 0.58835 | 0.71057 | 0.56485 | 0.67465 | 0.72874 | 0.76795 | 0.93953 | 0.86967 | 0.73628 | NA | 0.65586 | 0.75378 | NA | 0.75588 | 0.78691 | NA | 0.86390 | 0.63746 | 0.67247 |
| k__Bacteria;p__Proteobacteria;c__Alphaproteobacteria;o__Rhizobiales;f__Xanthobacteraceae;g__Other | NA | NA | NA | NA | 0.67892 | NA | NA | NA | NA | NA | 0.74728 | NA | NA | NA | NA | 0.76445 | NA | NA | NA | 0.65238 | NA | NA | 0.55796 | NA | NA | 0.58390 | 0.64807 | NA |
| k__Bacteria;p__Proteobacteria;c__Alphaproteobacteria;o__Rhodospirillales;f__Other;g__Other | 0.84081 | 0.41514 | 0.80997 | 0.79901 | 0.46188 | 0.81256 | 0.74206 | NA | 0.48300 | NA | NA | NA | NA | 0.41233 | NA | NA | NA | NA | 0.81031 | 0.45264 | 0.82293 | 0.77039 | NA | NA | NA | NA | NA | NA |
| k__Bacteria;p__Proteobacteria;c__Alphaproteobacteria;o__Rickettsiales;f__Mitochondria;g__Other | 0.76868 | 0.28148 | 0.65879 | 0.73671 | 0.31577 | 0.74165 | 0.58471 | NA | NA | NA | NA | NA | NA | NA | NA | NA | NA | NA | NA | NA | NA | NA | NA | NA | NA | NA | NA | NA |
| k__Bacteria;p__Proteobacteria;c__Gammaproteobacteria;o__Burkholderiales;f__Sutterellaceae;g__Parasutterella | NA | NA | NA | NA | NA | 0.36509 | NA | NA | NA | NA | NA | 0.36249 | NA | NA | NA | NA | 0.28573 | NA | NA | NA | 0.75190 | NA | NA | 0.80460 | NA | 0.29333 | NA | 0.65295 |
| k__Bacteria;p__Proteobacteria;c__Gammaproteobacteria;o__Enterobacterales;f__Enterobacteriaceae;g__Escherichia-Shigella | NA | NA | 0.00979 | 0.15804 | 0.67871 | NA | NA | NA | 0.00898 | 0.16037 | 0.71463 | NA | NA | 0.00976 | 0.10104 | 0.72443 | NA | NA | 0.78546 | 0.00942 | 0.12481 | 0.08043 | 0.17663 | 0.62381 | 0.45452 | 0.49752 | 0.59580 | NA |
| k__Bacteria;p__Proteobacteria;c__Gammaproteobacteria;o__Enterobacterales;f__Morganellaceae;g__Proteus | NA | NA | NA | 0.33793 | NA | NA | NA | NA | NA | 0.37387 | NA | NA | NA | NA | 0.26133 | NA | NA | NA | 0.71225 | NA | NA | NA | 0.29276 | 0.83214 | 0.68168 | NA | NA | NA |
| k__Bacteria;p__Verrucomicrobiota;c__Verrucomicrobiae;o__Verrucomicrobiales;f__Akkermansiaceae;g__Akkermansia | 0.83947 | 0.52914 | 0.00984 | 0.00962 | 0.58526 | 0.01050 | 0.00943 | 0.64270 | 0.00898 | 0.00867 | 0.50034 | 0.00933 | 0.00858 | 0.00976 | 0.01075 | 0.70959 | 0.01044 | 0.01472 | 0.06999 | 0.00975 | 0.50707 | 0.36184 | 0.01020 | 0.41534 | 0.37635 | 0.01048 | 0.01110 | 0.91135 |

Highlighted cells Significant at P<0.05
