## Supplemental Figure 1 for "Addition of soluble fiber in low fat purified diets improves gut and metabolic health compared to traditional AIN diets"

**Supplementary Figure 1. (Study 2). Composition of fiber in the diet modulates the composition of the gut microbiota. Changes in Firmicutes/Bacteroidetes ratio in different groups dietary treatment for cecums (A) and colons (B) after 14 days on either a GBD (5001 or 5002), or high-fiber PDs (100CEL, 100IN, 100FOS, 200CEL, 200IN, 200FOS) with n = 8/group.**

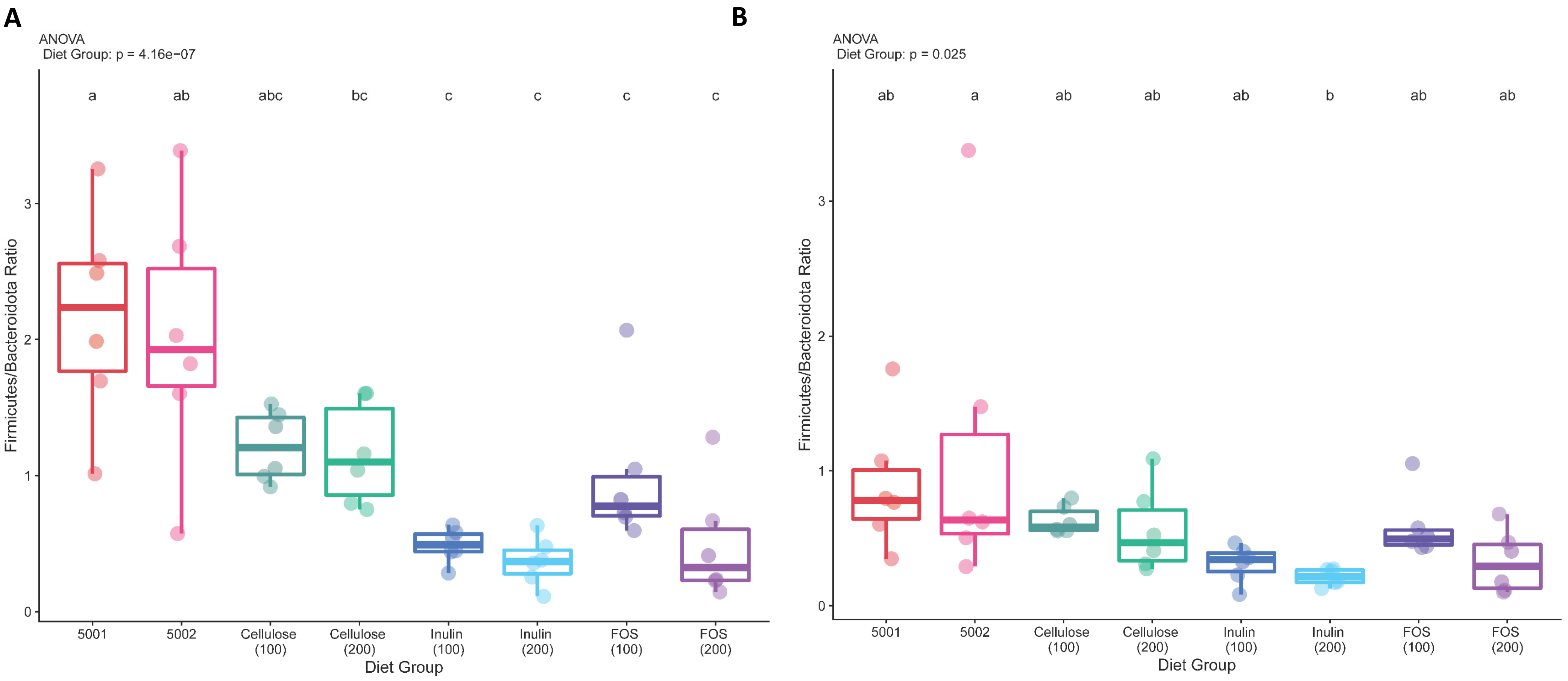
