## Supplementary figures and images for "Addition of soluble fiber in low fat purified diets improves gut and metabolic health compared to traditional AIN diets"

### Supplemental Figure 2

**Supplementary Figure 2.**

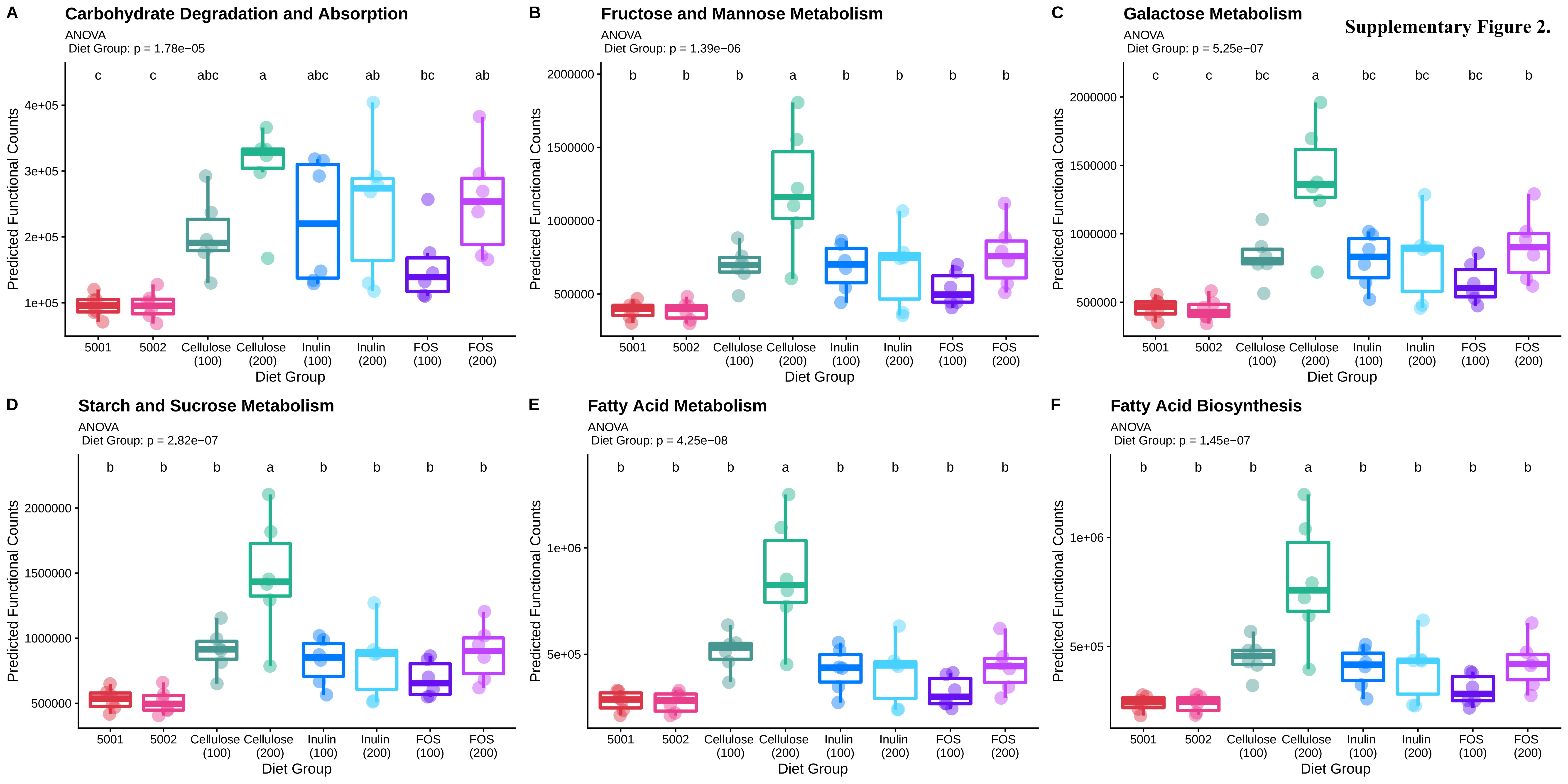
